# Causal Discovery of Feedback Networks with Functional Magnetic Resonance Imaging

**DOI:** 10.1101/245936

**Authors:** R Sanchez-Romero, J.D. Ramsey, K. Zhang, M. R. K Glymour, B Huang, C. Glymour

## Abstract

We test the adequacies of several proposed and two new statistical methods for recovering the causal structure of systems with feedback that generate noisy time series closely matching real BOLD time series. We compare: an adaptation for time series of the first correct method for recovering the structure of cyclic linear systems; multivariate Granger causal regression; the GIMME algorithm; the Ramsey et al. non-Gaussian methods; two non-Gaussian methods proposed by Hyv¨arinen and Smith; a method due to Patel, et al.; and the GlobalMIT algorithm. We introduce and also compare two new methods, the Fast Adjacency Skewness (FASK) and Two-Step, which exploit non-Gaussian features of the BOLD signal in different ways. We give theoretical justifications for the latter two algorithms. Our test models include feedback structures with and without direct feedback (2-cycles), excitatory and inhibitory feedback, models using experimentally determined structural connectivities of macaques, and empirical resting state and task data. We find that averaged over all of our simulations, including those with 2-cycles, several of these methods have a better than 80% orientation precision (i.e., the probability a directed edge is in the true generating structure given that a procedure estimates it to be so) and the two new methods also have better than 80% recall (probability of recovering an orientation in the data generating model). Recovering inhibitory direct feedback loops between two regions is especially challenging.

## 1 Introduction and Background

At rest and at work, neural physiological processes in various regions of the brain influence processes in other regions. Indirect measures of the local processes, such as the BOLD signal from fMRI (Goense & Logothetis, 2008; Kahn et al., 2011; Winder et al., 2017), result in time series whose statistical relations are often used as keys in attempts to recover causal relations among neural processes. Establishing accurate, computationally feasible statistical methods for this purpose is difficult because the measurements are of tens of thousand of non-Gaussian, sometimes non-stationary times series that are noisy indirect measures of processes for which there is little possibility of direct experimental testing and only a very limited number of relevant animal studies. The principal method of testing the accuracies of proposed statistical methods has therefore been with simulated data from biophysical models where the generating structure is of course known. The most extensive simulation study of the accuracies of statistical methods for estimating neuronal connections from BOLD signals (Smith et al., 2011) generated simulated data from Dynamic Causal Models (DCM)(Friston et al., 2003), and found that no methods tested were useful for determining directions of influence represented by the directed graphs in the data generating models. Some extant methods were not tested and several methods have since appeared that show good accuracy in most of the simulated conditions. Only two of the 28 conditions in the Smith et al., study involved feedback cycles represented by directed graphs with cycles.

Feedback structures in functional brain networks at the cellular level include both amplifying (excitatory) and control (inhibitory) connections, and it is prudent to consider the possibility of both kinds of connections at the meso-level of voxels or clusters of voxels. In addition, based on anatomical evidence, a common assumption in brain connectivity analyses is that the majority of connections between clusters of cortical voxels are bidirected (Friston et al., 2011; Zeki & Shipp, 1988; Kötter & Stephan, 2003), although this does not imply that both directions are effective in a particular scanning episode or in response to a particular stimulus. These feedback structures can be represented by cyclic directed graphs in which each brain region is represented by a vertex or node, and a directed edge, *X → Y*, represents the hypothesis that *X* is a direct (relative to the set of represented regions) cause of *Y*, that is, values of *X* and *Y* would be associated were a hypothetical experiment carried out that varied *X* while controlling all other variables other than *Y*.

Of the many methods that have been proposed for recovering causal connectivity from BOLD signals, several are inappropriate for this task. Simple correlation and total partial correlation conditioning for all the rest of the measured variables as estimated by Lasso (Tibshirani, 1996) or Glasso (Friedman et al., 2008) techniques are a priori inappropriate for causal inference because they produce undirected graphs whose edges do not even in principle capture causal connections. Simple correlations and total partial correlations give neither directions of influence nor actual connectivities even if the direction is ignored. If *X → Z → Y* or *X ← Z → Y*, then *X* and *Y* will be correlated and a graphical representation of “functional connectivity” will find an *X − Y* connection. If there are several intermediates or common causes between *X* and *Y*, their correlation coefficient can be very large even though there is no real direct causal connection between them. If, instead, *X → Z ← Y*, partial correlation of *X* and *Y* controlling for (i.e., conditioning or “partialling” on) *Z* will find an *X − Y* connection, and, again, if there are multiple variables that, like *Z*, are direct effects of *X* and *Y*, the partial correlation of *X* and *Y* can be very large. Correlation and total partial correlation thus cannot be trusted to give directions of influence or to give correct information about causal pathways between neural regions. For these reasons they will not be considered here.

Assuming independent and identically distributed (i.i.d.) data from a linear system, an asymptotically correct algorithm for estimating cyclic structures, the Cyclic Causal Discovery (CCD) procedure (Richardson, 1996), has been available for more than 20 years. Because of limitations of complexity and accuracy with finite samples, CCD has rarely been applied, but it served to show that the identification of feedback structures from i.i.d. data is theoretically possible.

Ramsey et al. (2014) presented and tested several procedures for estimating directions of influence from adjacencies of an undirected graph. These procedures exploit the non-Gaussianity of the BOLD signal. In principle, all of the non-Gaussian pairwise directionality algorithms in Ramsey et al. (2014), can detect cyclic structures when the number of variables conforming the cycle (degree of the cycle) is greater than 2, but only two of them, denoted by R1 and R4, can estimate cycles of degree 2 (indicated here as 2-cycles). Ramsey et al. (2014) tested their algorithms on simple structures of five-node ring graphs with one 2-cycle, two 2-cycles sharing a node, and two 2-cycles not sharing a node; and a more complex ten node graph with four 2-cycles. Accuracy results show that the algorithms performed better when the sample size was increased by concatenation of datasets. Nevertheless, R1 rarely detected 2-cycles while R4 had good recall and precision for detecting 2-cycles in simple structures but lost accuracy in more complex models. A search procedure for Dynamic Causal Models (Friston et al., 2011), assumes that all connections are direct feedback cycles (2-cycles) in the generating network. The procedure has been tested with good results on simulated fMRI structures with six variables. For complexity reasons in the model search strategy, the method is limited to a very small number of nodes, although it remains an open question whether more efficient modifications are possible (Freenor & Glymour, 2010). The performance of the DCM search algorithm on more complex networks with higher degree cycles and where some but not all of the connec-2 tions are direct feedback cycles has not been systematically tested. Other procedures capable of inferring cyclic structures from i.i.d., data have been published but not tested on neuronal data. Itani et al. (2010) introduced a causal search procedure based on Bayesian Networks capable of inferring cyclic structures from i.i.d., data and applied it to a protein measurements dataset. To find cyclic structures, this procedure depends on the possibility of the researcher intervening experimentally to fix the value of each variable individually. Our study focuses on non-invasive fMRI data, so we do not consider this procedure here. Similarly to Ramsey et al. (2014), Mooij et al. (2016) made a systematic analysis of pairwise directionality algorithms based on different sets of assumptions, and tested them on empirical and synthetic datasets of various kinds, but did not consider their performance with cyclic structures or neuronal data.

Most efforts to identify causal structure in neuronal systems have exploited time-lagged dependences in BOLD time series. Some of these procedures, notably those based on Granger Causality (Granger, 1969) and Group Iterative Multiple Model Estimation (GIMME) (Gates & Molenaar, 2012), are routinely used in many functional connectivity applications (Gates et al., 2014; Price et al., 2017; Zelle et al., 2017; Bellucci et al., 2017; Sreenivasan et al., 2017; Juan-Cruz et al., 2017), and their accuracies in the presence of feedbacks deserve more extensive assessment. Granger Causal methods estimate causal connections in time series by multiple regression of time-indexed variables on lagged values of variables. This autoregressive method has been supplemented by procedures for estimating “contemporaneous” causal connections by applying machine learning methods to the joint distribution of the residuals after regression (Swanson & Granger, 1997; Hoover, 2005; Spirtes et al., 2000). On the cyclic simulations of Smith et al. (2011), different versions of Granger Causality performed poorly on data from structures with 5-cycles and structures with 2-cycles. Adapting LISREL procedures (Jöreskog & Sörbom, 1993) to time series, Gates & Molenaar (2012) introduced the GIMME algorithm, that combines autoregressive and cyclic structural equations, model fit scoring, and group voting. In principle, it can accommodate cyclic structures of any degree. Using multiple data sets on the cyclic simulations of Smith et al. (2011), it achieved an almost perfect accuracy on the 5-cycle but low accuracy on the structure containing 2-cycles. The computational complexity of the search strategy limits GIMME to small (e.g., 15) numbers of regions of interest.

In this paper, using the same BOLD synthetic data simulator employed by Smith et al. (2011), the following lag-based methods are compared: a multivariate implementation of Granger Causality; the GIMME algorithm; and the Global Mutual Information Transfer (GlobalMIT) algorithm, a method based on discrete dynamic Bayesian networks (Vinh et al., 2011), together with the following i.i.d-based methods: the CCD algorithm; the Ramsey et al. (2014) non-Gaussian methods; two non-Gaussian methods proposed by Hyv¨arinen & Smith (2013); and a method due to Patel et al. (2006). We introduce and also compare two new methods, the Fast Adjacency Skeweness (FASK) algorithm and Two-Step, which exploit non-Gaussian features of the BOLD signal in different ways. Theoretical justifications is given for the latter two algorithms. The tuning parameters of all methods are optimized on separate sets of data distinct from the test data. Our test models include feedback structures with and without 2-cycles, and amplifying and control feedbacks. The best procedures are tested on four complex models derived from experimentally determined structural connectivity of macaques from Markov et al. (2013). Finally, the procedures are applied to empirical fMRI resting state data from the medial temporal lobe from 23 individuals and empirical fMRI data for a rhyming task from 9 individuals.

## 2 Methods: Data Generation

### 2.1 Simple Networks Simulations

Smith et al. (2011) fMRI BOLD data simulations are based on the DCM architecture (Friston et al., 2003), where the regions of interest are nodes embedded in a directed network, and the temporal evolution of their neuronal signals follow the linear approximation, *dz/dt* = *σ***A***z*+C*u*. Where *dz/dt* is the temporal evolution of the neuronal signals; *σ* controls the within and between nodes neuronal lag; **A** is the directed connectivity matrix defining the causal structure between nodes; *z* are the time series of the neuronal signals of the regions of interests; **C** is a matrix controlling how the external neuronal inputs *u* feed into the network. To simulate resting state data, the network is not altered by external stimuli, as it would be in task fMRI experiments, instead the *u* stimuli are modeled by an inherent Poisson process noise term for each of the regions. Finally, the observed BOLD signals are obtained by passing the neuronal signals *z* through a balloon non-linear functional model relating neuronal and vascular systems, *ỹ* = *g*(*z, θ*), where *ỹ* are the BOLD signals, *z* the neuronal signals, and is a vector of parameters describing the hemodynamics of the brain vascular system.

The standard parameters of Smith et al. (2011) simulations were kept and are briefly summarized here: Values for the coefficients of the *A* matrix were sampled from a Gaussian distribution with *mean* = 0.5, *standard deviation* = 0.1 and values truncated between 0.3 and 0.7. The diagonal of the *A* matrix was set to -1 to simulate a self-decay in each node. The input matrix *C* was modeled with an identity matrix to represent that each region of interest has one exclusive neuronal noise term. The Poisson process generating the neuronal inputs (*u* timeseries) controls two states with mean duration of 2.5 s (up) and 10 s (down). The mean neuronal lag between regions is approximately 50 ms. To simulate differences in the vascular response across the brain, the hemodynamic response function delay has variations across nodes of *standard deviation* = 0.5 s. Gaussian measurement noise with *mean* = 0 and *standard deviation* = 1 was added to the resulting BOLD signal. This implies that the *observed* BOLD signals are properly defined as *y* = *ỹ* + *e_m_*, where *ỹ* is the measurement-error-free BOLD signal and *e_m_* is the added measurement noise, in our simulations, sampled from a Gaussian distribution with *mean* = 0 and *standard deviation* = 1.

The scanning session time was set to 10 minutes, but the repetition time (TR) was reduced from 3 seconds to 1.20 seconds to reflect current acquisition protocols with higher temporal resolution, as observed in Poldrack et al. (2015). The result is an increase in datapoints of the BOLD time series from 200 to 500. As reported in Ramsey et al. (2014) the Butterworth temporal filter used in Ramsey et al. (2014) reduces the original non-Gaussianity of the BOLD data and puts at disadvantage search methods that leverage higher order moments of the distributions. For this reason, an FSL high-pass filter of 1/200 Hz was applied, which has shown a minimal impact on the non-Gaussianity of the BOLD signal (Ramsey et al., 2014).

Using this data generator model, a series of networks containing different cyclic structures with different degrees of complexity were simulated. The networks are illustrated in Figure 1 and described below:

*Networks 1, 2*, and *3* are replications of Figure 14 in Ramsey et al. (2014), with one 2-cycle, two 2-cycles sharing a node, and two 2-cycles not sharing a node.
*Network 4* is a replication of Figure 16 in Ramsey et al. (2014), a graph with ten nodes, and four 2-cycles not sharing nodes.
*Network 5* is a 2-cycle network introduced in Richardson (1996) to illustrate the CCD algorithm. Two nodes feed into a 2-cycle which in turn feeds into a single node.
*Network 6* is a chain network where two nodes feed into a 2-cycle which then branches into three nodes.
*Network 7* is a variant of Smith et al. (2011) condition 14. One node feeds into a 4-cycle which in turn feeds into a single node
*Network 8* is a second-order cyclic system illustrated in the seminal work on causal analysis by Heise (1975, p.220), useful to study the effects of interactions between signal amplification cycles (where the product of the coefficients is positive) and signal control cycles (where the product of the coefficients is negative). The network is formed by one node that feeds into a second-order cyclic system (ie., a cycle inside a cycle) with six nodes, which in turn feeds into a single node.
*Network 9* is an alternative structure to study interactions of amplifying and control cyclic structures. One node feeds into a 4-cycle that shares one node with another 4-cycle that feeds into a single node.

**Figure 1:**
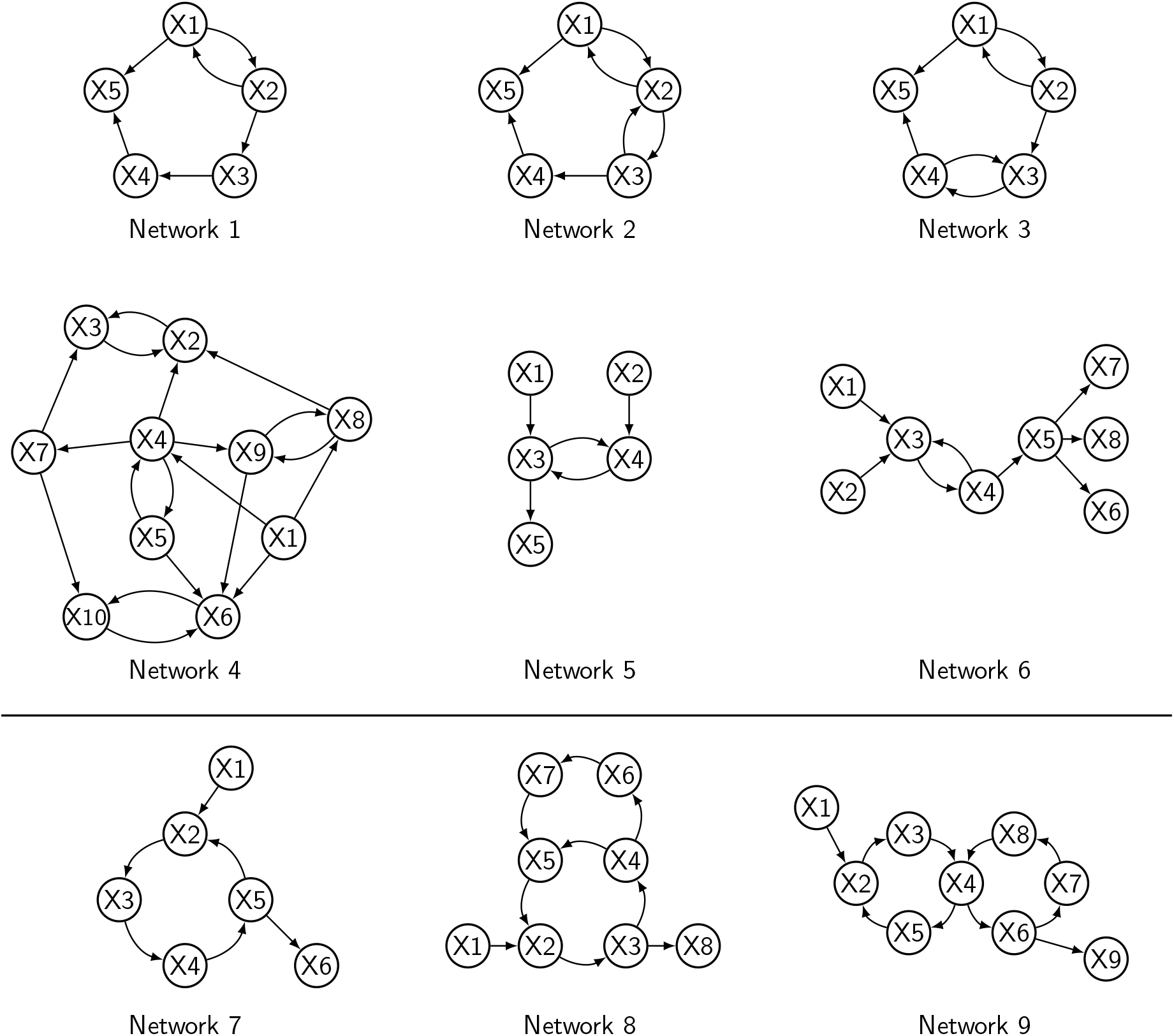
Networks containing 2-cycles: *Network 1* to *6*; and higher degree cycles: *Network 7* to *9*

Cyclic structures can amplify the signal in a system if the product of the coefficients in the cycle is positive, or control it if the product of the coefficients in the cycle is negative. Interactions (represented as node sharing) between amplification and control feedbacks can produce non-intuitive outcomes, such as unstable oscillatory signals produced by two cycles that would independently produce stable signals (Heise, 1975). Variants of the networks were made by adjusting the signs of the coefficients of the corresponding matrices to produce amplifying and control versions for *Networks 5, 6* and *7*; and combinations of amplifying and control structures for *Network 8* and *9*. In total we simulate fMRI BOLD signals for 18 cyclic networks.

For each network, 60 different individual datasets of 500 datapoints each were simulated, which can be conceived as 60 different scanning sessions. Ten of these datasets were selected at random without replacement, centered individually, and concatenated to use as a 5,000 datapoints input for the algorithms. The scans selection, centering and concatenation process was repeated 60 times, and average performance across the 60 repetitions is reported in Section 4.1.

The *centering* of individual datasets before concatenating is a necessary step to avoid spurious associations due merely to differences between the individuals means of the concatenated datasets. If datasets *D*_1_ and *D*_2_ are concatenated, and variables *X* and *Y* have zero covariance in each individual dataset, so that 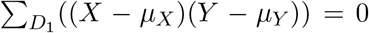 and 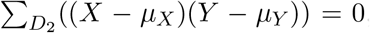, *unless* the means of the variables in each dataset are equal, *μ_X_*_(*D*1)_ = *μ_X_*_(*D*2)_ and *μ_Y_* (*D*_1_) = *μ_Y_* (*D*_2_), the resulting covariance between *X* and *Y* from the concatenated data, 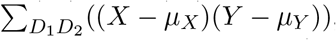, will not be zero as in the individual datasets (see Pearson et al., 1899; Ramsey et al., 2011).

### 2.2 Macaque-Based Networks Simulations

In principle, it would be straightforward to increase the complexity of the discovery problem by randomly simulating networks with more interacting excitatory and inhibitory cycles and increasing number of nodes and connections between regions of interest. A more formal way to increase the complexity and realism is to use empirical structural brain connectivity information as a blueprint. Axonal connectivity derived from tracer injection studies in non-human primates encode information about the directionality of the connections and so allow us to design simulations on which to test causal search algorithms. Recent studies in macaques have characterized axonal directed connectivity networks of cortical regions spanning the whole neo-cortex (Markov et al., 2012, 2013). Using retrograde tracer injection experiments Markov et al. (2013) build an axonal directed network of 91 cortical regions and 1,615 directed edges, with cyclic structures of different degree, including 214 2-cycles. The network is publicly available (http://core-nets.org) and is used here as a blueprint to design more complex and realistic synthetic networks.

Cyclic structures are prone to produce unstable output signals under certain conditions, for example if the product of the coefficients of the connections forming the cycle is greater than one, or if certain complex interactions are present, such as a control feedback embedded in an amplifying feedback (Heise, 1975). We have noticed in simulations that most parameterizations of the directed network of Markov et al. (2013) will produce unstable synthetic outputs. To produce stable synthetic signals, small coefficient values were often required for the coefficients of the macaque empirical directed graphs.

To simulate fMRI BOLD signals, we followed the DCM model linear parameterization as described above, except when noted. We build 4 different cases, described below, to study the performance of the algorithms under different conditions of connection density, number of nodes, complexity and number of cyclic paths, and linear coefficient strength. We test accuracies on the full Macaque network as reported by Markov et al. (2013) as well as subnetworks and pruned subnetworks. For each case we simulate 60 individual datasets.

#### Macaque SmallDegree

From the original Markov et al. (2013) macaque network we selected a subgraph from 28 target nodes where retrograde tracer injections were made, and randomly removed edges to force an average in-degree of 1.8 and out-degree of 1.8. The average in-degree and out-degree of the original macaque network is 17.7, so this considerable reduction in connectivity degree allowed us to assess the performance of the methods under sparser structures. This pruning process resulted in a subnetwork with 28 nodes, 52 edges and 10 cycles, out of which 5 are 2-cycles; the maximum cycle length is 6, and the mean cycle length is 3. All the 2-cycles are amplifying cycles. The coefficients were sampled from a Gaussian distribution with *mean* = 0.5, *std.dev* = 0.1 and values truncated between 0.3 and 0.7.

#### Macaque LongRange

The Markov et al. (2013) network reports distance in *mm* for the axonal connections. This information was used to select long range connections, defined as the connections in the top 10 percentile of the distance distribution in the network. This percentile corresponds to a threshold of 35.6*mm* (max distance 53.8*mm*). Thresholding the network using long range connections produced a subnetwork of 67 nodes, 161 directed edges, average in-degree of 2.4 and out-degree of 2.4; and 239 cycles, out of which 19 are 2-cycles, the maximum cycle length is 12, and the mean cycle length is 7. The Markov et al. (2013) network also encodes connection weights reflecting axonal density. Base connectivity coefficients were defined for the network by mapping the values of the axonal weights for the long-range connections to values in a range between 0.05 and 0.1. Then to simulate heterogeneity, for each dataset 50% of the coefficients were randomly chosen and values sampled from a Gaussian distribution with *mean* = 0.01, and *std.dev* = 0.01, were added to them.

#### Macaque LongRangeControl

This case is build as for the *Macaque LongRange* case described above, with the difference that for all the 60 datasets, the same nine 2-cycles (out of 19) were defined as control cycles by setting one of the coefficients in the cycle negative.

#### Macaque Full

We used the complete macaque axonal connectivity dense network as reported in Markov et al. (2013): 91 nodes and 1,615 edges, with average in-degree of 17.7 and out-degree of 17.7; and 214 2-cycles. As in the *Macaque LongRange* simulation, base connectivity coefficients were defined for the network by mapping the values of the axonal weights for all the connections to values in a range between 0.01 and 0.05. Then to simulate heterogeneity, for each dataset 50% of the coefficients were randomly chosen and we added values sampled from a Gaussian distribution with *mean* = 0.01, and *std.dev* = 0.01, truncating the values to 0.01 to avoid zero or negative values.

As with the previous simulations, 60 different individual datasets of 500 datapoints were generated for each of the four macaque-based networks. Ten of these datasets were selected at random without replacement, centered individually, and concatenated to use as a 5,000 datapoints input for the algorithms. The scans selection, centering and concatenation process was repeated 60 times, and average performance across the 60 repetitions is reported in Section 4.2.

For simulations with very small values for the coefficients (small size effects) as these, centering the data previous to the concatenation is particularly important. As mentioned at the end of the previous section, the concatenation of data can create spurious associations due merely to differences between the means of the individual datasets. These spurious associations may be weak, but comparable in size to real effects if the true coefficients are very small (as in our simulations). Therefore, centering is necessary to reduce the chance of false positive estimations in concatenated data.

The synthetic fMRI data described in Section 2.1 is available at http://github.com/cabal–cmu/Feedback-Discovery. The repository contains individual and concatenated datasets for each of the simple networks and Macaque-based networks, together with the corresponding graphs encoded as lists of directed edges and matrices in Matlab files.

### 2.3 Empirical Data

#### 2.3.1 Resting State Data

To assess the algorithms under real BOLD data conditions we use publicly available high-resolution 7T human resting state fMRI data from the medial temporal lobe. The fMRI data was acquired and preprocessed by Shah et al. (2017), and is publicly available at https://github.com/shahpreya/MTLnet. We summarize the data here and refer to Shah et al. (2017) for further details.

Resting state fMRI data for 23 human subjects was acquired at TR = 1 second, for 7 minutes, resulting in 421 datapoints per subject. (The original Shah et al. (2017) data contains 24 subjects, but the first one (S1) has 304 datapoints, so for consistency we exclude it from our analysis). The data was band-pass filter in the range 0.008–0.08 Hz, and no spatial smoothing was applied to avoid mixing of signals between neighboring regions (Shah et al., 2017). We consider the following seven regions of interest from the medial temporal lobe in each hemisphere: perirhinal cortex divided into Brodmann areas 36 and 35 (BA35 and BA36); parahippocampal cortex (PHC); entorhinal cortex (ERC); subiculum (SUB); Cornu Ammonis 1 (CA1); and a region comprising CA2, CA3 and dentate gyrus together (CA23DG); averaging the individual signals of these last three areas into one regional signal helps to reduce potential problems in connectivity estimation arising from the mixing of signals of neighboring areas (Smith et al., 2011), which can be especially acute in regions that are challenging to parcel out, such as CA2, CA3 and the dentate gyrus (Ekstrom et al., 2009; Preston et al., 2010).

FASK and Two-Step(Alasso) were run on 23 repetitions of ten individual subjects concatenated (4,210 datapoints), for the seven medial temporal lobe ROIs of the left and right hemispheres separately. The individual data was standardized (centered and variance normalized to one) before being concatenated. For comparison, we ran both algorithms on the 23 subjects datasets individually (421 datapoints). Each individual dataset was previously standardized. Results are reported in Section 5.

#### 2.3.2 Task Data

To test the performance of the algorithms with fMRI task data, we use data previously published in Ramsey et al. (2010), in which 9 subjects judged if a pair of visual stimuli rhymed or not. In each 20 seconds block, 8 pairs of words were presented for 2.5 seconds each. Four blocks of words were followed by four blocks of pseudowords. Task blocks were separated by fixation blocks of 20 seconds. Data was acquired with a 3T scanner, with TR = 2 seconds, resulting in 160 datapoints. Raw data is available at the OpenfMRI Project (https://openfmri.org/dataset/ds000003/); and the preprocessed data used here is available at https://github.com/cabal-cmu/Feedback-Discovery. We refer the reader to Ramsey et al. (2010), for more details about the acquisition and preprocessing of the data, and region of interest definition.

For our analysis we considered eight regions of interest: left and right occipital cortex (LOCC, ROCC); left and right anterior cingulate cortex (LACC, RACC); left and right inferior frontal gyrus (LIFG, RIFG); left and right inferior parietal (LIPL, RIPL). In addition, we included an Input variable build by convolving the rhyming task boxcar model with a canonical hemodynamic response function. If the algorithms infer orientations correctly then edges from the Input variable must feedforward into the regions of interest, and no edge should point backwards into the Input variable. This is a reliable first test that can be used to evaluate the performance of causal search algorithms on task data as shown in Ramsey et al. (2010, 2014) and Sanchez-Romero (2012).

Given the small number of subjects (9) and reduced sample size (160 datapoints), FASK and Two-Step were run on 1 repetition of 9 individual subjects concatenated (1,440 datapoints) for the Input variable and the eight regions of interest. As with the resting state data, the individual data was standardized before being concatenated. For comparison, we ran both algorithms on each of the nine subjects individually (160 datapoints). The datasets were standardized before running the algorithms. Results are reported in Section 5.

## 3 Methods: Cyclic Search Procedures

Previously published lag-based methods and methods that assume i.i.d., data are tested here. Their principal assumptions are mentioned and we refer the reader to the original papers for specific details. Two novel algorithms are presented that can discover cyclic structures: Fast Adjacency Skewness (FASK) algorithm and Two-Step.

Some of the procedures tested here are purely orientation algorithms and require as initial input a list of adjacencies (undirected edges); while others estimate a list of adjacencies as a first step followed by orientation rules. For the procedures that require it, we use the Fast Adjacency Search stable (FAS-stable) algorithm as an adjacency estimator. The FAS-stable algorithm is the order independent adjacency search of the PC-stable algorithm that avoids spurious connections between parents of variables (Spirtes et al., 2000; Colombo & Maathuis, 2014). FAS-stable builds an undirected graph by iteratively testing conditional independence facts under increasing size of the conditioning set. We use a Bayesian Information Criterion (BIC) (Schwarz et al., 1978) approach to perform the conditional independence tests needed by the algorithm, in the following way: For two variables *X, Y* and a set **S** of adjacent variables of *X* or *Y*, if the BIC score for the linear model *X* ← S is better than the BIC score for the increased model *X* ← {*Y* ∪ **S**}, we conclude that *X* is independent of *Y* conditional on **S** (*X* ╨ *Y*|**S**). Using the BIC score in this way we can ask if the inclusion of *Y* in the original model covariates (**S**) increases the explanatory power of the model. If this is not the case, we have evidence that *X* and *Y* are not dependent when we take into account the effect of the conditioning set **S**. The BIC score as used by FAS-stable has an additional user-input penalty discount term *c*, used to force extra sparsity on the estimated model if necessary. We defined the score here as: BIC^*^ = −2ln(*ML*) + *ck*ln(*n*), where *ML* is the maximum likelihood of the model, *c* is the extra penalty discount, *k* is the number of covariates, and *n* is the sample size. If the penalty discount *c* is set to 1, we end up with the original BIC score. We used the FAS-stable Java implementation from the Tetrad suite of algorithms (www.phil.cmu.edo/tetrad/ and www.ccd.pitt.edu/tools/), and pseudo-code for FAS-stable is included in Supplementary Material A.

Each search procedure has one or more “tuning parameters.” As described in Supplementary Material B (Section B1), the optimal tuning parameters were estimated using a subset of graphical models with a different set of model coefficients than those for the test data. Trade-offs between precision and recall in optimizing search parameters were decided using Matthews correlation (Matthews, 1975). The resulting optimal parameters often require very small values for independence tests.

### 3.1 Procedures Assuming Temporal Data

We run a multivariate implementation of Granger Causality, MVGC, by Barnett & Seth (2014) (Matlab code at www.sussex.ac.uk/sackler/mvgc/). MVGC assumes the structure is linear, Gaussian, and a stationary time series. For each pair of variables *X* and *Y* in the set, MVGC computes the Granger causal multiple regressions conditioning on the rest of the variables in the set, and decides if *X → Y*, *X ← Y* or *X* ⇄ *Y*. The order of the temporal lag for the regressions is chosen via a BIC score comparison across increasing lag models. For our BOLD simulated data the lag order chosen is always one or two, which is expected given the sampling resolution of fMRI data. MVGC as implemented by Barnett & Seth (2014), has a parameter α, that controls the false discovery rate (FDR) significance threshold (Benjamini & Hochberg, 1995), for the multiple Granger’s F-tests. Our parameter tuning process (described in Supplementary Section B1) set a value of *α* = 10^−5^. We show test data results for this parameter value.

For the GIMME algorithm (Gates & Molenaar, 2012) we used the authors’ R implementation (http://CRAN.R-project.org/package=gimme). GIMME assumes linear structure and Gaussian time series data. It combines autoregressive and non-recursive structural equations, model fit scoring and group voting to output a group graph, and then extends the graph separately for each individual dataset. Since in our simulations, for each graph and parameterizations all simulated scans are alike except for chance variation, in comparing GIMME with other algorithms we use the group graph. GIMME, as implemented in R, has a single parameter for the voting step that sets the minimum frequency of appearance for an edge across the individual graphs to be included in the group graph. The tuning parameter process found two optimal values of 50% and 60%. We show test data results for the algorithm using both values, and indicate them as GIMME-50 and GIMME-60 respectively.

GlobalMIT (Vinh et al., 2011) assumes the data comes from a temporal causal process that can be modeled as a dynamic Bayesian network, allowing for self-loops and contemporaneous relations. The procedure assumes linear systems and discrete multinomial data. We use the authors’ Matlab implementation (http://code.google.com/archive/p/globalmit/). GlobalMIT searches for each variable *X*, the set of parents across all the rest of the variables, **V** \ {*X*}, that maximize a mutual information target score. This search strategy is inefficient since with increasing number of variables, the number of combinations of possible sets of parents that have to be tested grows exponentially. In simulations, we have seen that even with ten variables GlobalMIT can take considerable time to return. To overcome this computational problem, FAS-stable with a penalty discount of *c* = 2, was used as a preprocessor to obtain for each variable, *X*, a set of adjacent variables *adj*(*X*). We restrict the GlobalMIT parent search to *adj*(*X*) instead of **V** \ {*X*}. This considerably reduces the running time of the procedure, since in general we expect |*adj*(*X*)| < |**V** \ {*X*}|. In our simulated structures GlobalMIT with FAS-stable preprocessing is 50 times faster than regular GlobalMIT. GlobalMIT is considered here because it has recently been applied to fMRI data from the human hippocampus recovering a well known cyclic structure between the cornu ammonis, the dentate gyrus and the entorhinal cortex (P. Santos et al., 2017). For our simulated continuous data, the best training results were obtained when the data was discretized into three values, the presence of self-loops was assumed (consistent with the DCM model), and an α value controlling the significance level for the mutual information test for independence was set to α = 10^−16^. We show test data performance results for these parameters and refer to the procedure as GlobalMIT(FAS).

### 3.2 Procedures Assuming i.i.d., Data

Theoretically, the temporal separation between recorded BOLD observations within an fMRI scan, usually between 1 and 3 seconds depending on the scanning protocol, should make most pairs of records from different sampling times practically independent. This independence has been shown by Dubois et al. (2017) using MyConnectome resting state data (Poldrack et al., 2015) (available at http://myconnectome.org/wp/data-sharing/). The BOLD fMRI recordings can be well approximated as linear temporal aggregations of neural states occurring at a faster rate (Boynton et al., 2012). Under this assumption, Gong et al. (2017) showed that the linear instantaneous causal relations between the aggregated time series correspond to the true, time-delayed causal relations in the original causal processes if the aggregation window is large compared to the underlying causal events.

The proof of large sample correctness of the CCD algorithm (Richardson, 1996) assumes linearity and i.i.d., data. In principle it can find cyclic structures from conditional independence relations. We used an optimized version of CCD, called CCD-max, that improves performance in simulation by enhancing the orientation accuracy of unshielded colliders (paths of the form *X → Z ← Y*, with *X* not adjacent to *Y*). This is done by finding the conditioning set S that confirms the conditional independence of *X* and *Y* given **S** with the best BIC^*^ score, and orienting a collider *X → Z ← Y* if *Z* is not in the optimal conditioning set S. We use the Java implementation in the Tetrad suite of algorithms (www.phil.cmu.edu/tetrad/). CCD-max uses as parameter a penalty discount *c* for the BIC^*^ score for the conditional independence decisions required by the algorithm. The penalty discount was set to *c* = 2 according to the results from the parameter tuning procedure. We show test data results for this value.

The Ramsey et al. (2014) algorithms, R1, R2 and R3 are orientation methods that assume i.i.d., non-Gaussian data. R2 and R3 assume acyclicity of the model. R1 can in principle find 2-cycles and higher degree cycles. They require as input a set of adjacencies from an undirected graph. Thus, we complemented them with the output of the FAS-stable adjacency search. R1, R2 and R3 do not require any parameter, but FAS-stable requires a penalty discount for the BIC^*^ score for the conditional independence tests. Following the parameter tuning results, the penalty discount was set to *c* = 2 for all the methods that use FAS-stable as a first step for adjacency search. These methods are implemented in the Java Tetrad suite of algorithms (www.phil.cmu.edu/tetrad/ and www.ccd.pitt.edu/tools/) and source code is available at http://github.com/cmu-phil/tetrad. We refer to them in the results section as FAS+R1, FAS+R2 and FAS+R3.

Skew and RSkew from Hyv¨arinen & Smith (2013), and Patel from Patel et al. (2006), as implemented by Smith et al. (2011), are pairwise orientation methods that assume i.i.d., non-Gaussian data. As with the Ramsey et al. (2014) methods, these algorithms require as input a set of adjacencies, so we complemented them with the output of the FAS-stable algorithm with a penalty discount of *c* = 2. Skew, Rskew and Patel do not require any parameter. These orientation methods are also implemented in the Tetrad suite. We refer to them in the results section as FAS+Skew, FAS+RSkew and FAS+Patel.

#### 3.2.1 The FASK Algorithm

The idea of the FASK algorithm is as follows: First, FAS-stable is run on the data, producing an undirected graph. We use the BIC^*^ score as a *conditional independence test* with a specified penalty discount *c*. This yields undirected graph *G*_0_. The reason FAS-stable works for sparse cyclic models where the linear coefficients are all less than 1 is that correlations induced by long cyclic paths are statistically judged as zero, since they are products of multiple coefficients less than 1. Then, each of the *X − Y* adjacencies in *G*_0_ is oriented as a 2-cycle *X* ⇄ *Y*, or *X → Y*, or *X ← Y*. Taking up each adjacency in turn, one tests to see whether the adjacency is a 2-cycle by testing if the difference between *corr*(*X, Y*) and *corr*(*X, Y* | *X* > 0), and *corr*(*X, Y*) and *corr*(*X, Y* |*Y* > 0), are both significantly not zero. If so, the edges *X → Y* and *X ← Y* are added to the output graph *G*_1_. If not, the Left-Right orientation is rule is applied: Orient *X → Y* in *G*_1_, if (*E*(*XY* |*X* > 0)*/*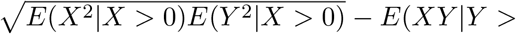 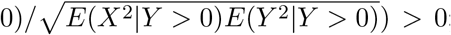; otherwise orient *X ← Y*. *G*_1_ will be a fully oriented graph. For some models, where the true coefficients of a 2-cycle between *X* and *Y* are more or less equal in magnitude but opposite in sign, FAS-stable may fail to detect an edge between *X* and *Y* when in fact a 2-cycle exists. In this case, we check explicitly whether *corr*(*X, Y* |*X* > 0) and *corr*(*X, Y* |*Y* > 0) differ by more than a set amount of 0.3. If so, the adjacency is added to the graph and oriented using the aforementioned rules.

Justification and pseudo-code for FASK are in Supplementary Material A. Java source code for FASK is at http://github.com/cmu-phil/tetrad. FASK uses two parameters, a penalty discount *c* for the BIC^*^ score in the FAS-stable adjacency search and an α threshold for the test of difference of correlations in the 2-cycle detection step. Following results from the parameter tuning procedure, we set the penalty discount at *c* = 2 and α = 10^−6^ for the test data runs.

#### 3.2.2 The Two-Step Algorithm

The Two-Step algorithm represents a linear causal structure, with possible unmeasured confounding variables and cyclic relationships, as **X = BX + MC + E**. Where **X** is a vector of measured variables, **B** is the directed connectivity matrix defining the causal structure between measured variables; **M** is a vector indicating which observed variables are affected by unmeasured confounders; **C** is a vector of unmeasured confounders, if they exist; and **E** is the vector of mutually independent noise terms. Two-Step applies the principles of independent component analysis (ICA) (Hyvarinen, 1999) and to do so it expresses the observed variables **X** as a mixture of a particular set of unmeasured components: **X = (I - B)^-1^(MC + E)**. **I** is the identity matrix; (**I-B**)^-1^ defines the mixing matrix; and **(MC + E)** defines the unmeasured components. Generally speaking, the previous equation does not follow the ICA model since the components defined by **(MC + E)** are not necessarily mutually independent because of the possible presence of unmeasured confounders in **C**. Properly, the equation corresponds to the independent subspace analysis (ISA) model (Theis, 2007; Gruber et al., 2009): The components in **(MC + E)** can be divided into mutually independent variables or groups of variables, where the variables within the same group are not independent. Each of these groups is a set of confounders plus noise terms that influence one another or are influenced by some common unknown mechanism. Under mild assumptions, the solution to ISA can be found by applying ICA and then testing for the independence between the ICA outputs. Otherwise, if there are no confounders, the equation is a standard ICA model.

The above equation can be expressed as (**I - B**)**X = Y**, where **Y = (MC + E)**. Now, independent components methods aim to find a suitable matrix **B** so that the components contained in **Y** are as independent as possible. Without constraints on **B**, we may end up with local optimal solutions, for which **B** is not necessarily the real directed connectivity matrix, or with considerable random error in the estimated **B** matrix on finite samples. To solve this problem in a computationally efficient and reliable way, the Two-Step algorithm estimates the directed connectivity matrix **B** in a divide-and-conquer manner. In a first step the algorithm infers the undirected adjacencies between the observed variables in **X**, and makes use of these adjacencies to reduce the free-parameters in **B**. In a second step, the algorithm tunes the nonzero entries of **B** to make the components in **Y** as independent as possible, with a sparsity constraint on **B**.

In particular, in the Two-Step algorithm, the first step finds the undirected adjacencies over the nodes **X**. If two nodes, *X_i_* and *X_j_*, are not adjacent, then the entries *B_ij_* and *B_ji_* are constrained to zero in the **B** connectivity matrix. We test two different alternatives for Step 1. The first learns the undirected adjacencies using FAS-stable. The FAS-stable algorithm outputs a set of undirected adjacencies among the variables **X**. If two variables are not adjacent in the set, then the corresponding entries in the **B** matrix are constrained to zero. The second alternative learns the undirected adjacencies via adaptative Lasso (ALasso) (Zou, 2006), a regularized regression method with adapted penalization for each individual coefficient. Each node *X_i_* is regressed on the rest of the nodes in **X** using ALasso. For two variables, *X_i_* and *X_j_*, if the value of the ALasso regression coefficient *b_ij_* shrinks to zero, then the *B_ij_* entry in the **B** connectivity matrix is constrained to zero. All the non-zero entries in **B** will be considered as free parameters for the estimation of **B** in the second step of the algorithm. Step 2 estimates the free parameters in the directed connectivity matrix **B** that maximize the independence of the components of (**I - B**)**X = Y**, also with a sparsity constraint on all remaining free parameters of **B**. The current implementation uses ICA with sparse connection matrices (Zhang et al., 2009). Because of the sparsity constraint in Step 2, the **B** matrix entries are expected to be as small as possible. As a consequence of this constraint and initializing the free parameters for the estimation of *B* with small values, we avoid the permutation issue present in ICA-LiNGAM (Shimizu et al., 2006), and in the presence of a cyclic structure the method tends to find the most stable solution for which the connectivity coefficients in the cycles are small. In addition, the **B** matrix final values can be thresholded to censor values close to zero that were not penalized in the previous steps.

We used a Matlab implementation of the Two-Step algorithm available at http://github.com/cabal-cmu/Two-Step/. From the parameter tuning process described in Supplementary Material B (Section B2), for Two-Step using FAS-stable we set a penalty discount of *c* = 2 for the FAS-stable step; a sparsity penalty of *log*(*N*)*λ* for the **B** matrix estimation, where *N* is the sample size and λ was set to 64; and the absolute values of the final **B** matrix were thresholded at 0.15. We refer to the procedure as Two-Step(FAS). For Two-Step using Alasso, a penalization term equal to *log*(*N*)*/*2 was used for the Alasso regressions; the sparsity penalty for the **B** matrix estimation was set to *log*(*N*)32; and the final absolute values of the **B** matrix were thresholded at 0.15. We refer to the method as Two-Step(Alasso).

As noted above, the search procedures require at least one user-input parameter; some require two or three. In order to give fair comparisons under parameters optimal for the respective algorithms and kind of data, datasets were created exactly as the ones described in Section 2.1, but changing the coefficient values range, by sampling them from a Gaussian distribution with *mean* = 0.4, *std.dev* = 0.1, and truncated between 0.2 and 0.6. We refer to these data as *training* datasets. Full description of the parameter tuning process for all algorithms tested and simple and macaque-based networks is included in Supplementary Material B.

## 4 Simulations Results

### 4.1 Simple Networks Results

Each algorithm is parameterized as indicated in Section 3 and its performance measures are averaged across the 18 simulated networks of Section 2.1. The performance of the algorithms was measured using *Precision* = *True Positives/*(*True Positives* + *False Positives*) and *Recall* = *True Positives/*(*True Positives* + *False Negatives*), for adjacencies, orientations and 2-cycle detection. Precision and recall range from 0 to 1. If both precision and recall are equal to 1, this indicates a perfect recovery of the true graph with no extra edges (false positives) and no missing edges (false negatives). Running time in seconds for each algorithm is also reported. All the runs were executed on a MacBook Pro, 2.8 GHz Intel Core i7 processor, 16GB of memory, with macOS Sierra 10.12.6.

The results are presented in two parts. First for the structures with 2-cycles (*Network 1-6*) and then for the structures with higher degree cycles but not 2-cycles (*Network 7-9*).

Figure 2 shows precision and recall averaged across the ten simulations that contain 2-cycles: *Networks 1* to *6* with amplifying and control variants (see Supplementary Material C, Section C1, for details). For each simulation, precision and recall results were averaged across 60 repetitions of ten concatenated datasets.

**Figure 2:**
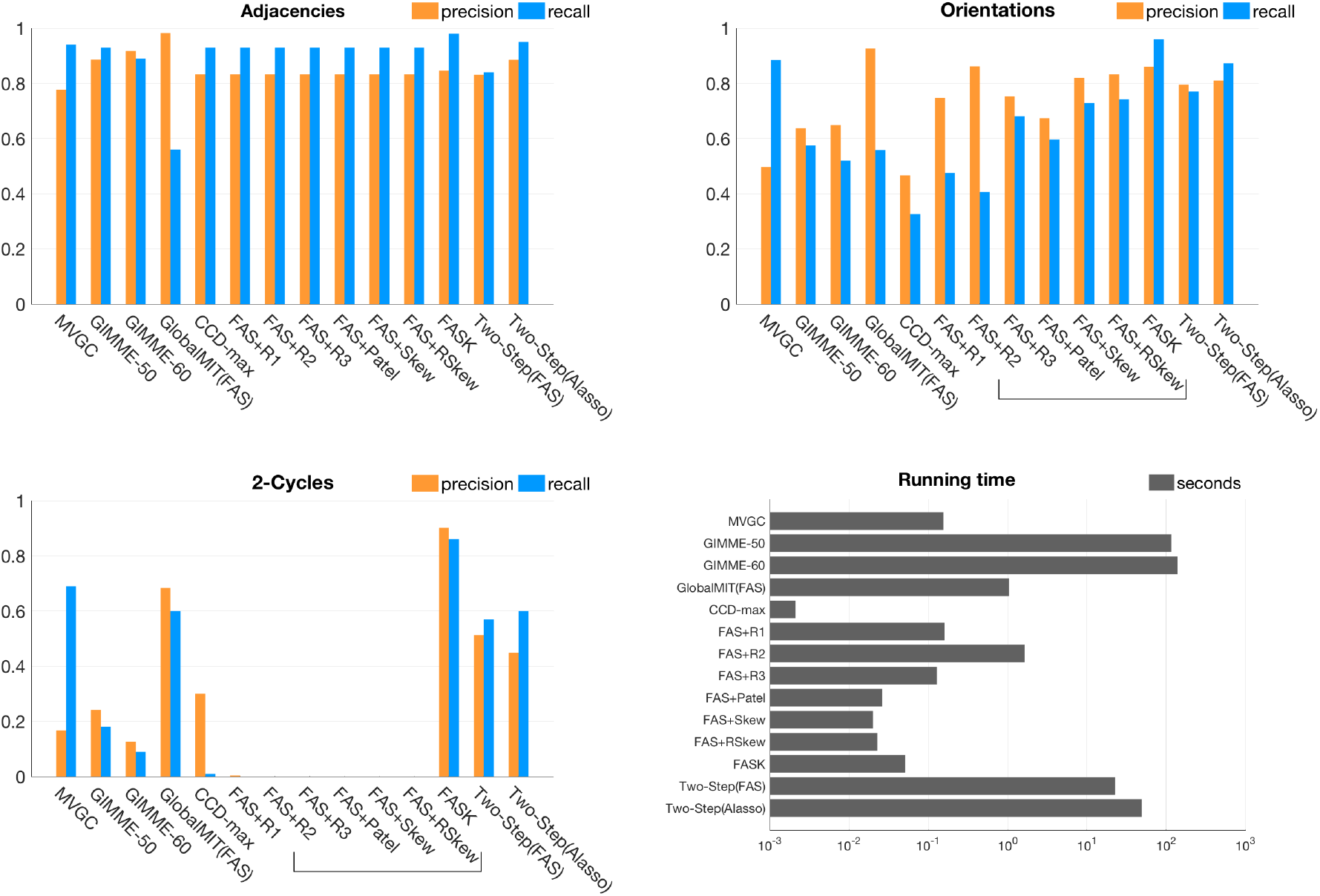
Precision and recall for adjacencies, orientations and 2-cycles for each algorithm averaged across the ten structures in Figure 1 containing 2-cycles: *Network 1* to *Network 6* with amplifying and control variants. Algorithms that cannot detect 2-cycles by design are inside a bracket: R2, R3, Patel, Skew and RSkew.

Figure 3 presents average precision and recall across eight simulations containing higher-degree cycles but *not* 2-cycles: *Network 7* to *9* with their amplifying and control variants (see Supplementary Material C, Section C2, for details). For each simulation, precision and recall results were averaged across 60 repetitions of ten datasets concatenated. In these results there is no recall for 2-cycles given that the true structures do not have 2-cycles. Instead, the average number of 2-cycle false positives is plotted to show algorithms that are prone to false detections. A bracket is drawn around those methods that by design cannot detect 2-cycles: R2, R3, Patel, Skew and RSkew.

Tables with complete results for average precisions, recalls and running times for each of the tested algorithms under each of the 18 simulated networks for 60 repetitions of ten test datasets concatenated are included in Supplementary Material C (Sections C1 and C2).

**Figure 3:**
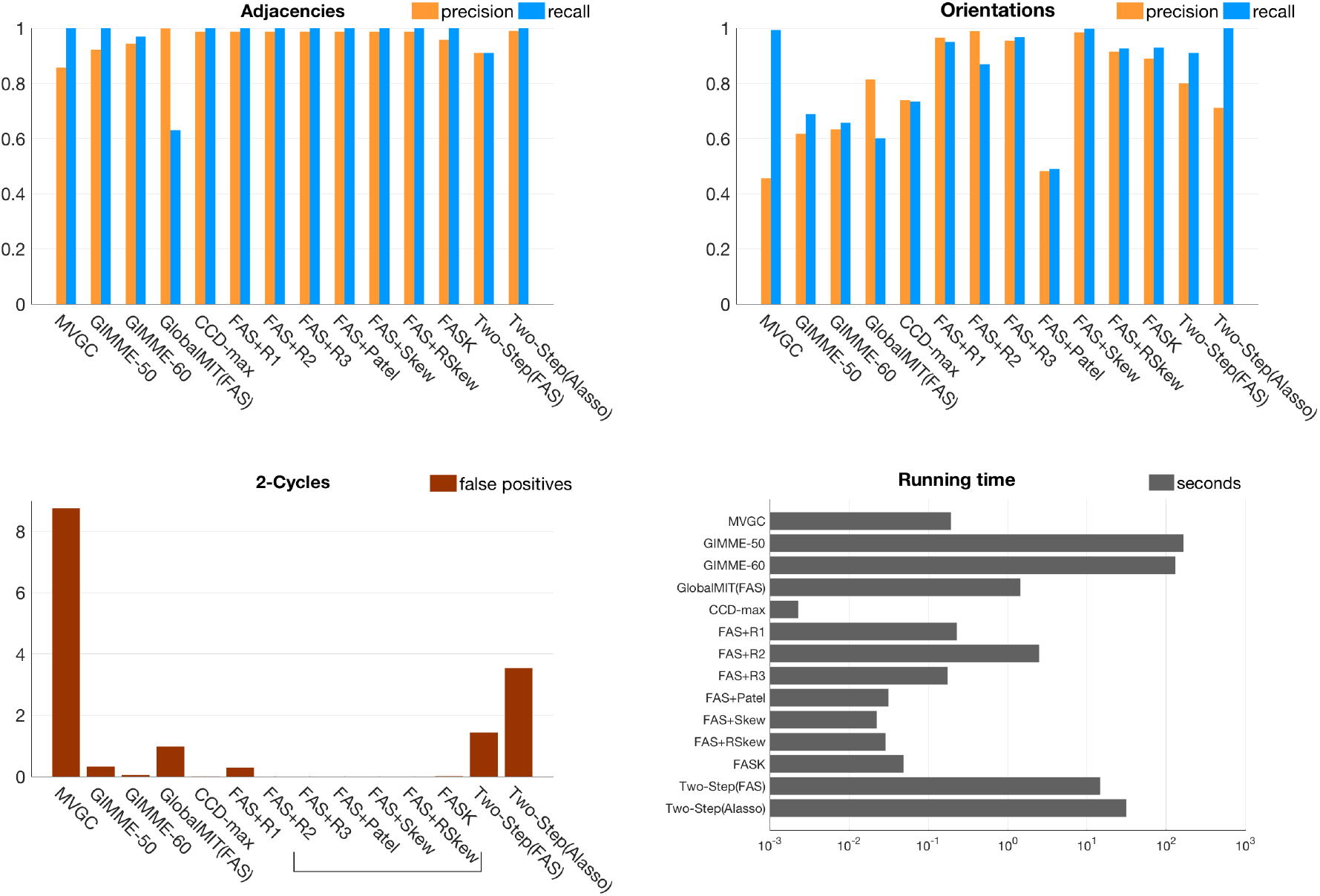
Precision and recall for adjacencies and orientations; number of 2-cycle false positives, and running times for each algorithm averaged across eight structures containing higher degree cycles but not 2-cycles: *Network 7* to *Network 9* and their amplifying and control variants. Algorithms that cannot detect 2-cycles by design are inside a bracket. Note that false positives for 2-cycles are the average number of false positives over the 60 repetitions.

In simple networks containing 2-cycles and higher degree cycles (Figures 2 and 3), all methods have in average high precision and recall for adjacencies, with the exception of GlobalMIT(FAS) that has high precision but very low recall, meaning it fails to detect true adjacencies. The GlobalMIT lag-based procedure may eliminate true edges when searching for the parent set for each variable. In terms of orientations, FASK and Two-Step(Alasso) have in average the best combination of precision and recall. Of the lag-based methods, MVGC has a good recall but low precision due to incorrectly judging many adjacencies as 2-cycles. GlobalMIT(FAS) has low orientation recall due to the low adjacency recall, but good precision. GIMME fails in both orientation precision and recall. Of the methods that can detect 2-cycles, FASK has by far the best 2-cycle precision and recall, followed by GlobalMIT(FAS). Both implementations of Two-Step have similar 2-cycle recall and precision, and MVGC loses in precision by producing a lot of false positive 2-cycles. Figure 3 shows that MVGC is by far the algorithm with the largest average number of 2-cycle false positives.

GIMME is the slowest method of all, with running times that can reach 13 minutes for 10 variable problems. This limitation of GIMME is probably caused by an inefficient model search strategy adapted from LISREL. Two-Step(Alasso) is also relatively slower and can take up to 2 minutes in 10 variable problems. Two-Step speeds-up if FAS-stable is used as the first step, which indicates that FAS-stable is a faster algorithm to infer adjacencies than Alasso if the true graph is sparse. The rest of the algorithms are very fast and take milliseconds to run in lower dimension networks such as these. These running times have to be considered taking into account the machine used (MacBook Pro, 2.8 GHz Intel Core i7 processor, 16GB memory, with macOS Sierra 10.12.6) and the various softwares used to implement the different algorithms: Java, Matlab and R.

### 4.2 Macaque-Based Networks Results

FASK and Two-Step(Alasso) were the two algorithms that achieve the best combination of precision and recall for adjacencies and orientations under the simple cyclic networks of Figure 1. So, their performance was analyzed under more complex networks based on macaque structural connectivity.

Both algorithms were run on 60 repetitions of ten concatenated datasets sampled without replacement from 50 individual test datasets. Average precision and recall for adjacencies, orientations and 2-cycles, and running times in seconds, are reported in Table 1. Corresponding standard deviations are in Supplementary Material C (Section C3).

**Table 1:**
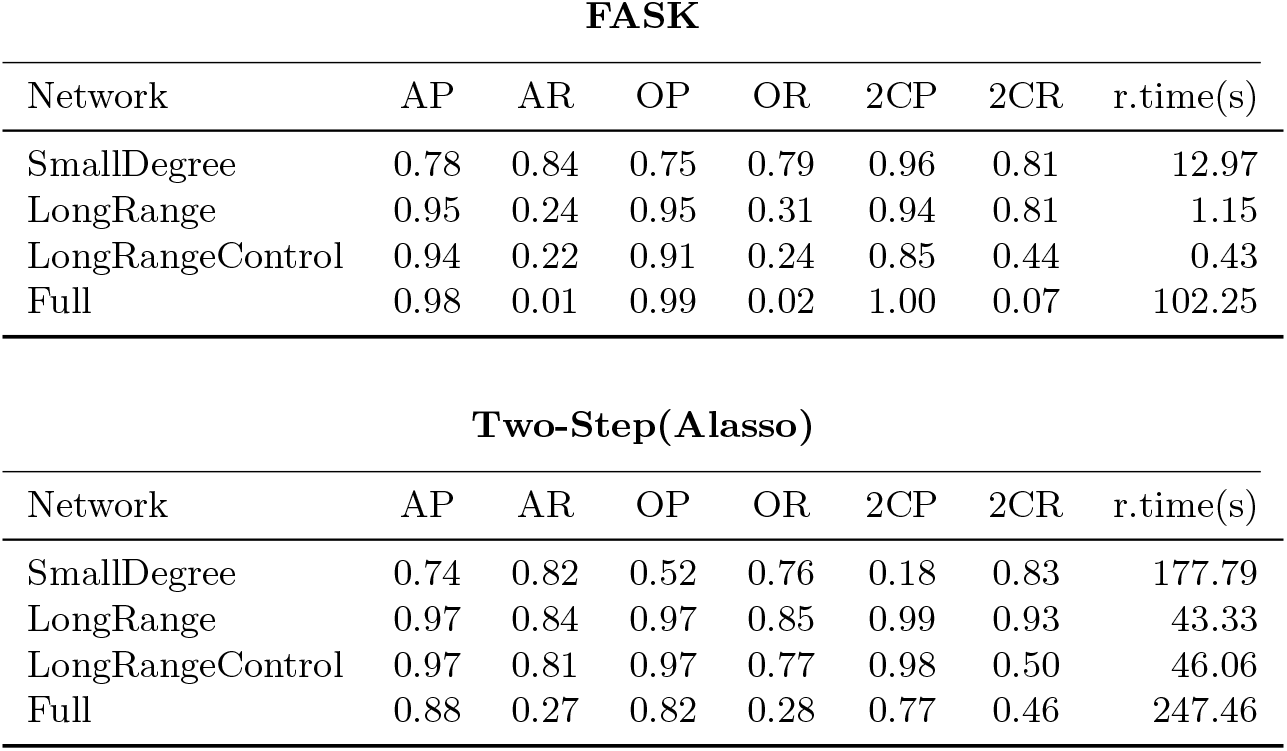
Average over 60 repetitions of ten concatenated datasets. Results for FASK and TwoStep(Alasso) for the four different macaque-based networks described in Section 2.2. Precision and recall for adjacencies (AP, AR), orientations (OP, OR), 2-cycles (2CP, 2CR), and running times in seconds (r.time).

We choose running parameters for the algorithms according to the procedure described in Supplementary Material **B** (Section B2). FASK was run with a penalty discount of *c* = 1 and α = 10^−7^ for the *SmallDegree* network; with penalty discount *c* = 1 and α = 10^−^1 for *LongRange* and *LongRange Control*; and with penalty discount *c* = 2 and α = 10^−1^ for the *Full* network. Two-Step with Alasso was run with λ = 2 and **B** threshold = 0.10 for the *SmallDegree* network; λ = 2 and **B** threshold = 0.05 for *LongRange* and *LongRange Control*; and α = 10 and **B** threshold = 0.005 for the *Full* network.

FASK showed excellent adjacency precision but unusably low recall in the *LongRange* and *Full* networks, probably due to the small valued coefficients used to guarantee cyclic stability in these simulations. As the coefficients values get really small the BIC^*^ score used by the FAS-stable adjacency search loses recall but keeps its high precision. Two-Step(Alasso) achieves a precision comparable to FASK, but with a superior performance in recall. This result indicates that Alasso may be a better method than FAS-stable to detect adjacencies when coefficients are very small and the sample size is large enough.

The orientation recall for both algorithms is low due to the low adjacency recall, but their precision is excellent considering the complexity of these problems, with the exception of the *SmallDegree* case for which the orientation precision is considerably lower. Two quantitative measures of the non-Gaussianity of the data, the Anderson-Darling score and the skewness, show that the simulated data for the *SmallDegree* macaque network is more Gaussian relative to the other simulated macaque networks. This we think is the reason why both algorithms do not perform as well in orientation precision as with more non-Gaussian data.

Both FASK and Two-Step(Alasso) show high precision with lower recall for 2-cycle detection, with the exception of the *LongRangeControl* macaque network. For the *LongRangeControl* simulation, 9 out of 19 2-cycles were set as control (inhibitory) 2-cycles, with one coefficient positive and the other negative. Control 2-cycles are challenging because the interaction of the positive and negative coefficients in the cycle can produce data that make the cycle variables look independent. None of the seven tested algorithms capable of inferring 2-cycles were good at detecting control 2-cycles in our simulations (See Tables C6 and C5 in Supplementary Material C for individual results for networks with control 2-cycles).

Consistent with the running times of the previous simple networks, FASK is considerably faster than Two-Step(Alasso), sometimes by an order of magnitude. For the dense network of 91 nodes and 1,615 edges (*Full* macaque network), FASK took 1.7 minutes, and TwoStep(Alasso) 4.1 minutes in average to return. These are excellent running times considering the complexity of the structure.

One way to improve the recall of FAS-stable is to increase the sample size. We explore sample size effects by concatenating increasing numbers of datasets for the *LongRange* network and present results for FASK in Table 2. Results are for one repetition of 10, 20, 30 and 40 subjects concatenated. For comparison we also include results for Two-Step(Alasso). Both algorithms were run with the aforementioned parameters. Table 2 shows that indeed, FASK adjacency and orientation recall can be improved by increasing the sample size, without detriment for the precision. Sample size increase improvements also work for Two-Step, but in lesser degree.

**Table 2:** Results for FASK and Two-Step with Alasso for increasing sample size under *LongRange* simulation, which has very small coefficients for the linear interactions. Results for one repetition for different numbers of concatenated test datasets. Precision and recall for adjacencies (AP, AR), orientations (OP, OR), 2-cycles (2CP, 2CR), and running times in seconds (r.time).

| FASK |  |  |  |  |  |  |  |
| --- | --- | --- | --- | --- | --- | --- | --- |
| sample size | AP | AR | OP | OR | 2CP | 2CR | r.time (s) |
| 5,000 (10 datasets) | 0.97 | 0.26 | 0.98 | 0.31 | 1 | 0.68 | 3.33 |
| 10,000 (20 datasets) | 1 | 0.40 | 0.99 | 0.47 | 0.95 | 0.95 | 17.53 |
| 15,000 (30 datasets) | 1 | 0.51 | 0.99 | 0.57 | 0.95 | 1 | 162.03 |
| 20,000 (40 datasets) | 1 | 0.58 | 0.97 | 0.63 | 0.86 | 1 | 1354.78 |

| Two-Step(Alasso) |  |  |  |  |  |  |  |
| --- | --- | --- | --- | --- | --- | --- | --- |
| sample size | AP | AR | OP | OR | 2CP | 2CR | r.time (s) |
| 5,000 (10 datasets) | 0.97 | 0.85 | 0.97 | 0.86 | 1 | 0.90 | 37.48 |
| 10,000 (20 datasets) | 1 | 0.89 | 1 | 0.89 | 1 | 0.90 | 77.00 |
| 15,000 (30 datasets) | 1 | 0.92 | 1 | 0.93 | 1 | 1 | 81.00 |
| 20,000 (40 datasets) | 1 | 0.94 | 1 | 0.95 | 1 | 1 | 117.48 |

### 4.3 Sensitivities to Measurement Noise and Temporal Undersampling

#### 4.3.1 Measurement Noise Sensitivity

As mention in Section 2.1, following Smith et al. (2011), all the simulations have Gaussian noise added to the BOLD synthetic signals to model measurement noise in the observations, such that *y* = *ỹ* + *e_m_*, where *ỹ* is the measurement-noise-free BOLD signal and *e_m_* is the measurement noise distributed as a standard Gaussian with *mean* = 0 and *std.dev* = 1. In principle, for each of the causal inference methods presented in Section 3, the corresponding estimator was derived by minimizing some error or maximizing the likelihood of a model. To do so one has to assume a generative model, and the assumed generative models do not contain measurement noise. So, adding measurement noise will be a violation of the model assumptions. In practice, some causal discovery methods will not have a good performance when model assumptions are violated, others may be more robust, and this is why simulation studies in which model assumptions are violated are conducted to assess the robustness of causal inference algorithms. Scheines & Ramsey (2016) assess the effect of increasing magnitudes of measurement noise on the performance of the FGES causal inference algorithm (Ramsey et al., 2017), and report a reduction on adjacency and orientation precision and recall, as the measurement noise becomes more prominent. They also note that the detrimental effect on accuracy of measurement noise can be partially mitigated if the sample size is increased. Zhang et al. (2017) present a formalization of a generative model with added measurement noise and derive sufficient conditions under which the causal model governing the true measurement-noise-free variables can be identified. In general, the problem of additive measurement noise arises from the fact that observed variables with measurement noise are not exact proxies of the true measurement-noise-free underlying variables and thus, the structural equations and conditional independence relations governing the true variables will not necessary hold for the observed variables. Zhang et al. (2017) and Scheines & Ramsey (2016) show how, given the difference between true and observed variables due to measurement noise, significant correlations that hold in the true variables may disappear in the observed variables (increasing the risk of false negative edges), and zero partial correlations that hold in the true variables may become significant in the observed variables (increasing the risk of false positive edges), as the magnitude of the measurement noise increases.

To have a sense of the effect of measurement noise on the estimation of cyclic structures for Two-Step and FASK, we generated data for the *Network 4* simulation without standard Gaussian measurement noise, and computed the percentage change in performance between the data with measurement noise and the data without. For comparison, lag-based procedures, MVGC and GlobalMIT, are also included. Figure 4 shows average percentage *change* of precision and recall for adjacencies, orientations and 2-cycles detection for 60 repetitions of ten datasets concatenated when the Gaussian measurement noise is removed. All considered, elimination of measurement noise decreases recall and precision for MVGC, helps GlobalMIT(FAS) minimally with precision but not recall, FASK is mostly unaffected, and helps Two-Step(Alasso) considerably across adjacencies, orientations and 2-cycle detection. The improvements for Two-Step are expected since without measurement noise the data are now from the true model, but the sensitivities for the other methods, especially MVGC and GlobalMIT are unclear. Remarkably, FASK is essentially unaffected by measurement noise, at least within the limits we have explored here.

**Figure 4:**
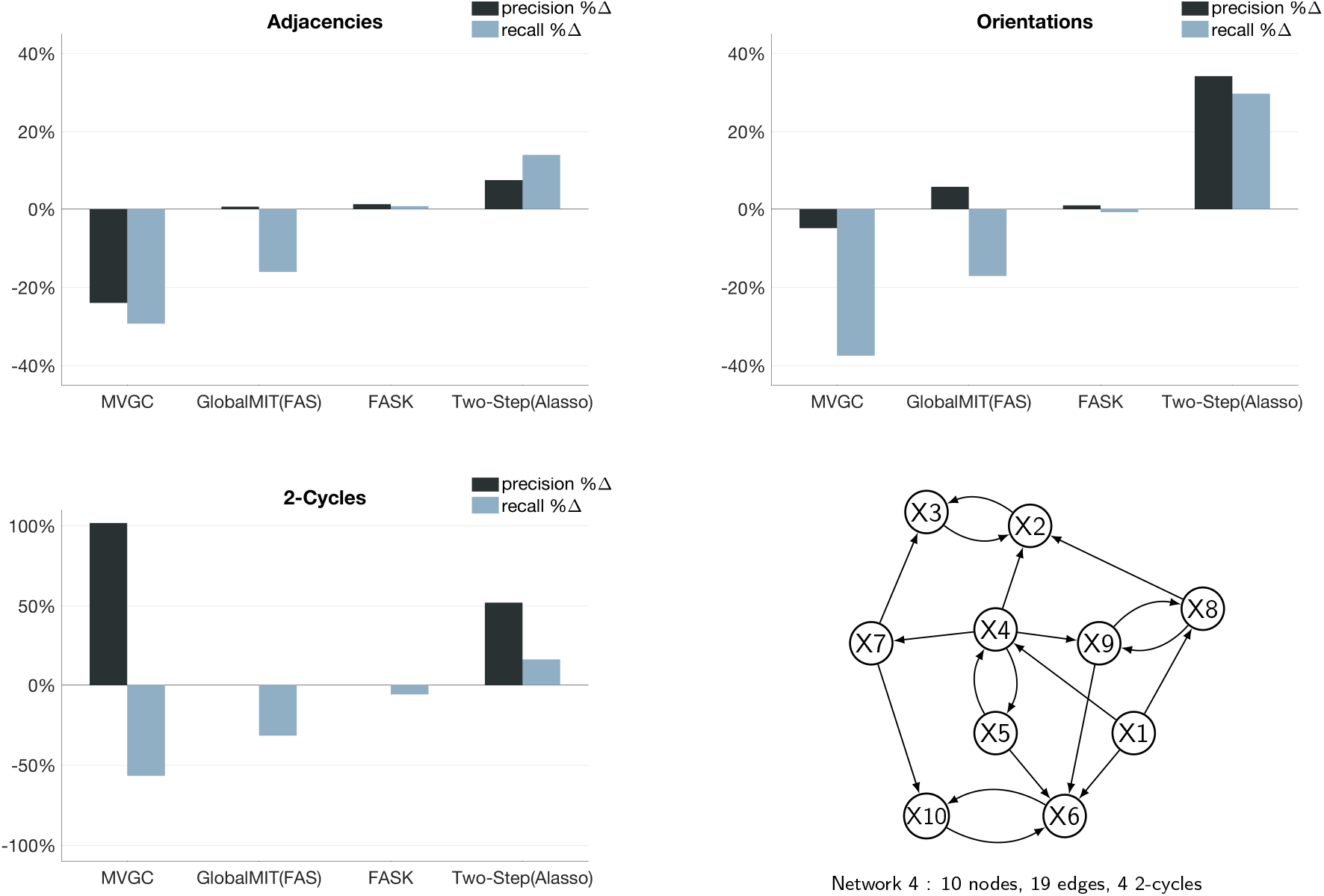
Effect of removing measurement noise. Percentage change (%Δ) of precision and recall when the standard Gaussian measurement noise is removed from the data. Results for adjacencies, orientations and 2-cycles, averaged across 60 runs of ten concatenated sessions. Data simulated from *Network 4*.

#### 4.3.2 Temporal Undersampling Sensitivity

As with measurement noise, reductions in the temporal resolution of the observed data can affect the performance of causal inference algorithms, especially for lag-based methods. As the sampling rate of the fMRI data recording gets lower, the recall of lag-based methods is expected to decrease given that the information about variables dependency recovered from previous time steps will not contain relevant causal information when the real causal processes occur at a faster rate than the sampling rate (Seth et al., 2013). In contrast, with data collected at a faster sampling rate the recall of lag-based methods is expected to improve. In fact, when the MVGC algorithm was run on noiseless 5ms resolution synthetic neuronal data from the DCM simulator, recall and precision for adjacencies, orientations and 2-cycles detection, were perfect. Following the measurement noise analysis, synthetic BOLD data was produced from *Network 4* at a lower sampling rate of TR = 3 seconds (datapoints sampled each 3 seconds), in order to compare it to the original data with a higher sampling rate of TR = 1.2 seconds. For the 3s TR simulated data, the scanning session was increased to 25 minutes to guarantee a sample size of 500 datapoints in each individual dataset, as in the original test data. Figure 5 shows average percentage *change* of precision and recall for adjacencies, orientations and 2-cycle detection for 60 repetitions of ten datasets concatenated when the sampling resolution was reduced from 1.2s TR (higher sampling resolution) to 3s TR (lower sampling resolution), for MVGC, GlobalMIT(FAS), FASK and Two-Step(Alasso). As expected, the reduction in sampling rate reduced the adjacencies, orientations and 2-cycle detection recall for the lag-based methods, MVGC and GlobalMIT(FAS), by more than 40%. In contrast, for FASK and Two-Step, which assume i.i.d., data, the reduction in sampling rate did not significantly affect their precision or recall.

**Figure 5:**
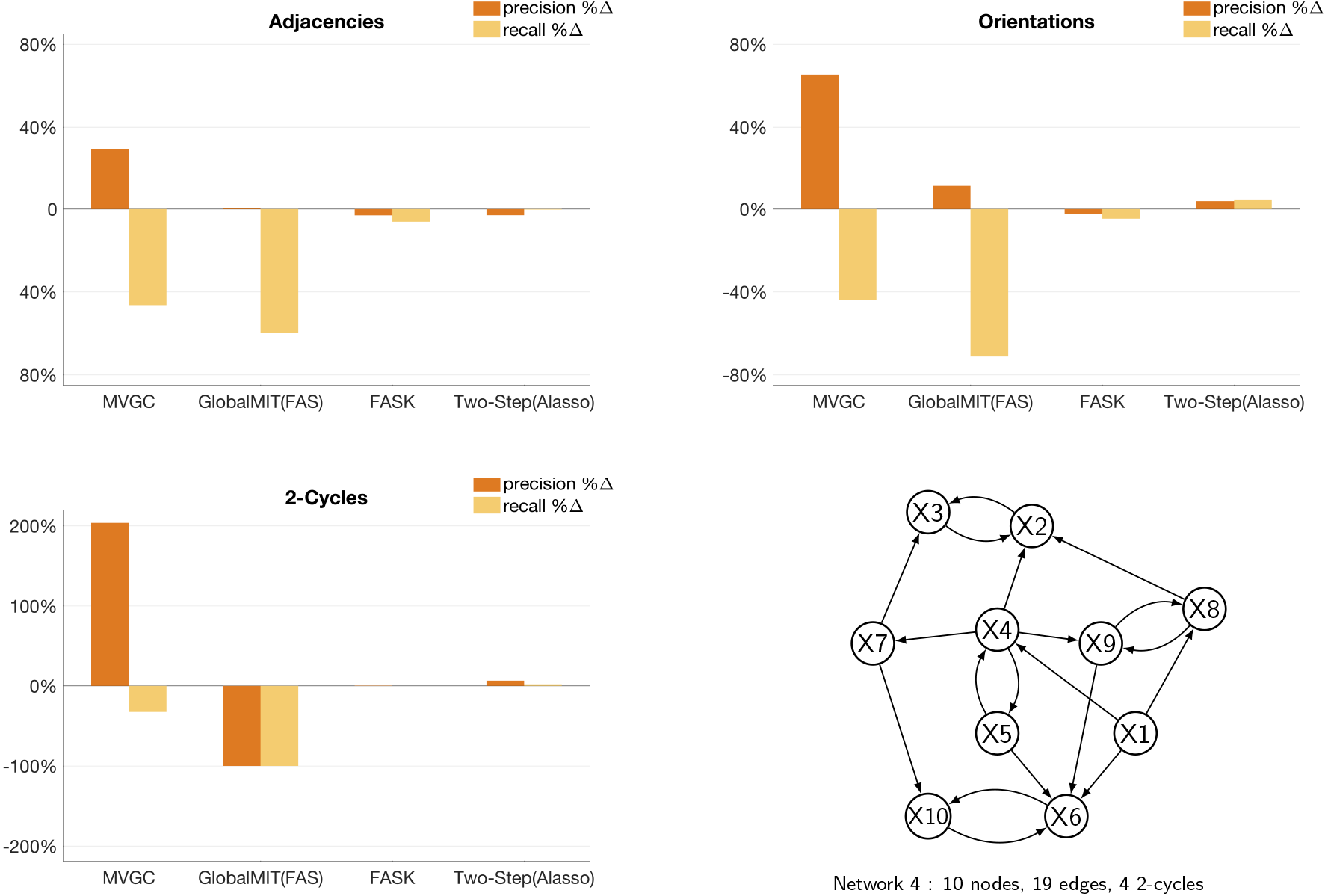
Effect of reducing the temporal sampling resolution. Percentage change (%Δ) of precision and recall when the sampling resolution is reduced from one sample each 1.2 seconds (TR =1.2s) to a slower rate of one sample each 3 seconds (TR = 3s). Results for adjacencies, orientations and 2-cycles, averaged across 60 repetitions of ten datasets concatenated. Data simulated from *Network 4*.

## 5 Empirical Data Results

### 5.1 Resting State Data Results

FASK and Two-Step(Alasso) were run on 23 repetitions of ten individual subjects selected at random, standardized individually and concatenated. The resulting 23 concatenated datasets comprise seven regions of interest from the medial temporal lobe, with 4,210 datapoints. Left and right hemispheres were analyzed separately.

In contrast to simulated data, with empirical data we do not have a fully defined ground truth to exactly assess the performance of causal search algorithms. Instead, we have partial knowledge about the presence of structural connections between brain regions derived experimentally in animal models (Naber et al., 2001; und Halbach & Albrecht, 2002), post-mortem in humans (Mufson et al., 1990; Molnár et al., 2006; Mori et al., 2017), and in-vivo in humans using diffusion imaging based tractography (Zeineh et al., 2017), which nevertheless has a systematic problem of false positive inferred structural connections (Maier-Hein et al., 2017). In some cases we can also obtain knowledge about the direction of the structural connections by tracer injection experiments in animal models (Witter & Amaral, 1991; Markov et al., 2012). This partial information about the connectivity can be used to evaluate to a certain degree the performance of search algorithms, for example, by examining the presence or absence of directed edges relative to previously established reports. In addition, we can also register robust edges that are consistently estimated under different instantiations of the data. We evaluate the performance of FASK and Two-Step(Alasso) on the medial temporal lobe data by comparing robustly estimated edges for each of the algorithms.

Graphical results for the right hemisphere medial temporal lobe are in Supplementary Material C (Section C4.2). We also include for each algorithm, each hemisphere, and for the concatenated and individual cases, the list of directed edges and their frequency of appearance across the 23 instantiations, and the list of 2-cycles ordered by frequency of appearance (Supplementary Material C, Section C4). Agreement with the partial knowledge is maximized when we considered edges that appear in 48% of the 23 repetitions. We define a robust edge as an edge estimated in 48% or more of the 23 repetitions of ten concatenated subjects, or in the case of the individual subject analysis, in 48% or more of the 23 individual subjects.

Figure 6 presents graphically, for the seven regions of interest from the left hemisphere medial temporal lobe, the directed edges estimated in 48% or more of the 23 repetitions of ten concatenated subjects. We show the most robust edges produced by FASK parameterized with a penalty discount of *c* = 1 and threshold for 2-cycle detection of α = 0.05; and compare these edges with the most robust edges estimated by Two-Step(Alasso) parameterized with λ = 2 for the **B** matrix estimation, and a threshold of 0.10 for the absolute values of the final **B** matrix.

**Figure 6:**
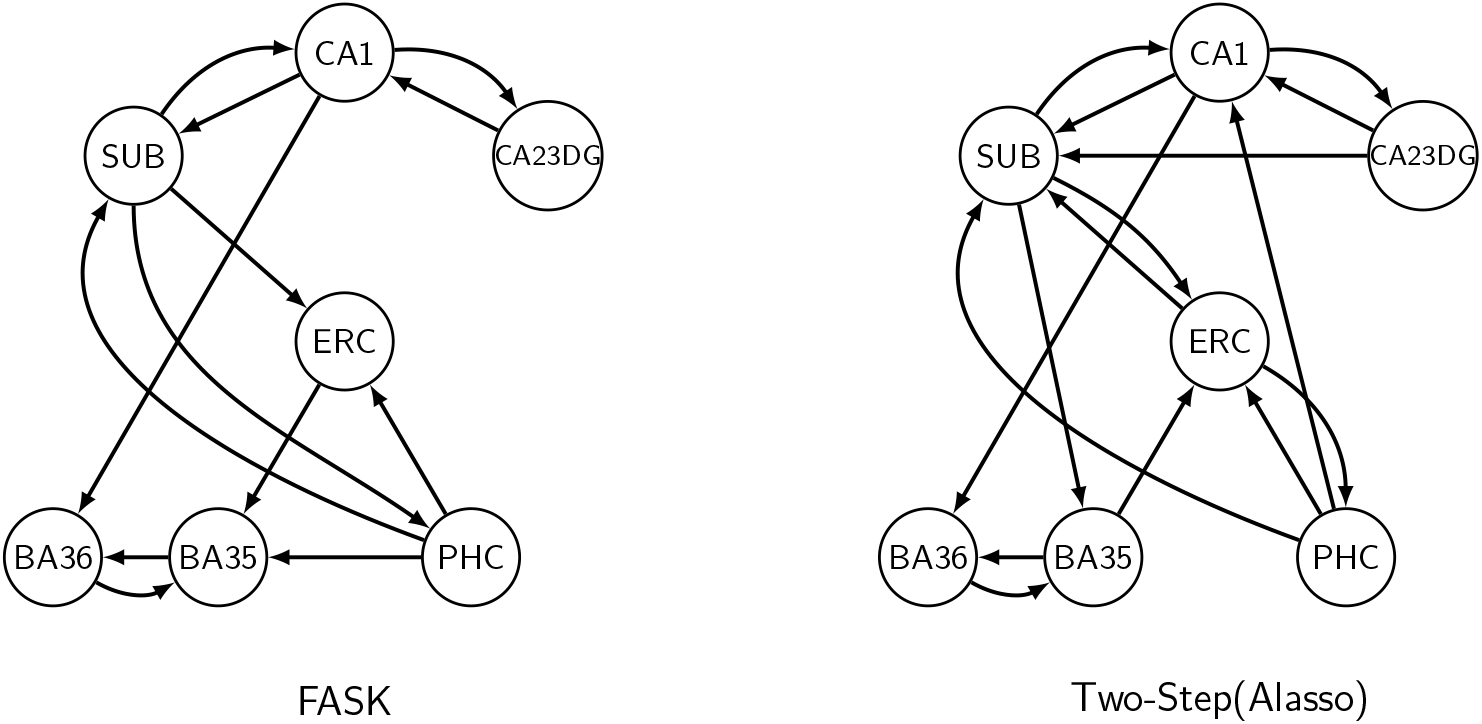
Comparison of the most robust edges estimated by FASK and Two-Step(Alasso) in 23 repetitions of ten subjects concatenated for ten regions of interest from the left hemisphere medial temporal lobe. A directed edge is depicted if it is estimated in 48% or more of the 23 repetitions of ten subjects concatenated. Regions of interest include: Cornu Ammonis 1 (CA1); CA2, CA3 and dentate gyrus together (CA23DG); entorhinal cortex (ERC); perirhinal cortex divided in Brodmann areas (BA35 and BA36); and parahippocampal cortex (PHC).

Overall both algorithms produce a closely similar set of robust edges for the medial temporal lobe left hemisphere data. With one exception every directed edge found by FASK is found by Two-Step(Alasso). For matters of graph comparison, 2-cycles count as one single adjacency. Given this, it can be seen that the adjacencies output by both algorithms are almost the same with the exception of BA35 – PHC that is found by FASK but not by Two-Step; and SUB – CA23DG, SUB – BA35 and PHC – CA1 that are found by Two-Step but not by FASK. Regarding inferred 2-cycles, both algorithms output the 2-cycles for SUB – CA1, CA1 – CA32DG and BA35 – BA36; the 2-cycle for SUB – PHC is present in the FASK output but not in Two-Step, while the SUB – ERC and ERC – PHC 2-cycles are present in Two-Step but not in FASK. The edge BA35 – ERC is oriented in opposite directions by the two algorithms. The adjacencies and orientations are consistent with the medial temporal lobe model presented in Lavenex & Amaral (2000), capturing the flow of information from the medial temporal lobe cortices (BA35, BA36, PHC) directly into the entorhinal cortex (ERC), which works as a gateway to the hippocampal formation, where the signals travel to CA23DG to CA1 (Schaffer collaterals) to the subiculum (SUB) and back to ERC, and from ERC back to BA35, BA36 and PHC. As suggested by Lavenex & Amaral (2000) there are also direct two-way connections between PHC, BA35 and BA36 (perirhinal cortex) and subiculum and CA1, captured with some degree of agreement by FASK and Two-Step. The presence of 2-cycles in the output of both algorithms is consistent with reported feedback in medial temporal lobe information processing (Insausti et al., 2017). There are two main discrepancies of our results with the standard model of the hippocampus as presented in Lavenex & Amaral (2000). First, neither FASK nor Two-Step robustly inferred the ERC–CA23DG edge (perforant pathway). This is surprising since this is the main pathway connecting the medial temporal lobe cortices with the hippocampus. One possible explanation is that the signal between these regions measured with resting state fMRI is not strong enough to be captured by our methods. Another explanation is that the regions of interest for ERC, or CA23DG or both were incorrectly spatially defined and necessary voxels driving the associations between the regions were left out. The second discrepancy is related to the CA23DG – SUB edge which shows in the Two-Step output but not in the FASK result. Again, a possible explanation of this spurious edge is related to a weak mixing of variables between CA1 and CA23DG that affects the more sensitive Alasso adjacency search in Two-Step, but not FASK which is less sensitive to weak signals. Equivalent results are observed for the right hemisphere medial temporal lobe, with the difference that the right hemisphere data show more robust 2-cycles but loses some of the direct connections from the parahippocampal and perirhinal cortices to the hippocampus. Results for the right hemisphere are shown in Supplementary Material C. Surprisingly, however Two-Step with Alasso finds the ERC – CA23DG connection in 57% of the individual datasets (Table C22 in Supplementary Material C)

The analysis using individual datasets showed similar results as the concatenated datasets, with the difference that the frequencies of estimation of the edges across the 23 individuals is lower than with the concatenated datasets. This is expected since reductions of the sample size reduces the ability to detect real effects.

### 5.2 Task Data Results

FASK and Two-Step(Alasso) were run on one repetition of 9 subjects concatenated (1,440 datapoints) from the rhyming task data, for eight bilateral regions of interest and one Input variable representing the dynamics of the stimuli presentation. FASK was run separately using FAS-stable and FAS-original (the order dependent adjacency search of the PC algorithm (Spirtes et al., 2000)) using the same penalty for the BIC score of *c* = 1 and α = 0.001 for the 2-cycle detection. Two-Step(Alasso) was run with a λ = 2 for the penalization and a threshold of the absolute values of the **B** matrix of 0.10. The resulting graph for each algorithm is shown in Figure 7 below.

**Figure 7:**
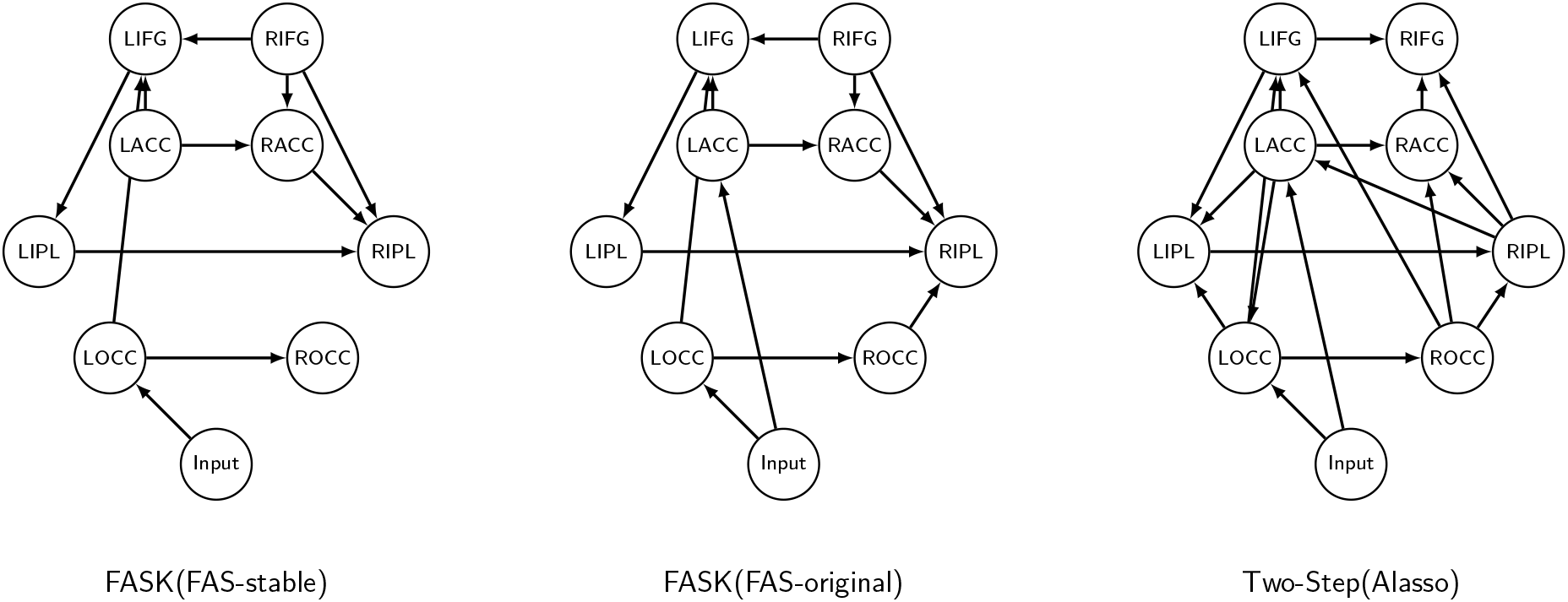
Comparison of the output graphs from FASK with FAS-stable and FAS-original, and Two-Step (Alasso) in one repetition of 9 subjects concatenated (1,440 datapoints), for eight bilateral regions of interest and one Input variable. Regions of interest include: left and right occipital cortex (LOCC, ROCC); left and right anterior cingulate cortex (LACC, RACC); left and right inferior frontal gyrus (LIFG, RIFG); left and right inferior parietal (LIPL, RIPL). The output of FASK(FAS-original) is identical to FASK(FAS-stable) with the addition of the Input → LACC and ROCC → RIPL connections.

As mentioned in Section 2.3, when analyzing task data we can model the dynamics of the task with an Input variable for which we expect feedforward edges into the regions of interest and not vice versa. This is a simple limited but gold standard test for the accuracy of orientation algorithms. Both, FASK and Two-Step(Alasso) with concatenated data, correctly output feedforward edges from the Input variable to the regions of interest. Of particular relevance is the edge going from the Input to the left occipital cortex, which we expect as this is a task in which visual stimuli are presented first. Both algorithms present feedforward edges from the left occipital cortex to the frontal lobe (LIFG, left inferior frontal gyrus), and then back to the parietal lobe (LIPL, left inferior parietal). This causal order is consistent with the rhyming task (Ramsey et al., 2010), in which first, a pair of words (or pseudowords) are presented in a screen, and then the subjects have to conclude if the words rhyme or do not rhyme. In addition to the posterior anterior direction, we also see in the graphs a cascade of homotopic effects from the left hemisphere to the right hemisphere. The graph produced by the Two-Step algorithm is more dense, which is expected given Alasso higher sensitivity to detect adjacencies under reduced sample sizes, compared to FASK. Nevertheless, the adjacencies of FASK(FAS-stable) and FASK(FAS-original) are proper subsets of the Two-Step adjacencies.

There are some difference in orientations between the algorithms in the right hemisphere, but the left hemisphere orientations are very consistent, especially regarding the flow from occipital to frontal to parietal. Interestingly, neither FASK nor Two-Step produce 2-cycles between the regions of interest. If effects are weak, FASK may miss the presence of a 2-cycle, but Two-Step(Alasso) will not, which give us some guarantee that the absence of 2-cycles for these regions under this particular task is a real connectivity property and not a problem of false negatives. These task data have been previously analyzed with algorithms that assume acyclicity (Ramsey et al., 2010), and pairwise non-Gaussian orientation algorithms (Ramsey et al., 2014), with similar results in terms of the gross flow of information from occipital to frontal regions and from left to right hemisphere.

For comparison, in Supplementary Material C (Section C5) we present results for FASK(FAS-stable) and Two-Step(Alasso) on each of the 9 individual datasets (160 datapoints). For each algorithm we report frequency of estimation for each directed edge, including 2-cycles. Even with individual data (160 datapoints) we can recover robust edges that are consistent with those obtained when we concatenated the nine subjects (Figure 7), especially those regarding homotopic connectivity from left to right hemisphere. We also observed that for FASK the directed edge from the Input variable to the left occipital region (LOCC) is hardly obtained at the individual level, while Two-Step(Alasso) has a better performance producing that edge in seven out of nine individual subjects. Two-Step seems prone to false positive edges when the sample size is small as in the individual scans.

## 6 Discussion

At present temporal resolution for fMRI data, lagged methods are unreliable in two respects. They overfit and do poorly in finding feedback cycles. The two non-Gaussian i.i.d., methods presented here, FASK and Two-Step, are far from perfect but they are more accurate than available lagged methods. FASK application is very simple with only two adjustable parameters and short running time even in large dimension problems. In contrast, due to the optimization procedure to infer the **B** matrix coefficients, Two-Step can be slow, especially with large dimension problems or when the sample size is very large. This can be mitigated by initializing the optimization of the mixing matrix with small values for the coefficients and adjusting the penalization parameters as we did here. Both FASK and Two-Step can infer cycles of any degree, and they are distinctively good at finding 2-cycles. FASK is robust to measurement noise in the observed variables, a relevant property considering that measurement noise has shown to be detrimental to the performance of certain search algorithms including Two-Step. The non-Gaussianity leveraged by FASK to orient edges comes from the skewness of the distribution around the center, if the distribution of the data is non-Gaussian but symmetric around the center, such as the Uniform distribution, FASK is not guaranteed to infer the correct orientations. In contrast, Two-Step works accurately with any type of non-Gaussian distribution, even with non-Gaussian distributions symmetric around the center.

Our results show that FAS-stable, the adjacency search of FASK, is prone to false negatives if the true structure is dense and/or the associations between variables are weak (small coefficients). This can be partially alleviated if the sample size is increased or by appropriate concatenation of scans. Nevertheless, FASK is a modular algorithm and FAS-stable can be substituted by any other adjacency search method deemed more appropriate if we expect weak effects or a very low sample size. Two-Step is also modular and if required, the Alasso procedure may be replaced by any other adjacency search method, as we did here with Two-Step(FAS).

The Two-Step algorithm outputs estimates of connection strengths (weighted directed matrix), which may be relevant for some applications, for example, connectivity-based classification. FASK does not give estimates of coefficients, its output is a possibly cyclic graph that can be encoded as a binary directed matrix. Notwithstanding, the graph output by FASK can be parameterized as a structural equation model and coefficients estimated from the data. We implement FASK in Java, but the 2-cycle detection and left-right rules are flexible enough to be implemented in other programming languages such as R, Python or Matlab. Two-Step was implemented in Matlab but it can also be implemented in R, Python or Java as long as the optimization method used has efficient running times.

In theory, Two-Step accurately finds causal connections among measured variables in the presence of unrecorded, unknown confounding variables, but we have not tested that aspect of the procedure here. None of the other procedures tested here have any theoretical guarantees when there are unrecorded confounders. The FGES algorithm (Ramsey et al., 2017) has been run on fMRI data for all voxels in the human cortex, but assumes no unknown confounders and no feedback. Notably, runs of FGES with higher penalization produced sparser graphs that were almost proper subsets of the denser graphs obtained with lower penalization, indicating a “vertical” stability to the procedure. FASK is sufficiently fast to be run at the voxel level with approximately 51,000 variables, and tests of its “vertical” stability are needed but not provided here. Still further, researchers need a reasonably accurate procedure that can be run on such high-dimensional problems and is robust both to feedback cycles and to unmeasured confounders.

## Supplementary Material A

### The FASK Algorithm

The Fast Adjacency Skewness (FASK) procedure is given as algorithms 1, 2, 3 and 4. The idea is as follows: First, we run Fast Adjacency Search stable (FAS-stable) on the data, producing an undirected graph. FAS-stable is the order independent adjacency search of the PC-stable algorithm (Spirtes et al., 2000; Colombo & Maathuis, 2014). We use the linear, Gaussian BIC score as a *conditional independence test* with a specified penalty discount (See Section 3 in the main text). This yields undirected graph *G*_0_. The reason FAS-stable works for sparse cyclic models where the linear coefficients are all less than 1, is that correlations induced by long cyclic paths are statistically judged as zero, since they are products of multiple coefficients less than 1. One then orients each of the adjacencies in *G*_0_ as a 2-cycle or a directed edge left or right. Taking up each adjacency in turn, one tests to see whether the adjacency is a 2-cycle using the 2-cycle Detection. If so, one adds edges *X → Y* and *X ← Y* in the output graph *G*_1_. If not, one applies the Left-Right orientation rule and orients *X → Y* or *X ← Y* in the output graph *G*_1_. *G*_1_ is a fully oriented graph. For some models, where the true coefficients of a 2-cycle between *X* and *Y* are close but opposite in sign, a correlation test may eliminate the edge between *X* and *Y* when in fact a 2-cycle exists. In this case, we check explicitly whether *corr*(*X, Y* |*X* > 0) and *corr*(*X, Y* |*Y* > 0) differ by more than a set amount of 0.3. If so, we add the adjacency to the graph and orient it using the aforementioned rules. The idea for the heuristic extra adjacency rule is that if X and Y are independent, then |*corr*(*X, Y* |*X* > 0) − *corr*(*X, Y* |*Y* > 0)| = 0, so if this difference is not zero, there must be a trek between X and Y. If the difference is very different from zero, we infer a very short path and so add an adjacency. The specific threshold 0.3 is a parameter than can be tuned.

A comment on notation: When we write *E*(*X|X* > 0) we mean *E*(*X*) under the condition *X* > 0, and similarly for *corr*(*X, Y* |*X* > 0). When we write *corr*(*X, Y* |*Z, X* > 0) we mean *corr*(*X, Y |Z*) under the condition *X* > 0.

#### Algorithmal 1 FASK Algorithm

~~~
1: *D* a continuous dataset, *c* a penalty discount for a SEM-BIC score, α a threshold level for the 2-cycle detection rule, δ a tunable parameter for the Left-Right rule
2: **procedure** FASK(*D, c, α*) ▹ Returns a fully oriented graph possibly with cycles.
3:          Center *D*
4   *G*_0_ ← FAS-stable(*D, c*)
5:  *G*_1_ ← <>
6:    **V** ← variables in *D*
7:    **for** *i* = 1 to |**V**| − 1 **do**
8:        **for** *j* = *i* + 1 to |**V**| **do**
9:           *X* = *i*th variable in **V**
10:          *Y* = *j*th variable in **V**
11:            **if** adjacency *X − Y* is in *G*_0_ or |*corr*(*X, Y |X* > 0) − *corr*(*X, Y |Y >* 0)| > 0.3 **then**
12:                 **if** TwoCycle(*D, X, Y, G*_0_,α) **then**
13:                            Add *X → Y* and *X Y* to *G*_1_
14:                 **else if** LeftRight(*D, X, Y*,δ) **then**
15:                            Add *X → Y* to *G*_1_
16:                 **else**
17:                            Add *X ← Y* to *G*_1_
18:                 **end if**
19:             **end if**
20:       **end for**
21:  **end for**
22:  **return** *G*_1_
23: **end procedure**
~~~

#### Algorithm 2 FAS-stable Algorithm

~~~
1: *D* a continuous dataset, *c* a penalty discount for a SEM-BIC score.
2:   **procedure** FAS-stable(*D, c*)
3:       **V** ← variables in *D*
4:     *G*_0_ ← complete undirected graph over **V**
5:       **for all** *depth* = 0, 1, …, so long as adjacencies can be removed **do**
6:           **for all** *X* ∊ **V do**
7:                *a*(*X*) ← *adj*(*X, G*_0_) ▹ Adjacent variables set for *X* in *G*_0_.
8:           **end for**
9:           **for all** *X, Y*, **S** for *X, Y* ∊**V, S** ⊆ *a*(*X*) \ {*Y*}, |**S**| = *depth* **do**
10:                **if** X is independent of Y conditional on **S then** ▹ Use SEM-BIC score with penalty *c*.
11:                         Remove adjacency *X − Y* in *G*_0_
12:                **end if**
13:          **end for**
14:   **end for**
15:   **return** *G*_0_
16: **end procedure**
~~~

#### Algorithm 3 2-Cycle Detection Rule

~~~
1: *D* a continuous dataset, a threshold level, *G*_0_ an undirected graph, *X, Y* a pair of adjacent variables.
2:   **procedure** TwoCycle(*D, X, Y, G*_0_,α) ▹ Returns *true* if *X* ⇄ *Y*.
3:       **for all S** ⊆ {*adj*(*X, G*_0_) ∪ *adj*(*Y, G*_0_)} \ {*X, Y*} **do**
4:        *c ← corr*(*X, Y* |**S**)
5:        *c*_1_ ← *corr*(*X, Y* |**S**, *X* > 0)
6:        *c*_2_ ← *corr*(*X, Y* |**S**, *Y* > 0)
7:        *z* ← 0.5(*ln*(1.0 + *c*) − *ln*(1.0 − *c*)) ▹ Fisher’s z-transformation.
8:        *z*_1_ ← 0.5(*ln*(1.0 + *c*_1_) − *ln*(1.0 − *c*_1_))
9:        *z*_2_ ← 0.5(*ln*(1.0 + *c*_2_) − *ln*(1.0 − *c*_2_))
10:       *n* ← #*X* ▹ #*i* sample size for variable *i*.
11:       *n*_1_ ← #*X* > 0
12:       *n*_2_ ← #*Y* > 0
13:       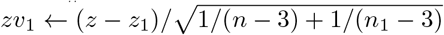 ▹ Two-sample testing for diff. in correlations.
14:       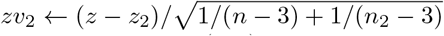
15:      *p*_1_ ← two-tailed *N*(0, 1) *p*-value for *zv*_1_
16:      *p*_2_ two-tailed *N*(0, 1) *p*-value for *zv*_2_
17:      *rejected*_1_ *p*_1_ < α
18:      *rejected*_2_ *p*_2_ < α
19:      *possibleTwoCycle ← false*
20:       **if** *rejected*_1_ ∧ *rejected*_2_ **then**
21:           *possibleTwoCycle ← true*
22:      **else if** *zv*_1_ < 0 ∧ *zv*_2_ > 0 ∧ *rejected*_1_ **then** ▹ conditions in case of *control* 2-cycles.
23:           *possibleTwoCycle ← true*
24:      **else if** *zv*_1_ > 0 ∧ *zv*_2_ < 0 ∧ *rejected*_2_ **then**
25:           *possibleTwoCycle ← true*
26:      **end if**
27:      **if** not *possibleTwoCycle* **then**
28:             **return** *false*
29:      **end if**
30:   **end for**
31:   **return** *true*
32: **end procedure**
~~~

#### Algorithm 4 Left-Right Orientation Rule

~~~
1: *D* a continuous dataset, *X, Y* a pair of adjacent variables, δ in (−1, 0) a tunable parameter.
2:  **procedure** LeftRight(*D, X, Y*,δ) ▹ Returns *true* iff the edge should be oriented *X → Y*.
3:  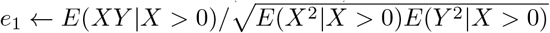
4:  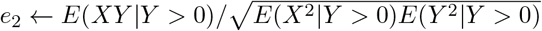
5:   *lr* := *e*_1_ − *e*_2_
6:   *c* := *corr*(*X, Y*)
7:   *sx* := *skewness*(*X*)
8:   *sy* := *skewness*(*Y*)
9:   *c* := *c * sign*(*sx*) * *sign*(*sy*)
10:  *lr* := *lr * sign*(*c*)
11:    **if** *c* < δ **then**
12:      *lr* := *−lr*
13:    **end if**
14:    **return** *lr* > 0
15: **end procedure**
~~~

### 6.1 The Left-Right Orientation Rule

Assuming *X → Y* or *X ← Y* without a 2-cycle between *X* and *Y*, we give a procedure to orient the adjacency left or right. To do this, we define a certain relationship that must obtain if the probability distributions of the variables *X, Y* and their residuals, *e_X_* and *e_Y_*, are *smoothly positively skewed*, defined thus:

#### Definition 1.

*We say a variable X with p.d.f., f_X_*(*x*) *centered around c is smoothly positively skewed if for every* 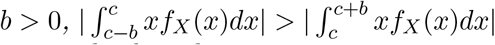. *X is smoothly negatively skewed if −X is smoothly positively skewed*.

Variables that are smoothly positively skewed are positively skewed in Pearson’s sense, 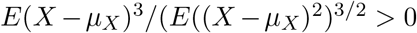 (Pearson, 1894). There may be variables skewed positively in Pearson’s sense that are are not smoothly positively skewed, as for instance case where *f_X_*(*x*) has a small bump directly to the right of the center *c* but is zero to the left within some interval (*c − b, c* + *b*) but overall skewed in Pearson’s sense. In such cases, for small *b*, it may be that 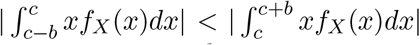, whereas for sufficiently larger *b*, this will not hold.

These cases we set aside.

We assume the following for pairwise direct causal relationships:

1. *Y* = *aX* + *e_Y_*.
2. *X* and *e_Y_* are independent: *X* ╨ *e_Y_*.
3. *X* and *e_Y_* are centered.
4. *X* is smoothly positively skewed.
5. *e_Y_* is smoothly positively skewed.
6. *a* > 0.

#### Lemma 1.

*Assume 1-6, then E*(*Xe_Y_* |*Y* > 0) < 0.

#### *Proof*.

See Figure A1 for *X* and *eY* centered around zero. The *E*(*Xe_Y_* |*Y* > 0) integrates over regions *A, B, C* and *D*, with *x*_0_, *e_y_*_0_ > 0. The total expectation can be decomposed into the expectation for regions *A* and *B* taken together and the expectation for regions *C* and *D* taken together. The expectation for regions *A* and *B* taken together is given by:

**Figure A1.**
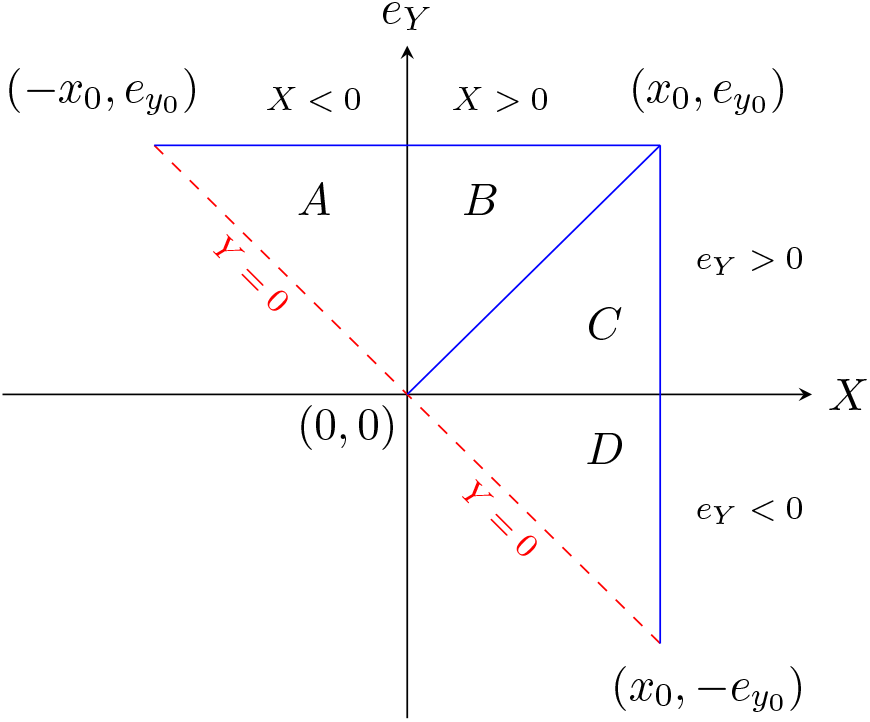
Integration regions *A, B, C, D* for *E*(*Xe_Y_* |*Y* > 0). The dashed line indicates *Y* = 0 ⇒ *e_Y_* = −*aX*. The area to the right of the dashed line correspond to *Y* > 0.

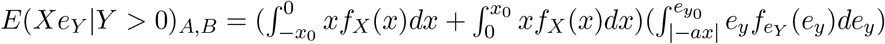

The sum inside the first parenthesis we know to be negative because of assumption 4 and definition 1. The term inside the second parenthesis is positive since *e_y_* integrates over a positive range. Thus, the total product will be negative. Equivalently, for regions *C* ad *D* taken together: 

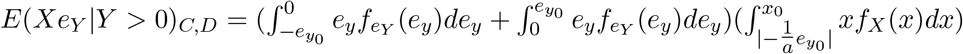

Equivalently, by assumption 5 and definition 1 we know the sum inside parenthesis is negative, and the term inside the second parenthesis is positive since *x* integrates over a positive range. Thus, the total product is negative. Therefore, *E*(*Xe_Y_* |*Y* > 0) = *E*(*Xe_Y_* |*Y* > 0)_*A,B*_ + *E*(*Xe_Y_* |*Y* > 0)_*C,D*_ < 0. □

Lemma 1 may be weakened in several ways. All that is necessary is that *E*(*Xe_Y_* |*Y* > 0) integrates to a negative number over regions *A, B, C*, and *D*. So if *X* is smoothly positively skewed but *e_Y_* is symmetric about zero, this integral will still be negative, and the same is true in reverse. Also, clearly *X* or *e_Y_* do not need to be smoothly positively skewed over their entire domains; if one or the other is smoothly positively skewed under any restriction of its domain, the conclusion will follow, so long as it is constant over the rest of its domain. In addition, small aberrations may be tolerated in the smooth skewness condition, so long as they are overpowered sufficiently soon in the integration.

Note that additional paths between *X* and *Y* do not alter the conclusion of Lemma 1, so long as assumptions 2, 3, 4, 5 and 6 are upheld. (Confounder paths do not satisfy these conditions, since *X* is not independent of *e_Y_* in these cases.) Here, we simply use the *marginal* relationship between *X, Y*, and *e_Y_* for Figure A1.

#### Lemma 2.

*Assume 1–6, then* 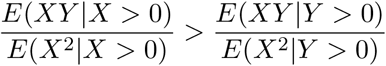.

#### Proof.

Multiplying each term by *X* and taking expectations conditional on *Y* > 0 for *Y* = *aX* + *e_Y_*, we have that *E*(*XY |Y* > 0) − *aE*(*X*^2^|*Y* > 0) = *E*(*Xe_Y_* |*Y* > 0) < 0, by Lemma 1, so 

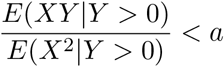

If now we condition on *X* > 0 we have *E*(*XY* |*X* > 0) − *aE*(*X*^2^|*X* > 0) = *E*(*Xe_Y_* |*X* > 0) = 0 by assumption of *X* ╨ = *e_Y_*, so 

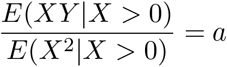

Consequently, 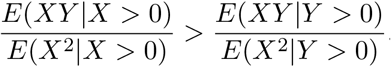.

Lemma 2 may be weakened as indicated for Lemma 1.

#### Theorem 1.

*Assume 1–6, and additionally that Y and e_X_ are smoothly positively skewed and that X and Y do not form a 2-cycle, then* 

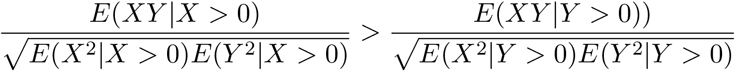

#### Proof.

If *X → Y*, we have by Lemma 2 that 
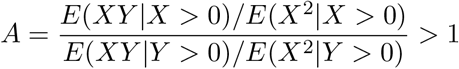
and following the same strategy as in Lemma 2 but multiplying the terms by *Y* instead of *X*, we can derive, 

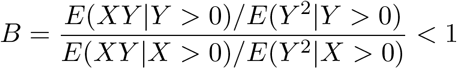

This suggests a rule that if *A > B* we orient *X → Y*; if the reverse, *X ← Y*. *A > B* simplifies to 

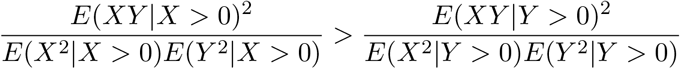

From this, the conclusion of the theorem follows.

Theorem 1 may be weakened as indicated for Lemma 1.

A comment should be added as to how negative coefficients and negative skewnesses (violations of assumptions 4 – 6) can be handled. We may assume fairly in most cases that *X* and *e_X_* are either both smoothly positively skewed together or both smoothly negatively skewed together, and that the same is true of *Y* and *e_Y_*. Then if *X* and *Y* are both smoothly positively skewed and *a* > 0, Theorem 1 tells us that *X → Y*. If *a* < 0, *E*(*Xe_Y_*) is negative for regions A and **B** and positive for the regions now symmetrically opposite to regions C and D. So whether the total integral for *E*(*Xe_Y_*) is positive or negative will depend on the value of *a*. There is a threshold δ, empirically tuned for particular types of data, for *a* greater than which this integral is negative and less than which it is positive. A δ of about −0.2 is suitable for our fMRI simulations. If *a < δ*, then the orientation of the causality needs to be reversed. If *X* and *Y* are both smoothly negatively skewed, the same conclusion follows, since *X* and *Y* will both need to be replaced by their negations for Lemmas 1 and 2 and Theorem 1. For judging the δ threshold condition, one will need to replace *X* or *Y* or both by their negations when using correlation of *X* and *Y* as an estimate of the coefficient *a* for Lemmas 1, 2 and Theorem 1. The formulation of these ideas as an algorithm is given in Algorithm 4.

If the data is centered, the formulation in Theorem 1 is very close to the claim that *X → Y* if and only if *corr*(*X, Y* |*X* > 0) > *corr*(*X, Y* |*Y* > 0). The latter claim will be true as well, if *X* and *e_Y_* are not centered around zero but they are smoothly positively skewed. In practice, we find that the formulation in Theorem 1 works better.

The Left-Right rule is similar in many ways to the pairwise orientation rules proposed by Hyv¨arinen and Smith (2013), though differently motivated. It is closest to their Robust Skew rule substituting *h*(*x*) = *max*(0*, x*) for *g*(*x*) = *log*(*cosh*(*max*(0*, x*))), a very similar function. There are some differences suggested by the algebra. In simulation, it has a noticeable advantage for the data we are concerned with. Also, the assumptions of Theorem 1 may be weakened as indicated. For these reasons, we prefer it.

### Left-Right Orientation Rule with Confounding

Although it is not exact, the above argument can be adjusted for the case where we have a linearly added confounding term like so, 

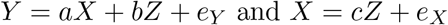

Assume *X* ╨ *e_Y_*, *Z*╨*e_X_*; *a* + *bc* > 0; and *Z, e_X_* and *e_Y_* centered and smoothly positively skewed. In the equation for *Y* multiply each term by *X* and take expectations to get, 

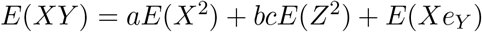

Conditioning on *X* > 0, together with the *X* ╨ *e_Y_* assumption, we have 

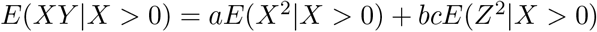

When conditioning on *Y* > 0 we get, 

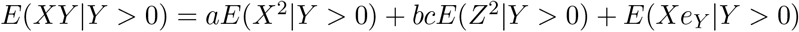

The question is if under the presence of a confounder, *E*(*XY* |*X* > 0) > *E*(*XY* |*Y* > 0). This question reduces to whether 

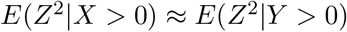

Algebraically, these expressions are not identical, but the conclusion of Lemma 1 still either holds directly (in the cases we investigate in this paper) or is very close. As an example where one expects strong confounding, let the errors all be distributed as *Beta*(1, 5) (which has a small variance and positive skew), and let coefficients *a, b, c* be drawn from (−0.9, −0.1) ∪ (0.1, 0.9), then over 100 runs of the simulation for this linear model, we have *mean*(*E*(*Z*^2^|*X* > 0)) = 0.0300, *variance*(*E*(*Z*^2^|*X* > 0)) = 0.0002, *mean*(*E*(*Z*^2^|*Y* > 0)) = 0.0377, *variance*(*E*(*Z*^2^|*Y* >0)) = 0.0003, with a mean difference of −0.0077. If the product of the coefficients *bc* is positive, as is generally the case with our simulations, the breaks in the right direction, but even if the product *bc* is negative, *bc*(*E*(*Z*^2^|*X* > 0) − *E*(*Z*^2^|*Y* > 0)) is quite small, since *b* and *c* are small in absolute value.

### 2-Cycle Detection Rule

In order to apply the Left-Right rule described above, we need to rule out the possibility that *X → Y* and *X ← Y* forming a 2-cycle. Fortunately, we can detect 2-cycles using ideas from the Left-Right rule, though extra conditioning on combinations of adjacent variables needs to be done to block the effect of other possible cyclic paths between *X* and *Y*. The 2-cycle detection rule *without conditioning* only discovers whether a cyclic path exists between *X* and *Y* but cannot distinguish if such a path is a 2-cycle or a higher degree cycle.

The 2-cycle detection rule is based on the claim that if there are no additional cycles in the graph and *X → Y* and *X ← Y*, then *corr*(*X, Y*) ≠ *corr*(*X, Y* |*X* > 0) and *corr*(*X, Y*) ≠ *corr*(*X, Y* |*Y* > 0). Here, we use centered correlations as opposed to the uncentered correlations of Theorem 1 because the calculations are very close, and with centered correlations we can calculate z-statistics. As a result, the calculations are approximations, but very good ones.

The justification is based on arguments for the Left-Right rule. Assume *X, Y*, *e_X_*, and *e_Y_* centered and smoothly positively skewed. If *X → Y*, with a linear model *Y* = *aX* + *e_Y_*, *a* > 0, and *X* ╧ *e_Y_*, then *E*(*XY*)*/E*(*X*^2^) = *a*. Additionally, by Lemma 2 we know that *E*(*XY |X* > 0)/*E*(*X*^2^|*X* > 0) = *a*, and *E*(*XY* |*Y* > 0)/*E*(*X*^2^|*Y* > 0) < *a*. Considering these relationships, if *X → Y* then these two conditions will hold: 

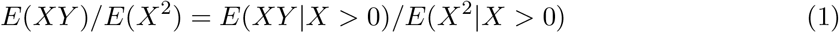

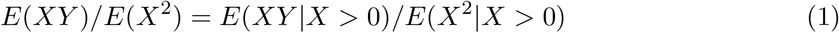

But for a 2-cycle, in which *X → Y* and *X ← Y*, with linear models *Y* = *aX* + *e_Y_* and *X* = *bY* + *e_X_*, these two conditions do not hold anymore. Condition (1) fails because *X* is not independent of *e_Y_* anymore, the necessary assumption for determining the equality. Condition (2) will still be an inequality but possibly with direction flipped depending on the relative strength of the coefficients. The important fact is that under a 2-cycle, conditions (1) and (2) will become inequalities that can be tested. In general, if *X → Y* and *X ← Y*, we expect: 

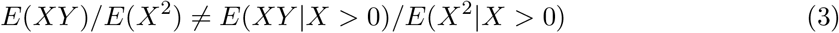

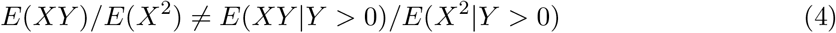

We have seen in simulations two problems with testing (3) and (4) as they are. First, small decimal differences make the conditions to hold even without the presence of a 2-cycle. Second, the conditions may hold even without a 2-cycle if the sample is not large enough. To overcome these problems we approximate the ratios in (3) and (4) with conditional and unconditional correlations and perform statistical tests of the difference of correlations. In particular, using centered rather than uncentered correlations, we test if *corr*(*X, Y*) ≠ *corr*(*X, Y* |*X* > 0) and *corr*(*X, Y*) ≠ *corr*(*X, Y* |*Y* > 0).

Specifically, we convert these correlations to z-scores and make two-tailed hypothesis testing of the differences of the conditioned z-score with the unconditioned z-score for *X* > 0 and *Y* > 0 correspondingly. If both differences are judged significantly different from zero, we may conclude that *X* and *Y* form a 2-cycle.

The one problem with this approach is that the coefficients in opposite directions in the 2-cycle may have opposite signs, a type of cycle called *control cycle*. In control 2-cycles, one or the other of the conditioned correlations might not be distinguishable from the unconditioned correlation because of cancellation or near cancellation of coefficients. That is, one might reject the null of the difference test only for *X* > 0 or *Y* > 0, but not both, therefore missing the presence of a 2-cycle. However, in control 2-cycles the differences in correlations will be of opposite sign, unlike the unidirectional *X → Y* case, in which these differences will be of the same sign. Leveraging this information it is possible to distinguish a control 2-cycle from an unidirectional edge. The exact condition is given in the pseudo-code for Algorithm 3. Following this approach, all 2-cycles in a causal graph that contains no other cycles are identifiable.

If the graph contains other cyclic paths between *X* and *Y*, we simply need to condition on some combination of the adjacent variables of *X* and the adjacent variables of *Y*, other than *X* and *Y* themselves, to block additional d-connecting cyclic paths between *X* and *Y*. The theory for this is given in (Richardson, 1996), for the Cyclic Causal Discovery (CCD) algorithm. For the 2-cycle detection rule, we may try every combination of such adjacent variables, looking for a conditioning set where the 2-cycle condition above fails to obtain. If we find such a conditioning set, we stop and apply the Left-Right rule. If there is no such conditioning set, we conclude that the adjacency is a 2-cycle. This idea is incorporated into the pseudo-code for Algorithm 3.

## Supplementary Material B

### B1 Parameter Tuning

All the search procedures tested here require at least one user-input parameter; some require two or three. In order to give fair comparisons under parameters optimal for the respective algorithms and kind of data, datasets were created exactly as the ones described in Section 2.1, but changing the coefficient values range, by sampling them from a Gaussian distribution with *mean* = 0.4, *std.dev* = 0.1, and truncated between 0.2 and 0.6. We refer to these data as *training* datasets.

Ten datasets were selected at random without replacement from 60 individual training datasets for *Network 5* and *Network 7* exclusively. *Network 5* is a representative case for a structure with 2-cycles; and *Network 7* represents structures with higher degree cycles.

Each of the search algorithms was run on the training datasets with varying parameters. The Matthews correlation coefficient (MCC) (Matthews, 1975) computed from the confusion matrix, was used to assess the performance of the methods. The MCC ranges from +1, when an algorithm perfectly recovers the true graph without false positive edges, to –1, when the algorithm produces false positive edges without recovering any of the true edges in the graph. The average of the Matthews correlation coefficient (MCC) for adjacency estimation and orientation estimation was computed for each combination of parameters, and the best-performing parameters were selected. In cases where two or more combinations of parameters performed equally well, the combination which was present in the list of highest performances for both networks was selected. If multiple combinations appeared in both high-performance lists, each combination of parameters was used in the test phase. Matthews correlation coefficient contains an implicit utility trade-off between recall and precision that may not be appropriate for some settings.

For both implementations of Two-Step (FAS-stable and Alasso), multiple combinations of parameters performed equally well, but there was no overlap between the high-performing lists of parameters for *Network 5* and *7*, and thus no combination of parameters could be selected following the above procedure. To solve this problem, Two-Step with FAS-stable and with Alasso were run on ten randomly selected training datasets without replacement generated from the *SmallDegree* macaque network with coefficients sampled from a Gaussian distribution with *mean* = 0.4, *std.dev* = 0.1 and values truncated between 0.2 and 0.6. The highest-performing combination of parameters on this network was selected for both implementations of Two-Step.

Using the parameter values obtained under the process described above, all the algorithms were run on test data from *Network 1* to *9* from Figure 1.

### B2 Parameter Tuning for Macaque-Based Simulations

As noted in Section 2.2, for the macaque-based simulations the coefficients were restricted to very small values to guarantee the stability of the synthetic BOLD signals. Given this restriction on the coefficients we did not find it feasible to create a training set based on a different range for the coefficients, as we did in Section B1. Instead, we followed this procedure: We centered and concatenated the first ten datasets for each simulation in Section 2.2, and used the concatenated data as training data. The remaining 50 individual datasets were used as the test set. FASK and Two-Step with Alasso were run for different parameters values and we chose those that maximize the average of the Matthews correlation coefficient for adjacencies and 2-cycle orientations in the case of FASK, and for adjacencies and the complete orientations for Two-Step, we explain the reason for this difference below. For FASK, the penalty discount *c* controls the sparsity of the adjacency search, while the α threshold controls the sensitivity for the difference of correlations used in the the 2-cycle detection step. The left-right orientation rule does not require a parameter. Thus, we look for the pair of parameters that maximize the average of the Matthew’s correlation coefficient for adjacencies and 2-cycles (MCCa2c). This process is illustrated in Figure B1, with heatmaps for the MCCa2c values obtained under different combinations of values for the penalty discount *c* and *α* the threshold for 2-cycle detection, for each of the four macaque-based networks in 2.2.

**Figure B1.**
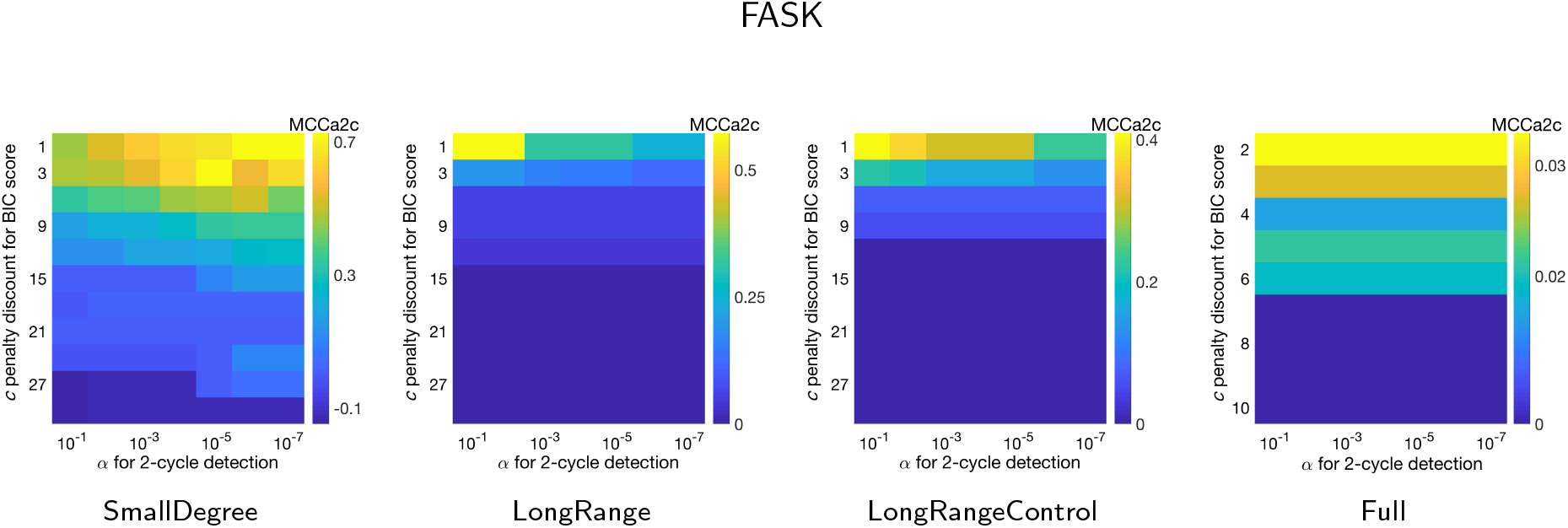
Average Matthew’s correlation coefficient for adjacencies and 2-cycles (MCCa2c) for the FASK algorithm, under different combinations of *c* penalty discount for the BIC^*^ score (Y axis) and α significance threshold for the 2-cycle detection rule (X axis). Results for the four macaque-based networks from Section 2.2

For Two-Step with Alasso, we vary the sparsity penalty λ of the **B** matrix estimation, and the threshold for the absolute values in the final **B** matrix. In contrast to FASK, the detection of 2-cycles is part of the global ICA estimation procedure of Two-Step, and so we use the average of the Matthew’s correlation coefficient for adjacencies and orientations (MCCao). Figure B2 shows the parameter tuning heatmaps for combinations of and the threshold for the absolute values of the **B** matrix, for each of the four macaque-based networks simulations in 2.2.

**Figure B2.**
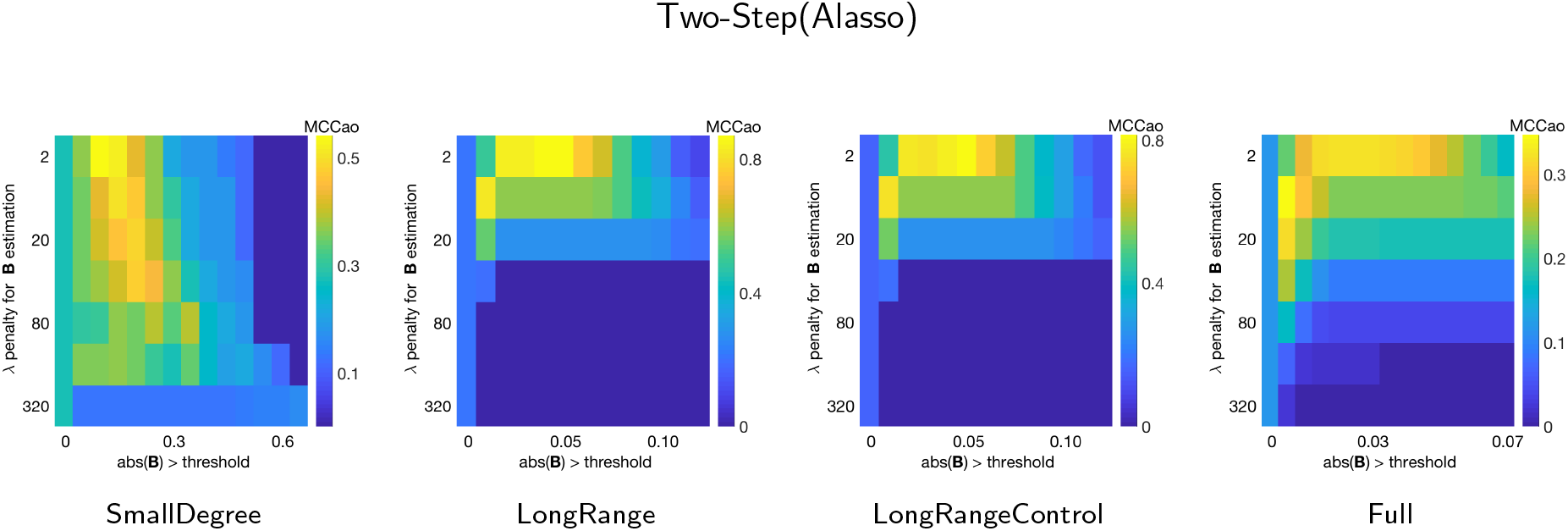
Average Matthew’s correlation coefficient for adjacencies and orientations (MCCao) for the Two-Step(Alasso) algorithm, under different combinations of the sparsity parameter λ (Y axis) and the threshold for the absolute **B** matrix values (X axis). Results for the four macaque-based structures from Section 2.2.

## Supplementary Material C

### C1 Results for simple networks with 2-cycles

Complete results for analysis shown in Figure 2, for *Network 1* to *6* and their amplifying and control variants. “amp” and “cont” after each network label indicates if the network has an amplifying (excitatory) or a control (inhibitory) cycle. For *Network 5* two additional control variants were tested, one where the 2-cycle is a control cycle with coefficients +0.3 and –0.7 (Net5_c_p3n7); and its inverse where the 2-cycle coefficients are +0.7 and –0.3 (Net5_c_p7n3). Tables show for each algorithm tested the average results over 60 repetitions of ten concatenated datasets from the corresponding simulations, for precision and recall of adjacencies, orientations, and 2-cycle detection; and running times in seconds. The average over all ten networks for each algorithm, is presented at the right of the table, followed by the corresponding standard deviation. These averages are plotted in Figure 2.

**Table C1:**
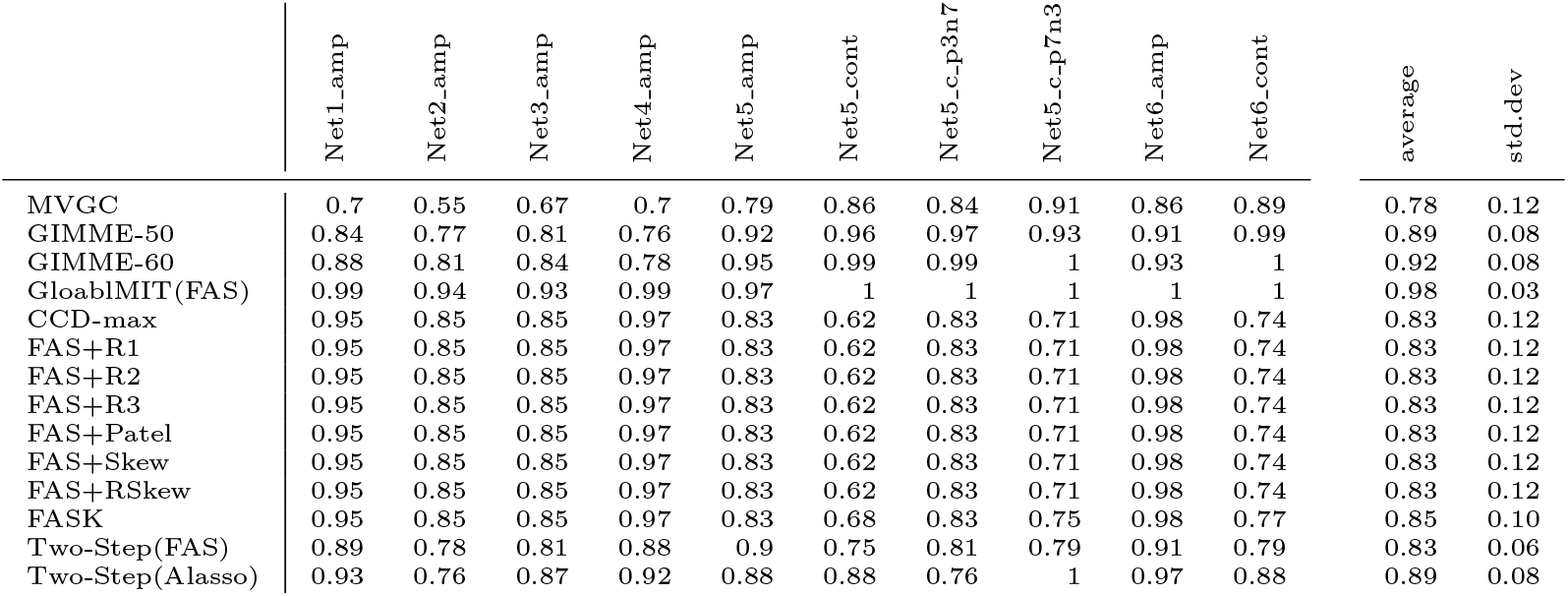
Adjacency Precision

**Table C2:**
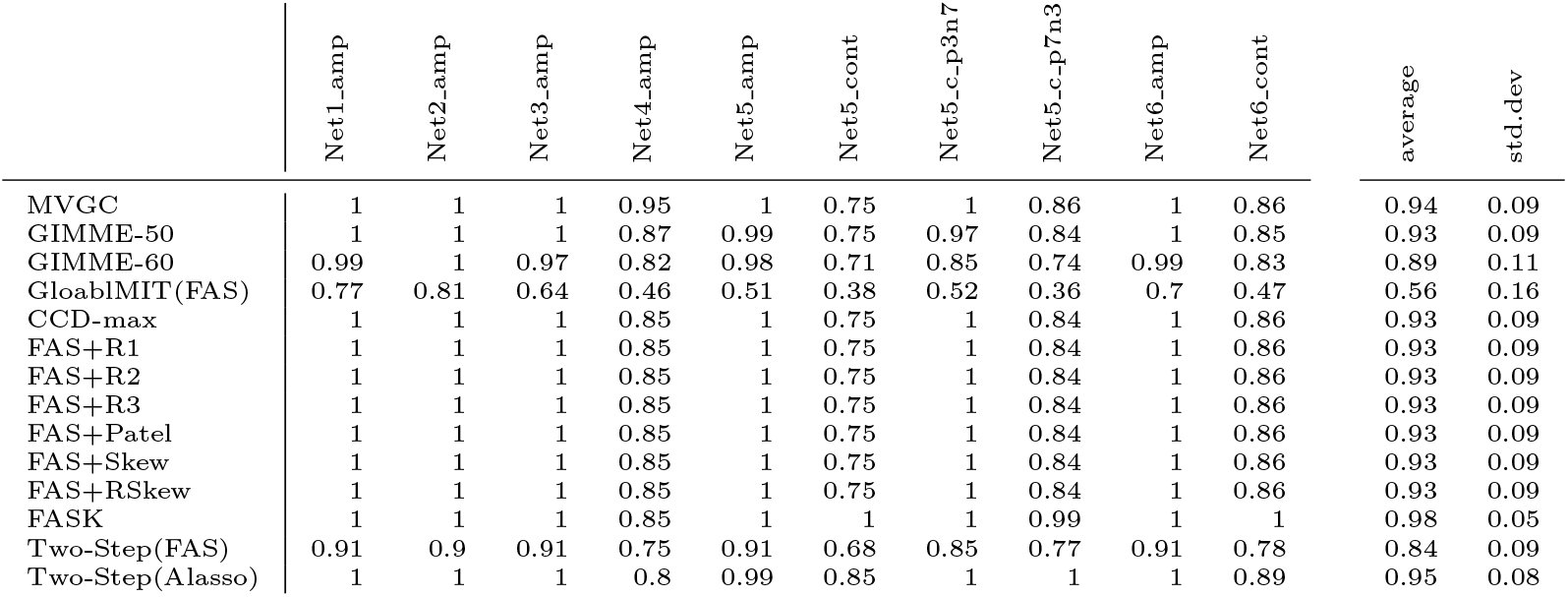
Adjacency Recall

**Table C3:**
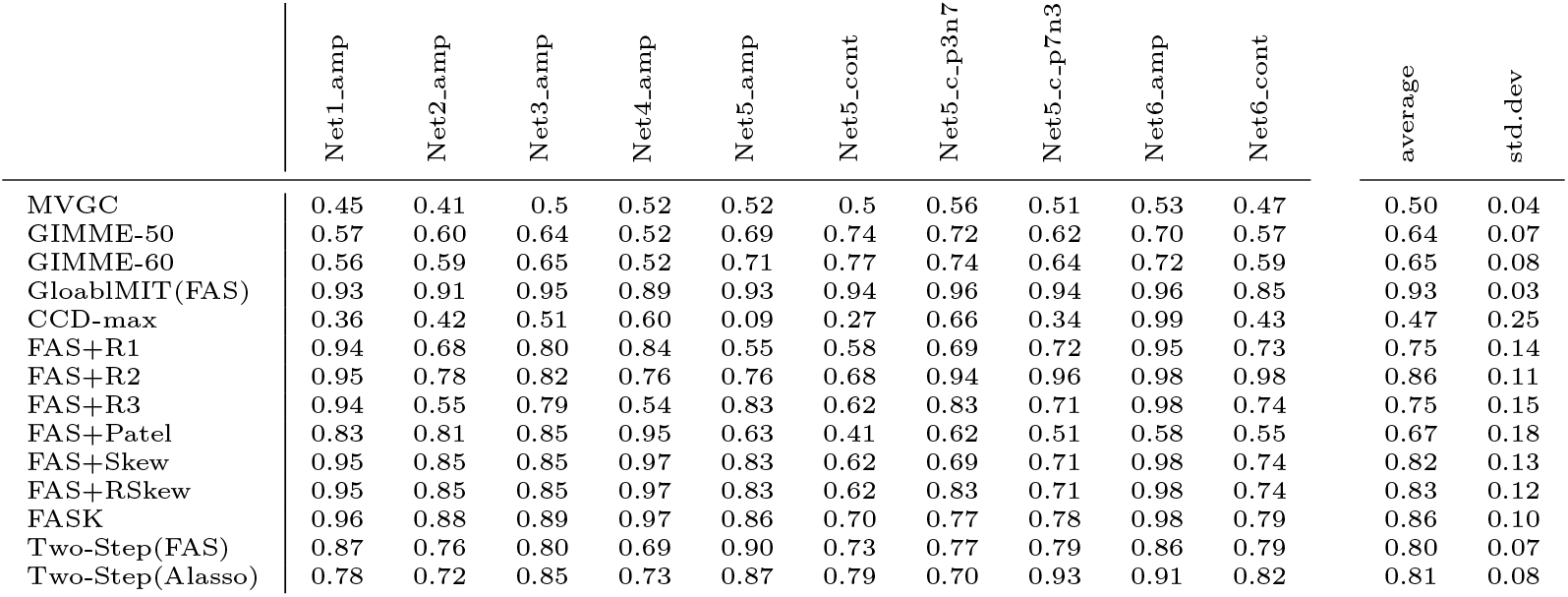
Orientation Precision

**Table C4:**
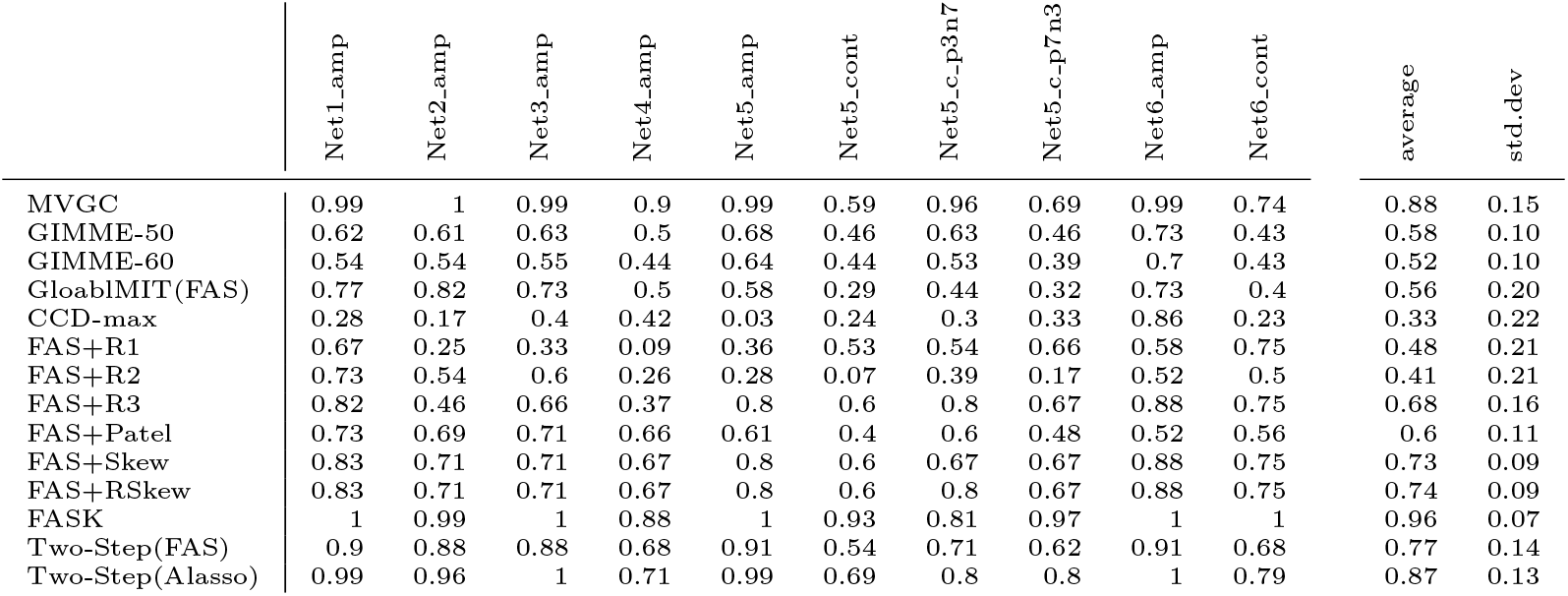
Orientation Recall

**Table C5:**
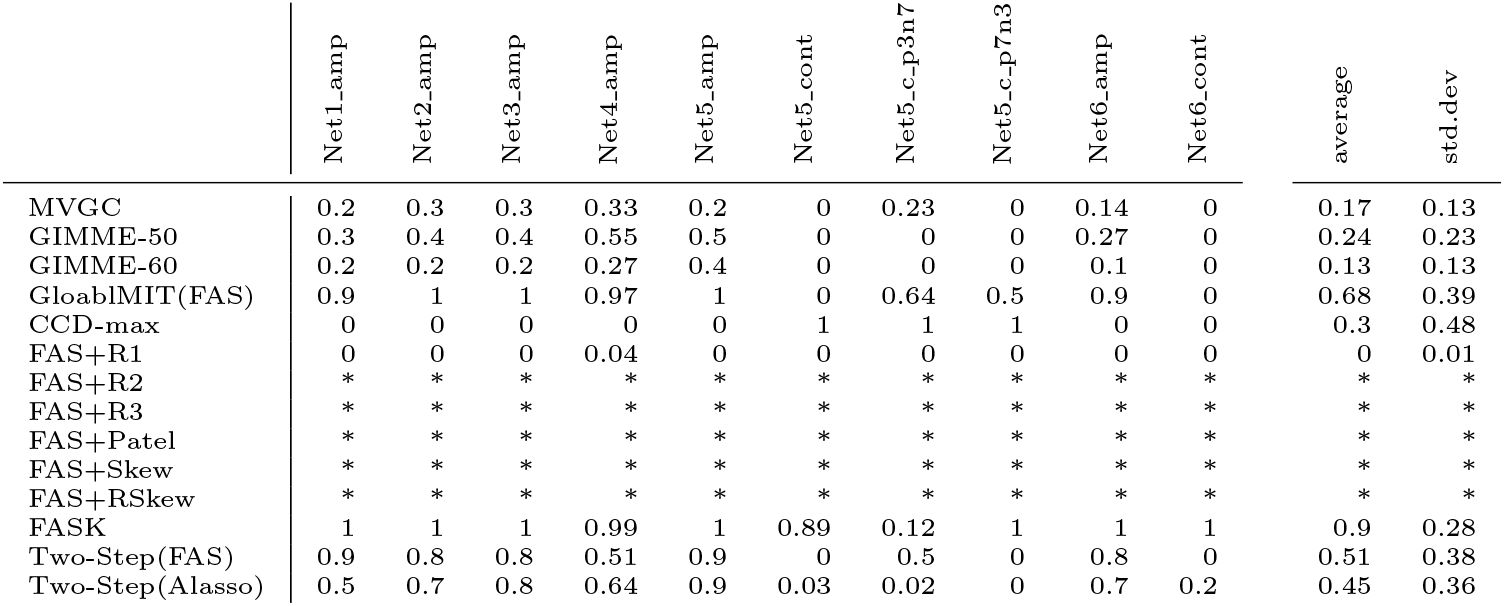
2-Cycle Precision. The (*) symbol indicates algorithms that cannot detect 2-cycles by design: R2, R3, Patel, Skew and RSkew.

**Table C6:** 2-Cycle Recall. The (*) symbol indicates algorithms that cannot detect 2-cycles by design:R2, R3, Patel, Skew and RSkew.

|  | Net1_amp | Net2_amp | Net3_amp | Net4_amp | Net5_amp | Net5_cont | Net5_c-p3n7 | Net5_c-p7n3 | Net6_amp | Net6_cont | average | std.dev |
| --- | --- | --- | --- | --- | --- | --- | --- | --- | --- | --- | --- | --- |
| MVGC | 1 | 1 | 1 | 1 | 1 | 0 | 0.88 | 0 | 1 | 0 | 0.69 | 0.48 |
| GIMME-50 | 0.3 | 0.21 | 0.3 | 0.2 | 0.6 | 0 | 0 | 0 | 0.3 | 0 | 0.18 | 0.18 |
| GIMME-60 | 0.2 | 0.09 | 0.1 | 0.1 | 0.4 | 0 | 0 | 0 | 0.2 | 0 | 0.09 | 0.12 |
| GloablMIT(FAS) | 1 | 0.93 | 1 | 0.9 | 1 | 0 | 0.12 | 0.13 | 1 | 0 | 0.60 | 0.47 |
| CCD-max | 0 | 0 | 0 | 0 | 0 | 0.02 | 0.02 | 0.08 | 0 | 0 | 0.01 | 0.03 |
| FAS+R1 | 0 | 0 | 0 | 0 | 0 | 0 | 0 | 0 | 0 | 0 | 0 | 0 |
| FAS+R2 | * | * | * | * | * | * | * | * | * | * | * | * |
| FAS+R3 | * | * | * | * | * | * | * | * | * | * | * | * |
| FAS+Patel | * | * | * | * | * | * | * | * | * | * | * | * |
| FAS+Skew | * | * | * | * | * | * | * | * | * | * | * | * |
| FAS+RSkew | * | * | * | * | * | * | * | * | * | * | * | * |
| FASK | 1 | 0.99 | 1 | 1 | 1 | 0.66 | 0.06 | 0.91 | 1 | 1 | 0.86 | 0.30 |
| Two-Step(FAS) | 0.9 | 0.88 | 0.8 | 0.7 | 0.9 | 0 | 0.52 | 0 | 0.9 | 0 | 0.57 | 0.41 |
| Two-Step(Alasso) | 1 | 0.93 | 1 | 0.9 | 1 | 0.03 | 0.02 | 0.02 | 1 | 0.2 | 0.60 | 0.46 |

**Table C7:** Running Time (seconds)

|  | Net1_amp | Net2_amp | Net3_amp | Net4_amp | Net5_amp | Net5_cont | Net5_c-p3n7 | Net5_c-p7n3 | Net6_amp | Net6_cont | average | std.dev |
| --- | --- | --- | --- | --- | --- | --- | --- | --- | --- | --- | --- | --- |
| MVGC | 0.13 | 0.11 | 0.12 | 0.39 | 0.10 | 0.10 | 0.11 | 0.10 | 0.23 | 0.17 | 0.15 | 0.09 |
| GIMME-50 | 44.0 | 54.8 | 49.4 | 446.5 | 42.6 | 28.8 | 34.6 | 57.8 | 199.2 | 206.8 | 116.5 | 133.64 |
| GIMME-60 | 101.2 | 72.7 | 87.7 | 766.7 | 36.1 | 27.0 | 30.9 | 27.5 | 108.0 | 140.2 | 139.8 | 223.78 |
| GloablMIT(FAS) | 0.41 | 0.67 | 0.68 | 3.37 | 0.58 | 0.42 | 0.58 | 0.52 | 1.54 | 1.64 | 1.04 | 0.93 |
| CCD-max | 0.00 | 0.00 | 0.00 | 0.01 | 0.00 | 0.00 | 0.00 | 0.00 | 0.00 | 0.00 | 0.002 | 0.00 |
| FAS+R1 | 0.11 | 0.13 | 0.12 | 0.44 | 0.10 | 0.10 | 0.11 | 0.10 | 0.18 | 0.20 | 0.16 | 0.10 |
| FAS+R2 | 0.74 | 1.26 | 1.14 | 6.20 | 0.90 | 0.69 | 0.91 | 0.77 | 1.67 | 2.15 | 1.64 | 1.67 |
| FAS+R3 | 0.10 | 0.11 | 0.11 | 0.32 | 0.09 | 0.09 | 0.09 | 0.08 | 0.14 | 0.15 | 0.13 | 0.07 |
| FAS+Patel | 0.01 | 0.02 | 0.02 | 0.12 | 0.01 | 0.01 | 0.01 | 0.01 | 0.03 | 0.02 | 0.03 | 0.03 |
| FAS+Skew | 0.01 | 0.01 | 0.01 | 0.11 | 0.01 | 0.01 | 0.01 | 0.01 | 0.02 | 0.01 | 0.02 | 0.03 |
| FAS+RSkew | 0.01 | 0.01 | 0.01 | 0.12 | 0.01 | 0.01 | 0.01 | 0.01 | 0.02 | 0.02 | 0.02 | 0.03 |
| FASK | 0.04 | 0.04 | 0.03 | 0.24 | 0.02 | 0.02 | 0.02 | 0.02 | 0.05 | 0.04 | 0.05 | 0.07 |
| Two-Step (FAS) | 22.45 | 59.00 | 28.86 | 54.80 | 9.50 | 6.08 | 16.95 | 6.92 | 11.83 | 9.47 | 22.59 | 19.47 |
| Two-Step(Alasso) | 33.78 | 79.31 | 59.74 | 110.59 | 32.39 | 29.56 | 27.97 | 20.20 | 70.95 | 30.42 | 49.49 | 29.45 |

### C2 Results for simple networks with higher degree cycles

Complete results for analysis shown in Figure 3, for *Network 7* to *9* and their amplifying and control variants. “amp” and “cont” after each network label indicates if the network has an amplifying (excitatory) or a control (inhibitory) cycle. Tables show for each algorithm tested the average results over 60 repetitions of ten concatenated datasets from the corresponding simulations, for precision and recall of adjacencies, orientations, and 2-cycle false positives; and running times in seconds. Average over all eight networks for each algorithm is shown to the right of each table, followed by the corresponding standard deviation. These averages are plotted in Figure 3.

**Table C8:**
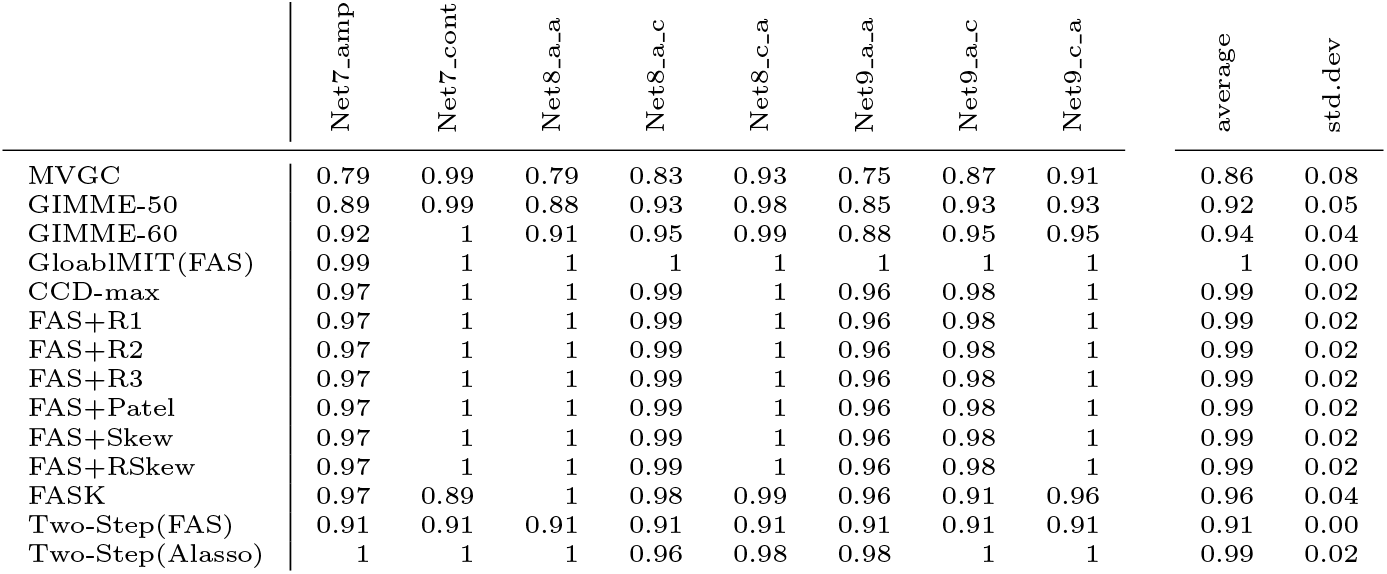
Adjacency Precision

**Table C9:**
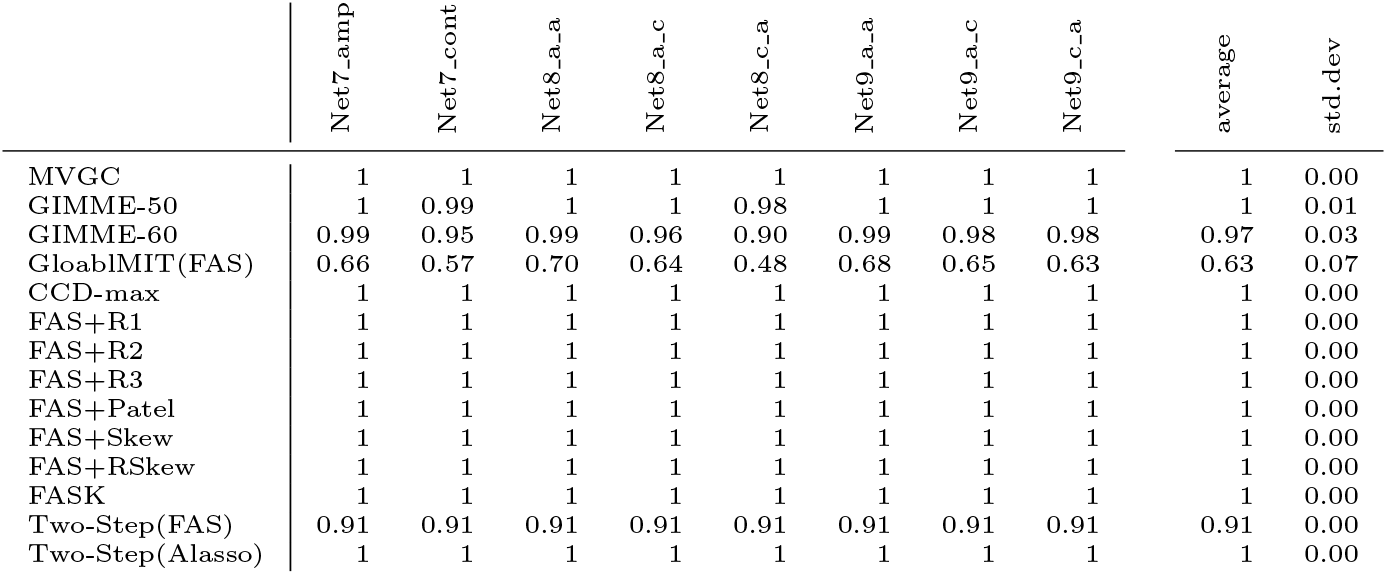
Adjacency Recall

**Table C10:** Orientation Precision

|  | Net7_amp | Net7_cont | Net8_a-a | Net8_a-c | Net8_c-a | Net9_a-a | Net9_a-c | Net9_c-a | average | std.dev |
| --- | --- | --- | --- | --- | --- | --- | --- | --- | --- | --- |
| MVGC | 0.42 | 0.5 | 0.4 | 0.45 | 0.47 | 0.4 | 0.47 | 0.48 | 0.46 | 0.03 |
| GIMME-50 | 0.64 | 0.6 | 0.6 | 0.66 | 0.57 | 0.6 | 0.61 | 0.64 | 0.62 | 0.03 |
| GIMME-60 | 0.66 | 0.61 | 0.6 | 0.69 | 0.57 | 0.7 | 0.63 | 0.65 | 0.63 | 0.04 |
| GloablMIT(FAS) | 0.76 | 0.89 | 0.8 | 0.8 | 0.81 | 0.8 | 0.84 | 0.84 | 0.81 | 0.04 |
| CCD-max | 0.84 | 0.59 | 0.7 | 0.7 | 0.64 | 0.8 | 0.89 | 0.8 | 0.74 | 0.11 |
| FAS+R1 | 1 | 0.97 | 1 | 0.97 | 0.98 | 1 | 0.96 | 0.88 | 0.97 | 0.04 |
| FAS+R2 | 0.97 | 1 | 1 | 0.99 | 0.99 | 1 | 0.99 | 1 | 0.99 | 0.01 |
| FAS+R3 | 0.97 | 0.98 | 1 | 0.94 | 0.98 | 1 | 0.92 | 0.9 | 0.96 | 0.03 |
| FAS+Patel | 0.55 | 0.41 | 0.5 | 0.53 | 0.56 | 0.4 | 0.43 | 0.48 | 0.48 | 0.06 |
| FAS+Skew | 0.96 | 1 | 1 | 0.99 | 1 | 1 | 0.98 | 1 | 0.99 | 0.02 |
| FAS+RSkew | 0.97 | 0.83 | 1 | 0.88 | 0.89 | 1 | 0.89 | 0.9 | 0.92 | 0.06 |
| FASK | 0.97 | 0.74 | 1 | 0.87 | 0.88 | 0.96 | 0.82 | 0.86 | 0.89 | 0.08 |
| Two-Step(FAS) | 0.87 | 0.75 | 0.9 | 0.78 | 0.84 | 0.8 | 0.74 | 0.74 | 0.80 | 0.06 |
| Two-Step(Alasso) | 0.76 | 0.69 | 0.7 | 0.7 | 0.72 | 0.7 | 0.69 | 0.74 | 0.71 | 0.03 |

**Table C11:**
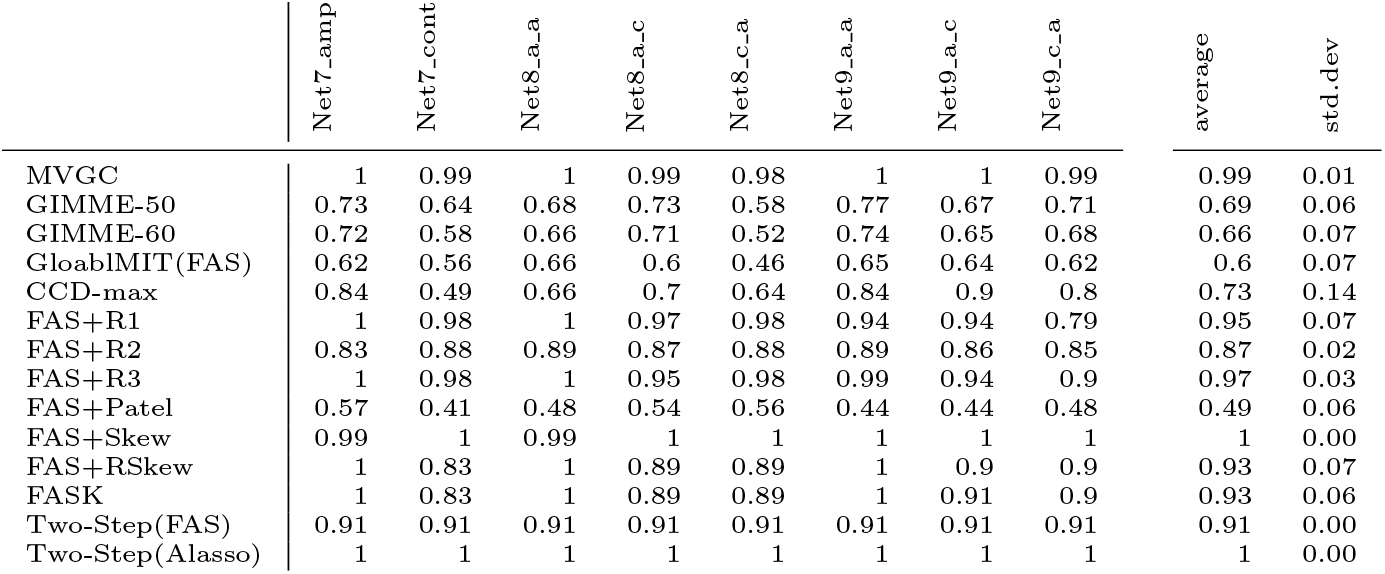
Orientation Recall

**Table C12:** 2-Cycle False Positives. The (*) symbol indicate algorithms that cannot detect 2-cycles by design: R2, R3, Patel, Skew and RSkew.

|  | Net7_amp | Net7_cont | Net8_a.a | Net8_a.c | Net8_c.a | Net9_a.a | Net9_a.c | Net9_c.a | average | std.dev |
| --- | --- | --- | --- | --- | --- | --- | --- | --- | --- | --- |
| MVGC | 6.57 | 5.88 | 9.27 | 8.92 | 9.15 | 10.65 | 9.95 | 9.6 | 8.75 | 1.66 |
| GIMME-50 | 0.22 | 0.37 | 0.32 | 0.37 | 0.2 | 0.37 | 0.33 | 0.43 | 0.33 | 0.08 |
| GIMME-60 | 0.03 | 0.05 | 0.03 | 0.05 | 0 | 0.12 | 0.08 | 0.03 | 0.05 | 0.04 |
| GloablMIT(FAS) | 0.92 | 0.35 | 1.07 | 0.92 | 0.82 | 1.58 | 1.12 | 1.07 | 0.98 | 0.34 |
| CCD-max | 0 | 0 | 0 | 0 | 0.02 | 0.02 | 0 | 0 | 0.01 | 0.01 |
| FAS+R1 | 0 | 0.23 | 0 | 0.23 | 0.13 | 0.35 | 0.33 | 1.07 | 0.29 | 0.34 |
| FAS+R2 | * | * | * | * | * | * | * | * | * | * |
| FAS+R3 | * | * | * | * | * | * | * | * | * | * |
| FAS+Patel | * | * | * | * | * | * | * | * | * | * |
| FAS+Skew | * | * | * | * | * | * | * | * | * | * |
| FAS+RSkew | * | * | * | * | * | * | * | * | * | * |
| FASK | 0.00 | 0.00 | 0.00 | 0.03 | 0.00 | 0.00 | 0.07 | 0.02 | 0.01 | 0.02 |
| Two-Step (FAS) | 0.59 | 1.48 | 0.86 | 1.6 | 0.96 | 1.56 | 2.26 | 2.18 | 1.44 | 0.60 |
| Two-Step(Alasso) | 2.03 | 2.72 | 3.53 | 3.7 | 3.58 | 4.67 | 4.62 | 3.48 | 3.54 | 0.88 |

**Table C13:** Running Time (seconds)

|  | Net7_amp | Net7_cont | Net8_a.a | Net8_a.c | Net8_c.a | Net9_a.a | Net9_a.c | Net9_c.a | average | std.dev |
| --- | --- | --- | --- | --- | --- | --- | --- | --- | --- | --- |
| MVGC | 0.13 | 0.13 | 0.19 | 0.19 | 0.18 | 0.27 | 0.22 | 0.23 | 0.19 | 0.05 |
| GIMME-50 | 156.8 | 159.4 | 166.6 | 129.6 | 120.4 | 220.1 | 201.2 | 173.2 | 165.9 | 33.27 |
| GIMME-60 | 53.8 | 49.6 | 120.0 | 118.9 | 112.1 | 209.1 | 189.6 | 186.6 | 130.0 | 60.75 |
| GloablMIT(FAS) | 0.89 | 0.65 | 1.38 | 1.36 | 1.29 | 2.22 | 1.89 | 1.88 | 1.45 | 0.53 |
| CCD-max | 0.00 | 0.00 | 0.00 | 0.00 | 0.00 | 0.00 | 0.00 | 0.00 | 0.00 | 0.00 |
| FAS+R1 | 0.15 | 0.13 | 0.22 | 0.32 | 0.21 | 0.28 | 0.26 | 0.25 | 0.23 | 0.06 |
| FAS+R2 | 1.49 | 1.09 | 2.34 | 3.04 | 2.30 | 3.63 | 3.16 | 2.95 | 2.50 | 0.87 |
| FAS+R3 | 0.12 | 0.11 | 0.23 | 0.17 | 0.17 | 0.21 | 0.19 | 0.19 | 0.17 | 0.04 |
| FAS+Patel | 0.02 | 0.01 | 0.05 | 0.03 | 0.03 | 0.05 | 0.03 | 0.03 | 0.03 | 0.01 |
| FAS+Skew | 0.01 | 0.01 | 0.04 | 0.02 | 0.02 | 0.04 | 0.02 | 0.02 | 0.02 | 0.01 |
| FAS+RSkew | 0.02 | 0.01 | 0.05 | 0.02 | 0.02 | 0.05 | 0.03 | 0.03 | 0.03 | 0.01 |
| FASK | 0.03 | 0.01 | 0.07 | 0.05 | 0.04 | 0.09 | 0.06 | 0.05 | 0.05 | 0.02 |
| Two-Step(FAS) | 9.54 | 10.23 | 11.61 | 14.23 | 13.53 | 17.03 | 19.16 | 21.96 | 14.66 | 4.40 |
| Two-Step(Alasso) | 19.96 | 18.71 | 30.18 | 36.24 | 28.87 | 42.01 | 33.68 | 42.11 | 31.47 | 8.91 |

### C3 Macaque-based networks (standard deviation)

**Table C14:** **Standard deviation** over 60 repetitions of 10 test datasets concatenated. Results for FASK and Two-Step with Alasso for the four different macaque-based networks in Section 2.2. Precision and recall for adjacencies (AP, AR), orientations (OP, OR), 2-cycles (2CP, 2CR), and running times in seconds (r.time).

| FASK (standard deviation) |  |  |  |  |  |  |  |
| --- | --- | --- | --- | --- | --- | --- | --- |
| Network | AP | AR | OP | OR | 2CP | 2CR | r.time(s) |
| SmallDegree | 0.03 | 0.03 | 0.05 | 0.05 | 0.09 | 0.14 | 1.98 |
| LongRange | 0.04 | 0.02 | 0.03 | 0.02 | 0.06 | 0.07 | 0.31 |
| LongRangeControl | 0.04 | 0.02 | 0.04 | 0.02 | 0.09 | 0.06 | 0.13 |
| Full | 0.04 | 0.00 | 0.03 | 0.00 | 0.02 | 0.01 | 11.42 |

| Two-Step(Alasso) (standard deviation) |  |  |  |  |  |  |  |
| --- | --- | --- | --- | --- | --- | --- | --- |
| Network | AP | AR | OP | OR | 2CP | 2CR | r.time(s) |
| SmallDegree | 0.05 | 0.04 | 0.03 | 0.05 | 0.03 | 0.16 | 10.71 |
| LongRange | 0.01 | 0.03 | 0.01 | 0.03 | 0.02 | 0.06 | 3.20 |
| LongRangeControl | 0.02 | 0.03 | 0.02 | 0.03 | 0.04 | 0.05 | 7.61 |
| Full | 0.02 | 0.01 | 0.02 | 0.01 | 0.03 | 0.03 | 26.83 |

### C4 Results for resting state medial temporal lobe data

#### C4.1 Left hemisphere results

Results for 23 repetitions of ten concatenated subjects, for seven regions of interest of the medial temporal lobe in the left hemisphere. For FASK and Two-Step(Alasso) we report the frequency of appearance in percentage, of each edge across the 23 repetitions. A value of 1.00 indicates that the edge shows in all 23 repetitions. A value of 0.04 indicates that the edge shows in only one of the repetitions. FASK was run with penalty *c* = 1 and α = 0.05; Two-Step(Alasso) was run with λ = 2 and threshold 0.10.

**Table C15:** FASK: medial temporal lobe edges ordered by frequency of appearance (left hemisphere). In bold the edges with frequency equal to 48% or more.

|  |  |  |  |  |  |  |  |  |  |  |  |
| --- | --- | --- | --- | --- | --- | --- | --- | --- | --- | --- | --- |
| **CA1** | → | **SUB** | **1.00** | **PHC** | → | **ERC** | **0.65** | BA36 | → | CA1 | 0.26 |
| **CA23DG** | → | **CA1** | **1.00** | **ERC** | → | **BA35** | **0.52** | ERC | → | SUB | 0.22 |
| **SUB** | → | **CA1** | **1.00** | **CA1** | → | **BA36** | **0.48** | CA1 | → | PHC | 0.17 |
| **BA36** | → | **BA35** | **1.00** | **SUB** | → | **PHC** | **0.48** | CA23DG | → | SUB | 0.17 |
| **CA1** | → | **CA23DG** | **0.96** | SUB | → | CA23DG | 0.43 | ERC | → | PHC | 0.17 |
| **SUB** | → | **ERC** | **0.91** | BA35 | → | PHC | 0.43 | CA1 | → | BA35 | 0.09 |
| **BA35** | → | **BA36** | **0.91** | SUB | → | BA35 | 0.26 | CA23DG | → | ERC | 0.09 |
| **PHC** | → | **SUB** | **0.83** | BA35 | → | SUB | 0.26 | PHC | → | CA1 | 0.09 |
| **PHC** | → | **BA35** | **0.78** | BA35 | → | ERC | 0.26 | BA35 | → | CA1 | 0.04 |

**Table C16:**
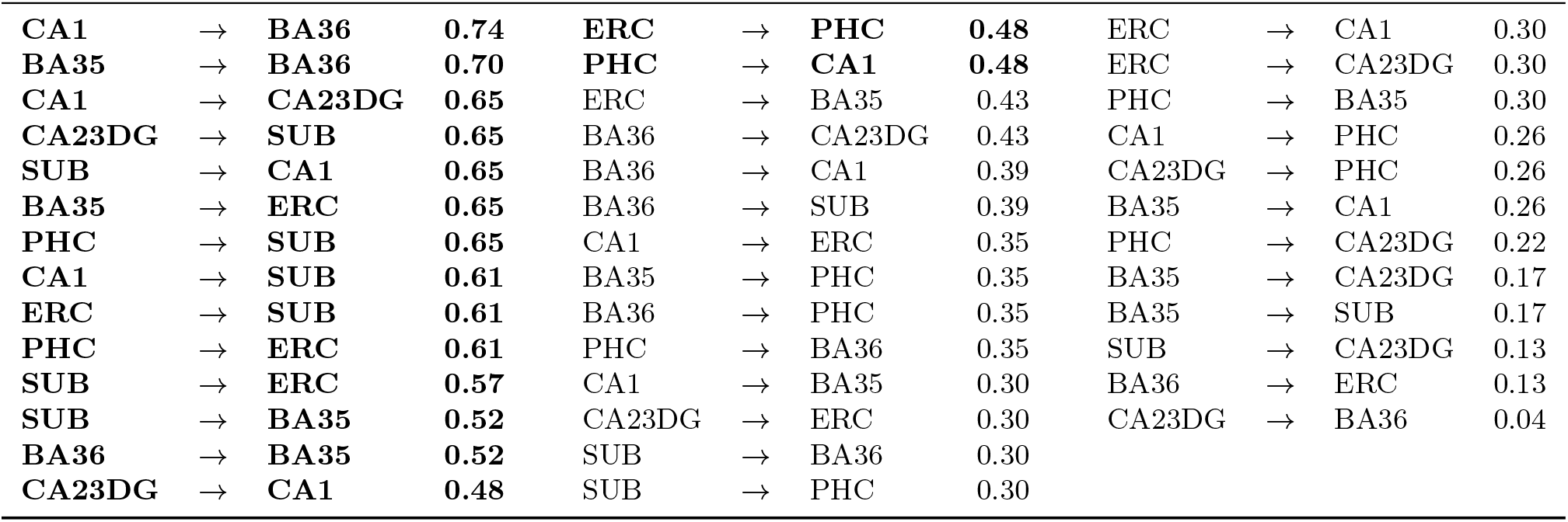
Two-Step(Alasso): medial temporal lobe edges ordered by frequency of appearance (left hemisphere). In bold the edges with frequency equal to 48% or more.

#### C4.2 Right hemisphere results

Results for 23 repetitions of ten concatenated subjects, for seven regions of interest of the medial temporal lobe in the right hemisphere. Figure C1 shows graphs depicting robust directed edges that appear in 48% or more of the 23 repetitions, for FASK and Two-Step(Alasso). For FASK and Two-Step(Alasso) we also report the frequency of appearance in percentage, of each edge across the 23 repetitions. A value of 1.00 indicates that the edge shows in all 23 repetitions. A value of 0.04 indicates that the edge shows in only one of the repetitions. FASK was run with penalty *c* = 1 and α = 0.05; Two-Step(Alasso) was run with λ = 2 and threshold 0.10.

**Figure C1.**
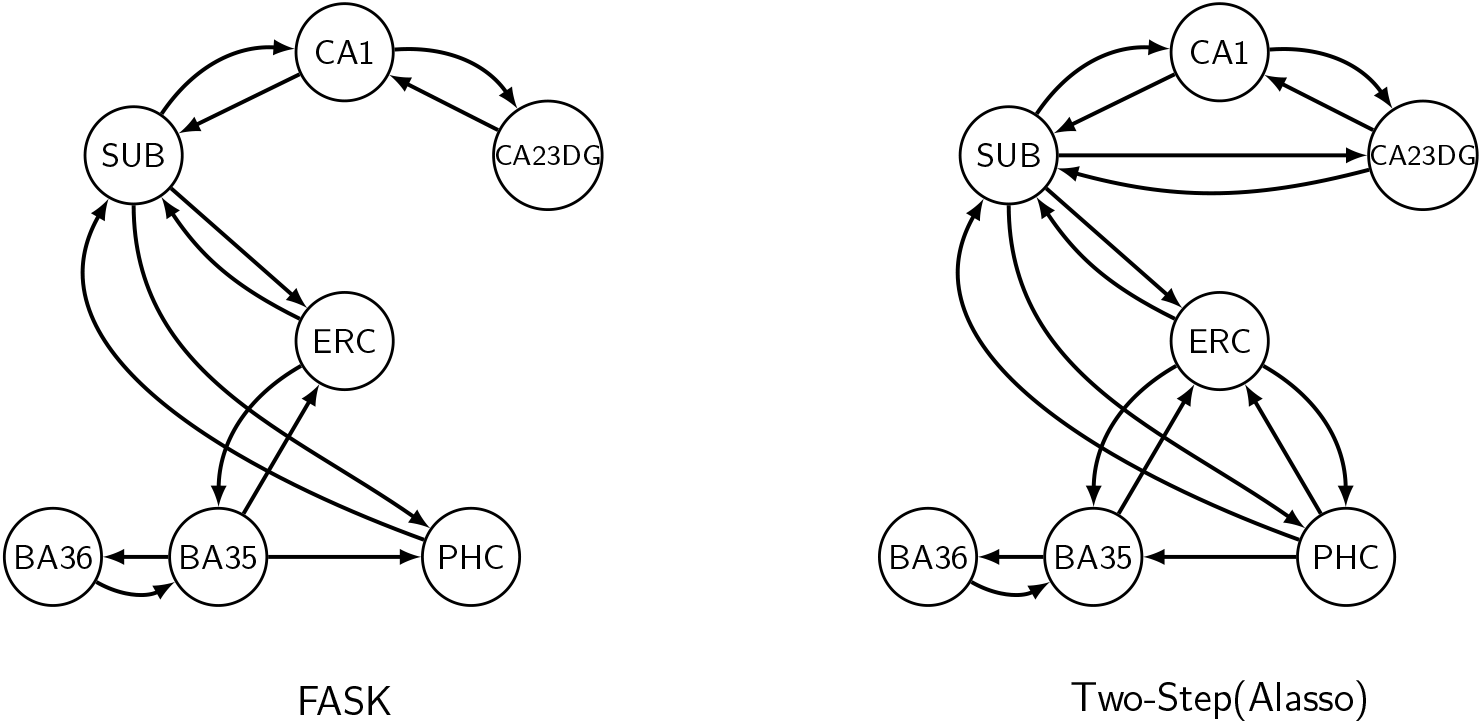
Comparison of the most robust edges estimated by FASK and Two-Step(Alasso) in 23 repetitions of ten concatenated subjects for seven regions of interest from the right hemisphere medial temporal lobe. A directed edge is depicted if it is estimated in 48% or more of the 23 repetitions of ten subjects concatenated. Regions of interest include: Cornu Ammonis 1 (CA1); CA2, CA3 and dentate gyrus together (CA23DG); entorhinal cortex (ERC); perirhinal cortex divided in Brodmann areas (BA35 and BA36); and parahippocampal cortex (PHC).

**Table C17:** FASK: medial temporal lobe edges ordered by frequency of appearance (right hemisphere). In bold the edges with frequency equal to 48% or more.

|  |  |  |  |  |  |  |  |  |  |  |  |
| --- | --- | --- | --- | --- | --- | --- | --- | --- | --- | --- | --- |
| CA1 | → | CA23DG | 1.00 | BA35 | → | ERC | 0.87 | BA36 | → | CA1 | 0.17 |
| CA1 | → | SUB | 1.00 | BA35 | → | PHC | 0.87 | SUB | → | CA23DG | 0.13 |
| CA23DG | → | CA1 | 1.00 | PHC | → | SUB | 0.83 | BA36 | → | CA23DG | 0.09 |
| SUB | → | ERC | 1.00 | SUB | → | PHC | 0.70 | CA1 | → | BA36 | 0.04 |
| ERC | → | SUB | 1.00 | CA1 | → | PHC | 0.39 | CA23DG | → | BA36 | 0.04 |
| BA36 | → | BA35 | 1.00 | PHC | → | CA1 | 0.39 | BA35 | → | SUB | 0.04 |
| SUB | → | CA1 | 0.96 | CA23DG | → | SUB | 0.26 | PHC | → | ERC | 0.04 |
| ERC | → | BA35 | 0.96 | PHC | → | BA35 | 0.26 |  |  |  |  |
| BA35 | → | BA36 | 0.96 | ERC | → | PHC | 0.17 |  |  |  |  |

**Table C18:** Two-Step(Alasso): medial temporal lobe edges by frequency of appearance (right hemisphere). In bold the edges with frequency equal to 48% or more.

|  |  |  |  |  |  |  |  |  |  |  |  |
| --- | --- | --- | --- | --- | --- | --- | --- | --- | --- | --- | --- |
| CA1 | → | SUB | 0.78 | CA23DG | → | SUB | 0.48 | SUB | → | BA36 | 0.17 |
| CA1 | → | CA23DG | 0.74 | SUB | → | PHC | 0.48 | BA36 | → | CA23DG | 0.17 |
| CA23DG | → | CA1 | 0.74 | ERC | → | PHC | 0.48 | BA36 | → | PHC | 0.17 |
| ERC | → | SUB | 0.70 | CA1 | → | PHC | 0.39 | CA23DG | → | ERC | 0.13 |
| SUB | → | CA1 | 0.65 | CA23DG | → | BA36 | 0.35 | ERC | → | CA23DG | 0.13 |
| BA35 | → | BA36 | 0.65 | BA35 | → | PHC | 0.35 | CA1 | → | BA35 | 0.09 |
| PHC | → | ERC | 0.65 | PHC | → | CA1 | 0.35 | BA35 | → | CA23DG | 0.09 |
| SUB | → | ERC | 0.61 | SUB | → | BA35 | 0.30 | BA35 | → | SUB | 0.09 |
| BA35 | → | ERC | 0.61 | ERC | → | BA36 | 0.26 | BA36 | → | CA1 | 0.09 |
| SUB | → | CA23DG | 0.57 | BA36 | → | ERC | 0.26 | ERC | → | CA1 | 0.04 |
| BA36 | → | BA35 | 0.57 | CA1 | → | BA36 | 0.22 | BA35 | → | CA1 | 0.04 |
| PHC | → | SUB | 0.57 | CA23DG | → | PHC | 0.22 | BA36 | → | SUB | 0.04 |
| ERC | → | BA35 | 0.52 | PHC | → | BA36 | 0.22 |  |  |  |  |
| PHC | → | BA35 | 0.52 | CA1 | → | ERC | 0.17 |  |  |  |  |

#### C4.3 2-cycles frequency of appearance

For FASK and Two-Step(Alasso) we report 2-cycles by frequency of appearance in percentage, across the 23 repetitions of ten concatenated subjects, for seven regions of interest from the medial temporal lobe for the left and right hemisphere correspondingly. A value of 1.00 indicates that the 2-cycle appears in all 23 repetitions. A value of 0.04 indicates that the 2-cycle appears in only one repetition. FASK was run with penalty *c* = 1 and α = 0.05; Two-Step(Alasso) was run with λ = 2 and threshold 0.10.

**Table C19:** FASK: 2-cycles by frequency of appearance for medial temporal lobe regions of interest.

| left hemisphere |  |  |  | right hemisphere |  |  |  |
| --- | --- | --- | --- | --- | --- | --- | --- |
| CA1 | ⇌ | SUB | 1.00 | CA1 | ⇌ | CA23DG | 1.00 |
| CA1 | ⇌ | CA23DG | 0.96 | SUB | ⇌ | ERC | 1.00 |
| BA35 | ⇌ | BA36 | 0.91 | CA1 | ⇌ | SUB | 0.96 |
| SUB | ⇌ | PHC | 0.43 | BA35 | ⇌ | BA36 | 0.96 |
| SUB | ⇌ | BA35 | 0.26 | ERC | ⇌ | BA35 | 0.83 |
| BA35 | ⇌ | PHC | 0.22 | SUB | ⇌ | PHC | 0.70 |
| CA1 | ⇌ | BA36 | 0.13 | CA1 | ⇌ | PHC | 0.22 |
| SUB | ⇌ | ERC | 0.13 | BA35 | ⇌ | PHC | 0.13 |
| ERC | ⇌ | BA35 | 0.04 | CA23DG | ⇌ | BA36 | 0.04 |
|  |  |  |  | ERC | ⇌ | PHC | 0.04 |

**Table C20:** Two-Step(Alasso): 2-cycles by frequency of appearance for medial temporal lobe regions of interest.

| left hemisphere |  |  |  | right hemisphere |  |  |  |
| --- | --- | --- | --- | --- | --- | --- | --- |
| CA1 | ⇌ | BA36 | 0.30 | CA1 | ⇌ | CA23DG | 0.48 |
| ERC | ⇌ | PHC | 0.30 | ERC | ⇌ | PHC | 0.48 |
| CA1 | ⇌ | SUB | 0.26 | CA1 | ⇌ | SUB | 0.43 |
| CA1 | ⇌ | ERC | 0.26 | CA23DG | ⇌ | SUB | 0.35 |
| BA36 | ⇌ | PHC | 0.26 | SUB | ⇌ | ERC | 0.30 |
| SUB | ⇌ | ERC | 0.22 | SUB | ⇌ | PHC | 0.22 |
| ERC | ⇌ | BA35 | 0.22 | BA35 | ⇌ | BA36 | 0.22 |
| BA35 | ⇌ | BA36 | 0.22 | ERC | ⇌ | BA35 | 0.17 |
| CA1 | ⇌ | BA35 | 0.17 | ERC | ⇌ | BA36 | 0.17 |
| CA23DG | ⇌ | PHC | 0.17 | CA23DG | ⇌ | BA36 | 0.13 |
| SUB | ⇌ | BA36 | 0.17 | BA36 | ⇌ | PHC | 0.13 |
| CA1 | ⇌ | CA23DG | 0.13 | CA23DG | ⇌ | ERC | 0.09 |
| CA1 | ⇌ | PHC | 0.13 | CA1 | ⇌ | ERC | 0.04 |
| CA23DG | ⇌ | SUB | 0.13 | CA1 | ⇌ | PHC | 0.04 |
| CA23DG | ⇌ | ERC | 0.13 | BA35 | ⇌ | PHC | 0.04 |
| SUB | ⇌ | BA35 | 0.13 | | | | |
| SUB | ⇌ | PHC | 0.13 | | | | |
| BA35 | ⇌ | PHC | 0.09 | | | | |
| CA23DG | ⇌ | BA36 | 0.04 | | | | |

#### C4.4 Individual datasets results for medial temporal lobe data

Results for 23 individual datasets, for seven regions of interest of the medial temporal lobe in the left and right hemisphere correspondingly. For FASK and Two-Step(Alasso) we report the frequency of appearance in percentage, of each edge across the 23 individual datasets. A value of 1.00 indicates that the edge shows in all 23 subjects. A value of 0.04 indicates that the edge shows in only one of the subjects. We also report frequency of appearance for 2-cycles. FASK was run with penalty *c* = 1 and α = 0.10; Two-Step(Alasso) was run with λ = 2 and threshold 0.10.

**Table C21:** FASK: medial temporal lobe edges by frequency of appearance (individual datasets) (left hemisphere).

|  |  |  |  |  |  |  |  |  |  |  |  |
| --- | --- | --- | --- | --- | --- | --- | --- | --- | --- | --- | --- |
| CA1 | → | SUB | 0.57 | ERC | → | PHC | 0.22 | BA35 | → | CA23DG | 0.09 |
| SUB | → | CA1 | 0.57 | BA36 | → | PHC | 0.22 | BA35 | → | SUB | 0.09 |
| BA35 | → | BA36 | 0.57 | PHC | → | BA36 | 0.22 | BA36 | → | CA23DG | 0.09 |
| CA23DG | → | CA1 | 0.52 | CA23DG | → | ERC | 0.17 | BA36 | → | ERC | 0.09 |
| CA1 | → | CA23DG | 0.43 | BA35 | → | PHC | 0.17 | CA1 | → | ERC | 0.04 |
| BA36 | → | BA35 | 0.39 | BA36 | → | CA1 | 0.17 | CA1 | → | PHC | 0.04 |
| SUB | → | ERC | 0.35 | BA36 | → | SUB | 0.17 | CA23DG | → | SUB | 0.04 |
| ERC | → | SUB | 0.35 | PHC | → | BA35 | 0.17 | CA23DG | → | PHC | 0.04 |
| PHC | → | ERC | 0.30 | CA1 | → | BA35 | 0.13 | SUB | → | CA23DG | 0.04 |
| ERC | → | CA23DG | 0.26 | CA1 | → | BA36 | 0.13 | SUB | → | BA35 | 0.04 |
| BA35 | → | ERC | 0.26 | ERC | → | BA36 | 0.13 | SUB | → | BA36 | 0.04 |
| PHC | → | SUB | 0.26 | CA23DG | → | BA35 | 0.09 | BA35 | → | CA1 | 0.04 |
| SUB | → | PHC | 0.22 | CA23DG | → | BA36 | 0.09 | PHC | → | CA1 | 0.04 |
| ERC | → | BA35 | 0.22 | ERC | → | CA1 | 0.09 | PHC | → | CA23DG | 0.04 |

**Table C22:** Two-Step(Alasso): medial temporal lobe edges by frequency of appearance (individual datasets) (left hemisphere).

|  |  |  |  |  |  |  |  |  |  |  |  |
| --- | --- | --- | --- | --- | --- | --- | --- | --- | --- | --- | --- |
| CA1 | → | CA23DG | 0.57 | SUB | → | CA23DG | 0.39 | BA35 | → | CA23DG | 0.30 |
| SUB | → | CA1 | 0.57 | ERC | → | BA36 | 0.39 | BA36 | → | CA1 | 0.30 |
| ERC | → | CA23DG | 0.57 | BA36 | → | ERC | 0.39 | CA1 | → | PHC | 0.26 |
| ERC | → | SUB | 0.57 | CA1 | → | BA35 | 0.35 | ERC | → | PHC | 0.26 |
| BA35 | → | BA36 | 0.57 | CA23DG | → | CA1 | 0.35 | BA35 | → | PHC | 0.26 |
| PHC | → | BA35 | 0.57 | CA23DG | → | BA35 | 0.35 | BA36 | → | PHC | 0.26 |
| BA36 | → | BA35 | 0.52 | SUB | → | ERC | 0.35 | PHC | → | CA1 | 0.26 |
| PHC | → | SUB | 0.52 | SUB | → | BA35 | 0.35 | PHC | → | CA23DG | 0.26 |
| CA1 | → | SUB | 0.48 | BA36 | → | SUB | 0.35 | PHC | → | BA36 | 0.26 |
| CA1 | → | BA36 | 0.48 | CA23DG | → | ERC | 0.30 | CA23DG | → | SUB | 0.17 |
| CA1 | → | ERC | 0.43 | CA23DG | → | BA36 | 0.30 | CA23DG | → | PHC | 0.13 |
| SUB | → | PHC | 0.43 | SUB | → | BA36 | 0.30 | BA35 | → | ERC | 0.13 |
| ERC | → | BA35 | 0.43 | ERC | → | CA1 | 0.30 | BA36 | → | CA23DG | 0.13 |
| PHC | → | ERC | 0.43 | BA35 | → | CA1 | 0.30 | BA35 | → | SUB | 0.04 |

**Table C23:** FASK: medial temporal lobe edges by frequency of appearance (individual datasets) (right hemisphere).

|  |  |  |  |  |  |  |  |  |  |  |  |
| --- | --- | --- | --- | --- | --- | --- | --- | --- | --- | --- | --- |
| CA23DG | → | CA1 | 0.78 | CA23DG | → | SUB | 0.17 | PHC | → | CA23DG | 0.09 |
| CA1 | → | SUB | 0.65 | SUB | → | CA23DG | 0.17 | CA1 | → | ERC | 0.04 |
| CA1 | → | CA23DG | 0.61 | BA36 | → | CA23DG | 0.17 | CA1 | → | BA35 | 0.04 |
| BA36 | → | BA35 | 0.52 | BA36 | → | ERC | 0.17 | CA23DG | → | ERC | 0.04 |
| ERC | → | SUB | 0.48 | BA36 | → | PHC | 0.17 | CA23DG | → | PHC | 0.04 |
| ERC | → | BA35 | 0.48 | CA23DG | → | BA36 | 0.13 | SUB | → | BA35 | 0.04 |
| BA35 | → | PHC | 0.43 | PHC | → | ERC | 0.13 | ERC | → | CA23DG | 0.04 |
| SUB | → | ERC | 0.39 | PHC | → | BA35 | 0.13 | ERC | → | BA36 | 0.04 |
| BA35 | → | ERC | 0.39 | PHC | → | BA36 | 0.13 | BA35 | → | CA1 | 0.04 |
| BA35 | → | BA36 | 0.35 | CA23DG | → | BA35 | 0.09 | BA35 | → | CA23DG | 0.04 |
| SUB | → | CA1 | 0.30 | SUB | → | PHC | 0.09 | BA35 | → | SUB | 0.04 |
| ERC | → | PHC | 0.30 | BA36 | → | CA1 | 0.09 | BA36 | → | SUB | 0.04 |
| PHC | → | SUB | 0.26 | PHC | → | CA1 | 0.09 |  |  |  |  |

**Table C24:** Two-Step(Alasso): medial temporal lobe edges by frequency of appearance (individual datasets) (right hemisphere).

|  |  |  |  |  |  |  |  |  |  |  |  |
| --- | --- | --- | --- | --- | --- | --- | --- | --- | --- | --- | --- |
| BA36 | → | BA35 | 0.65 | ERC | → | SUB | 0.39 | ERC | → | CA23DG | 0.26 |
| SUB | → | ERC | 0.61 | BA35 | → | CA23DG | 0.39 | ERC | → | PHC | 0.26 |
| CA1 | → | CA23DG | 0.52 | BA36 | → | CA1 | 0.39 | BA36 | → | ERC | 0.26 |
| CA23DG | → | CA1 | 0.52 | CA1 | → | ERC | 0.35 | BA36 | → | PHC | 0.26 |
| BA35 | → | ERC | 0.52 | CA1 | → | BA35 | 0.35 | PHC | → | CA1 | 0.26 |
| BA35 | → | PHC | 0.52 | CA1 | → | PHC | 0.35 | PHC | → | ERC | 0.26 |
| CA23DG | → | SUB | 0.48 | CA23DG | → | BA35 | 0.35 | SUB | → | BA35 | 0.22 |
| CA23DG | → | ERC | 0.48 | PHC | → | SUB | 0.35 | BA35 | → | CA1 | 0.22 |
| SUB | → | CA1 | 0.48 | PHC | → | BA35 | 0.35 | PHC | → | CA23DG | 0.22 |
| ERC | → | BA35 | 0.48 | PHC | → | BA36 | 0.35 | BA36 | → | SUB | 0.17 |
| BA35 | → | BA36 | 0.48 | CA23DG | → | PHC | 0.30 | SUB | → | BA36 | 0.13 |
| CA1 | → | SUB | 0.43 | ERC | → | BA36 | 0.30 | ERC | → | CA1 | 0.13 |
| SUB | → | PHC | 0.43 | CA23DG | → | BA36 | 0.26 | BA35 | → | SUB | 0.13 |
| BA36 | → | CA23DG | 0.43 | SUB | → | CA23DG | 0.26 | CA1 | → | BA36 | 0.09 |

**Table C25:**
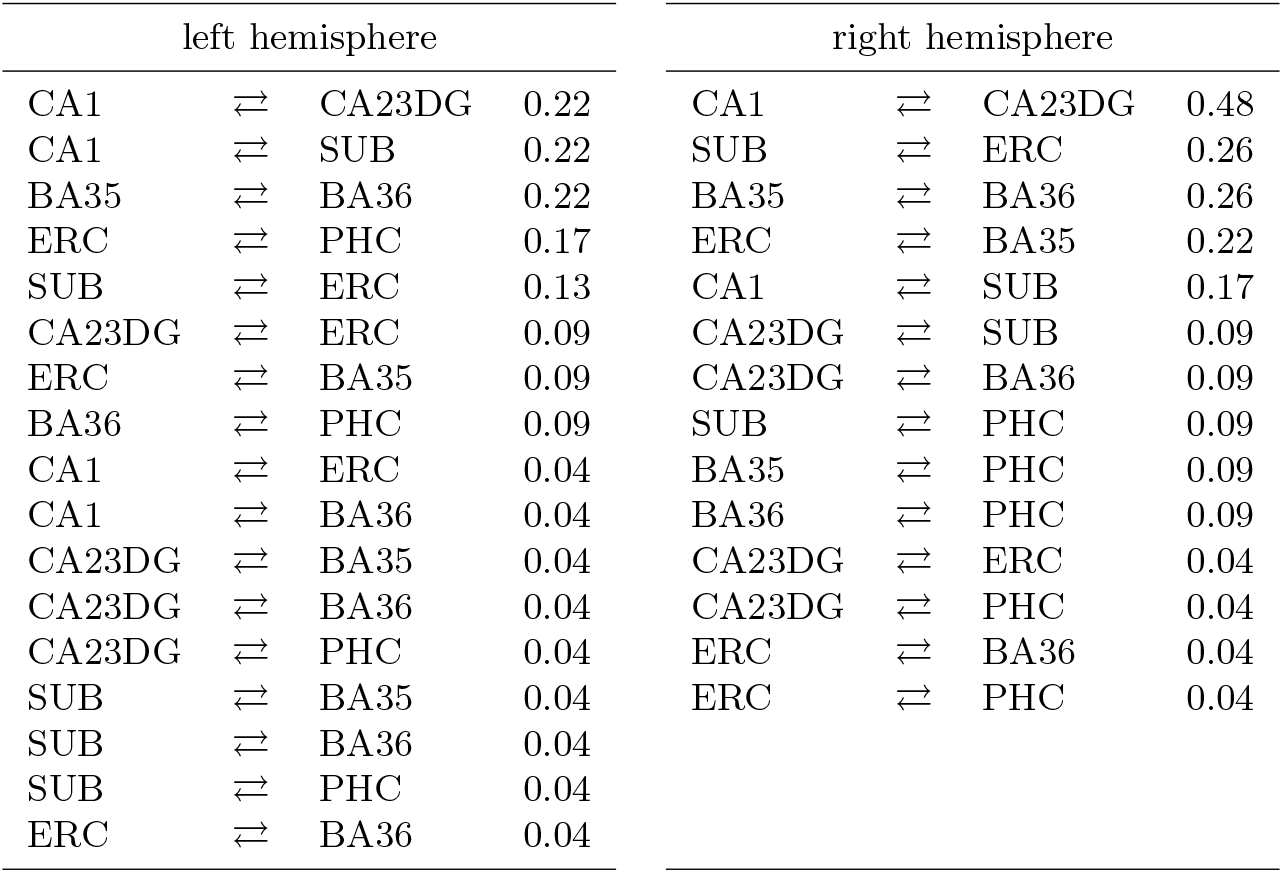
FASK: 2-cycles by frequency of appearance for medial temporal lobe regions of interest (individual data).

| left hemisphere |  |  |  | right hemisphere |  |  |  |
| --- | --- | --- | --- | --- | --- | --- | --- |
| CA1 | ⇌ | CA23DG | 0.22 | CA1 | ⇌ | CA23DG | 0.48 |
| CA1 | ⇌ | SUB | 0.22 | SUB | ⇌ | ERC | 0.26 |
| BA35 | ⇌ | BA36 | 0.22 | BA35 | ⇌ | BA36 | 0.26 |
| ERC | ⇌ | PHC | 0.17 | ERC | ⇌ | BA35 | 0.22 |
| SUB | ⇌ | ERC | 0.13 | CA1 | ⇌ | SUB | 0.17 |
| CA23DG | ⇌ | ERC | 0.09 | CA23DG | ⇌ | SUB | 0.09 |
| ERC | ⇌ | BA35 | 0.09 | CA23DG | ⇌ | BA36 | 0.09 |
| BA36 | ⇌ | PHC | 0.09 | SUB | ⇌ | PHC | 0.09 |
| CA1 | ⇌ | ERC | 0.04 | BA35 | ⇌ | PHC | 0.09 |
| CA1 | ⇌ | BA36 | 0.04 | BA36 | ⇌ | PHC | 0.09 |
| CA23DG | ⇌ | BA35 | 0.04 | CA23DG | ⇌ | ERC | 0.04 |
| CA23DG | ⇌ | BA36 | 0.04 | CA23DG | ⇌ | PHC | 0.04 |
| CA23DG | ⇌ | PHC | 0.04 | ERC | ⇌ | BA36 | 0.04 |
| SUB | ⇌ | BA35 | 0.04 | ERC | ⇌ | PHC | 0.04 |
| SUB | ⇌ | BA36 | 0.04 | | | | |
| SUB | ⇌ | PHC | 0.04 | | | | |
| ERC | ⇌ | BA36 | 0.04 | | | | |

**Table C26:** Two-Step(Alasso): 2-cycles by frequency of appearance for medial temporal lobe regions of interest (individual data).

| left hemisphere |  |  |  | right hemisphere |  |  |  |
| --- | --- | --- | --- | --- | --- | --- | --- |
| CA1 | ⇌ | BA35 | 0.17 | BA35 | ⇌ | BA36 | 0.26 |
| CA1 | ⇌ | BA36 | 0.17 | CA23DG | ⇌ | ERC | 0.17 |
| CA23DG | ⇌ | ERC | 0.17 | CA23DG | ⇌ | BA35 | 0.17 |
| SUB | ⇌ | ERC | 0.17 | SUB | ⇌ | ERC | 0.13 |
| ERC | ⇌ | BA36 | 0.17 | SUB | ⇌ | PHC | 0.13 |
| BA35 | ⇌ | BA36 | 0.17 | ERC | ⇌ | BA35 | 0.13 |
| CA1 | ⇌ | SUB | 0.13 | CA1 | ⇌ | CA23DG | 0.09 |
| CA1 | ⇌ | ERC | 0.13 | CA1 | ⇌ | ERC | 0.09 |
| CA23DG | ⇌ | BA35 | 0.13 | CA1 | ⇌ | BA35 | 0.09 |
| SUB | ⇌ | PHC | 0.13 | CA23DG | ⇌ | SUB | 0.09 |
| CA1 | ⇌ | PHC | 0.09 | CA23DG | ⇌ | PHC | 0.09 |
| SUB | ⇌ | BA36 | 0.09 | SUB | ⇌ | BA35 | 0.09 |
| CA1 | ⇌ | CA23DG | 0.04 | BA35 | ⇌ | PHC | 0.09 |
| CA23DG | ⇌ | SUB | 0.04 | CA1 | ⇌ | SUB | 0.04 |
| CA23DG | ⇌ | PHC | 0.04 | CA1 | ⇌ | BA36 | 0.04 |
| ERC | ⇌ | BA35 | 0.04 | CA1 | ⇌ | PHC | 0.04 |
| BA35 | ⇌ | PHC | 0.04 | CA23DG | ⇌ | BA36 | 0.04 |
| | | | | ERC | ⇌ | BA36 | 0.04 |
| | | | | BA36 | ⇌ | PHC | 0.04 |

### C5 Results for rhyming task individual data

FASK and Two-Step(Alasso) were run on 9 individual standardized datasets (160 datapoints) for eight regions of interest in the cortex and one Input variable modeling the dynamics of the visual stimuli presentation in the rhyming task. We report frequency of appearance in percentage, of individual edges across the 9 subjects. We also report 2-cycles frequency of appearance. A value of 1.00 indicates the edge appears in all the 9 subjects. A value of 0.11 indicates the edge appears in just one subject. FASK was run with a penalty discount of *c* = 1 and α = 0.10 to account for the reduction in sample size relative to the concatenated data (1,440 datapoints). Two-Step(Alasso) was run with = 2 and threshold for the absolute values of the **B** matrix of 0.10.

**Table C27:** FASK: rhyming task data edges ordered by frequency of appearance (individual datasets).

|  |  |  |  |  |  |  |  |  |  |  |  |
| --- | --- | --- | --- | --- | --- | --- | --- | --- | --- | --- | --- |
| LOCC | → | ROCC | 0.89 | LIPL | → | LIFG | 0.22 | RACC | → | LIPL | 0.11 |
| LACC | → | RACC | 0.78 | RIFG | → | LIFG | 0.22 | RACC | → | RIPL | 0.11 |
| ROCC | → | LOCC | 0.67 | RIFG | → | RACC | 0.22 | RACC | → | Input | 0.11 |
| LIPL | → | RIPL | 0.56 | LOCC | → | LIFG | 0.11 | RIFG | → | RIPL | 0.11 |
| LIFG | → | LIPL | 0.44 | LOCC | → | LIPL | 0.11 | RIFG | → | Input | 0.11 |
| LIFG | → | RIFG | 0.44 | LACC | → | Input | 0.11 | RIPL | → | RIFG | 0.11 |
| RACC | → | LACC | 0.44 | LIFG | → | LACC | 0.11 | Input | → | LOCC | 0.11 |
| RIPL | → | LIPL | 0.44 | LIPL | → | LACC | 0.11 | Input | → | LACC | 0.11 |
| LOCC | → | Input | 0.22 | LIPL | → | RACC | 0.11 | Input | → | RACC | 0.11 |
| LACC | → | LIFG | 0.22 | ROCC | → | RIPL | 0.11 |  |  |  |  |

**Table C28:** Two-Step(Alasso): rhyming task data edges ordered by frequency of appearance (individual datasets).

|  |  |  |  |  |  |  |  |  |  |  |  |
| --- | --- | --- | --- | --- | --- | --- | --- | --- | --- | --- | --- |
| ROCC | → | LOCC | 0.89 | LACC | → | LIPL | 0.33 | LOCC | → | RIPL | 0.11 |
| LOCC | → | ROCC | 0.78 | LIFG | → | LACC | 0.33 | LOCC | → | Input | 0.11 |
| LACC | → | RACC | 0.78 | LIPL | → | LACC | 0.33 | LACC | → | RIPL | 0.11 |
| LIFG | → | RIFG | 0.78 | LIPL | → | RIFG | 0.33 | LIFG | → | ROCC | 0.11 |
| Input | → | LOCC | 0.78 | RIFG | → | RIPL | 0.33 | LIFG | → | RIPL | 0.11 |
| LOCC | → | LIPL | 0.67 | Input | → | LIFG | 0.33 | LIFG | → | Input | 0.11 |
| LIFG | → | LIPL | 0.67 | Input | → | RIFG | 0.33 | LIPL | → | LOCC | 0.11 |
| LIPL | → | RIPL | 0.67 | LIFG | → | LOCC | 0.22 | LIPL | → | RACC | 0.11 |
| RACC | → | RIPL | 0.56 | LIFG | → | RACC | 0.22 | ROCC | → | LACC | 0.11 |
| RIFG | → | LIFG | 0.56 | ROCC | → | RACC | 0.22 | ROCC | → | LIFG | 0.11 |
| RIPL | → | RIFG | 0.56 | ROCC | → | RIPL | 0.22 | RACC | → | LOCC | 0.11 |
| LACC | → | LIFG | 0.44 | RACC | → | LACC | 0.22 | RIFG | → | LACC | 0.11 |
| LIPL | → | LIFG | 0.44 | RACC | → | LIFG | 0.22 | RIFG | → | LIPL | 0.11 |
| ROCC | → | RIFG | 0.44 | RACC | → | ROCC | 0.22 | RIPL | → | LOCC | 0.11 |
| RACC | → | RIFG | 0.44 | RIFG | → | ROCC | 0.22 | RIPL | → | LACC | 0.11 |
| RIFG | → | RACC | 0.44 | RIPL | → | ROCC | 0.22 | RIPL | → | RACC | 0.11 |
| RIPL | → | LIPL | 0.44 | Input | → | LACC | 0.22 | Input | → | RACC | 0.11 |
| LOCC | → | LACC | 0.33 | Input | → | LIPL | 0.22 |  |  |  |  |
| LOCC | → | LIFG | 0.33 | Input | → | ROCC | 0.22 |  |  |  |  |

**Table C29:** FASK: rhyming task data 2-cycles ordered by frequency of appearance (individual datasets).

|  |  |  |  |
| --- | --- | --- | --- |
| LOCC | ↔ | ROCC | 0.56 |
| LACC | ↔ | RACC | 0.33 |
| LIPL | ↔ | RIPL | 0.33 |
| LIFG | ↔ | RIFG | 0.11 |
| LIPL | ↔ | RACC | 0.11 |

**Table C30:** Two-Step(Alasso): rhyming task data 2-cycles ordered by frequency of appearance (individual datasets).

|  |  |  |  |
| --- | --- | --- | --- |
| LOCC | ⇌ | ROCC | 0.67 |
| LIFG | ⇌ | RIFG | 0.44 |
| LIFG | ⇌ | LIPL | 0.22 |
| LOCC | ⇌ | LIFG | 0.11 |
| LACC | ⇌ | LIPL | 0.11 |
| LIFG | ⇌ | ROCC | 0.11 |
| LIFG | ⇌ | RACC | 0.11 |
| LIPL | ⇌ | RIPL | 0.11 |
| RACC | ⇌ | RIFG | 0.11 |
| RACC | ⇌ | RIPL | 0.11 |
| RIFG | ⇌ | RIPL | 0.11 |

### C6 Measurement noise and sampling resolution results

Values used to compute percentage changes reported in Figures 4 and 5 for data with and without measurement noise (abbreviated as m.n) and a fast sampling rate TR = 1.2 seconds, and a slower of TR = 3 seconds. The table shows average results over 60 repetitions of ten datasets concatenated from Network 4, for precision and recall of adjacencies (AP, AR), orientations (OP, OR) and 2-cycle detection (2CP, 2CR), and running times in seconds. Corresponding standard deviations are presented next.

**Table C31:** Precision and recall average results over 60 repetitions of 10 datasets concatenated, for Network 4 under different conditions of sampling resolution (1.2s and 3s TR) and measurement noise(m.n or without m.n).

| **FASK (average)** |  |  |  |  |  |  |  |
| --- | --- | --- | --- | --- | --- | --- | --- |
| Network 4 | AP | AR | OP | OR | 2CP | 2CR | r.time(s) |
| BOLD (m.n. // 1.2s TR) | 0.97 | 0.85 | 0.97 | 0.88 | 1.00 | 1.00 | 0.24 |
| BOLD (without m.n // 1.20s TR) | 0.98 | 0.86 | 0.98 | 0.88 | 1.00 | 0.94 | 0.19 |
| BOLD (m.n // 3s TR) | 0.94 | 0.80 | 0.95 | 0.84 | 1.00 | 1.00 | 0.26 |

| **Two-Step(Alasso) (average)** |  |  |  |  |  |  |  |
| --- | --- | --- | --- | --- | --- | --- | --- |
| Network 4 | AP | AR | OP | OR | 2CP | 2CR | r.time(s) |
| BOLD (m.n. // 1.2s TR) | 0.92 | 0.80 | 0.73 | 0.71 | 0.64 | 0.86 | 64.53 |
| BOLD (without m.n // 1.20s TR) | 0.99 | 0.91 | 0.99 | 0.93 | 0.97 | 1.00 | 17.52 |
| BOLD (m.n // 3s TR) | 0.90 | 0.80 | 0.76 | 0.75 | 0.68 | 0.88 | 76.68 |

| **MVGC (average)** |  |  |  |  |  |  |  |
| --- | --- | --- | --- | --- | --- | --- | --- |
| Network 4 | AP | AR | OP | OR | 2CP | 2CR | r.time(s) |
| BOLD (m.n. // 1.2s TR) | 0.70 | 0.95 | 0.52 | 0.90 | 0.33 | 1.00 | 0.65 |
| BOLD (without m.n // 1.20s TR) | 0.53 | 0.67 | 0.50 | 0.56 | 0.66 | 0.43 | 0.71 |
| BOLD (m.n // 3s TR) | 0.91 | 0.51 | 0.87 | 0.51 | 0.99 | 0.68 | 0.29 |

| **GlobalMIT(FAS) (average)** |  |  |  |  |  |  |  |
| --- | --- | --- | --- | --- | --- | --- | --- |
| Network 4 | AP | AR | OP | OR | 2CP | 2CR | r.time(s) |
| BOLD (m.n. // 1.2s TR) | 0.99 | 0.46 | 0.89 | 0.50 | 0.97 | 0.93 | 4.66 |
| BOLD (without m.n // 1.20s TR) | 1.00 | 0.38 | 0.94 | 0.42 | 0.97 | 0.63 | 2.91 |
| BOLD (m.n // 3s TR) | 1.00 | 0.18 | 0.99 | 0.14 | NaN | 0.00 | 3.20 |

**Table C32:** Precision and recall standard deviation over 60 repetitions of 10 datasets concatenated, for Network 4 under different conditions of sampling resolution (1.2s and 3s TR) and measurement noise(m.n or without m.n).

| **FASK (standard deviation)** |  |  |  |  |  |  |  |
| --- | --- | --- | --- | --- | --- | --- | --- |
| Network 4 | AP | AR | OP | OR | 2CP | 2CR | r.time(s) |
| BOLD (m.n. // 1.2s TR) | 0.04 | 0.06 | 0.04 | 0.05 | 0.03 | 0.00 | 0.15 |
| BOLD (without m.n // 1.20s TR) | 0.04 | 0.06 | 0.03 | 0.05 | 0.03 | 0.12 | 0.18 |
| BOLD (m.n // 3s TR) | 0.06 | 0.06 | 0.04 | 0.04 | 0.00 | 0.00 | 0.16 |

| **Two-Step(Alasso) (standard deviation)** |  |  |  |  |  |  |  |
| --- | --- | --- | --- | --- | --- | --- | --- |
| Network 4 | AP | AR | OP | OR | 2CP | 2CR | r.time(s) |
| BOLD (m.n. // 1.2s TR) | 0.06 | 0.08 | 0.09 | 0.10 | 0.14 | 0.15 | 7.77 |
| BOLD (without m.n // 1.20s TR) | 0.02 | 0.04 | 0.04 | 0.04 | 0.07 | 0.00 | 8.05 |
| BOLD (m.n // 3s TR) | 0.07 | 0.07 | 0.08 | 0.07 | 0.15 | 0.16 | 14.76 |

| **MVGC (standard deviation)** |  |  |  |  |  |  |  |
| --- | --- | --- | --- | --- | --- | --- | --- |
| Network 4 | AP | AR | OP | OR | 2CP | 2CR | r.time(s) |
| BOLD (m.n. // 1.2s TR) | 0.06 | 0.05 | 0.04 | 0.06 | 0.04 | 0.00 | 0.07 |
| BOLD (without m.n // 1.20s TR) | 0.11 | 0.13 | 0.13 | 0.12 | 0.32 | 0.23 | 0.09 |
| BOLD (m.n // 3s TR) | 0.11 | 0.12 | 0.08 | 0.09 | 0.05 | 0.21 | 0.05 |

| **GlobalMIT(FAS) (standard deviation)** |  |  |  |  |  |  |  |
| --- | --- | --- | --- | --- | --- | --- | --- |
| Network 4 | AP | AR | OP | OR | 2CP | 2CR | r.time(s) |
| BOLD (m.n. // 1.2s TR) | 0.03 | 0.07 | 0.08 | 0.05 | 0.08 | 0.12 | 1.92 |
| BOLD (without m.n // 1.20s TR) | 0.02 | 0.06 | 0.06 | 0.06 | 0.10 | 0.22 | 1.07 |
| BOLD (m.n // 3s TR) | 0.02 | 0.08 | 0.05 | 0.07 | NaN | 0.00 | 1.62 |

